# The evolution of high-order genome architecture revealed from 1,000 species

**DOI:** 10.1101/2025.07.05.663309

**Authors:** Yizhuo Che, Stephen J. Bush, Hui Lin, Xiaofei Yang, Qi Xie, Yuchun Liu, Deyu Meng, Kai Ye

## Abstract

Spatial genome organization plays a crucial regulatory role, but its evolution remains unknown. Leveraging Hi-C data from 1,025 species, we trace the evolutionary trajectories of 3D genome, through two higher-order architectures, ‘global folding’ (spatial organization of karyotype) and ‘checkerboard’ (chromatin compartments). Early life forms, including prokaryotes and unicellular eukaryotes, mostly display random configurations. Through the evolution of plants, global folding became the prominent architecture with eudicots and remained dominant. Animals, however, progressively developed stronger checkerboard throughout vertebrate and invertebrate evolution, and even during early embryogenesis, suggesting a conserved gene co-regulation mechanism. In contrast, plants prefer linear gene clusters over checkerboard. Both strategies of gene arrangement reinforce the biological principle ‘structure determines function’: divergent evolutionary paths converge on architectural solutions to meet the greater regulatory demands.

## INTRODUCTION

Among the regulatory mechanisms, three-dimensional (3D) genome structures have become a key area of interest. The advent of genome-wide chromatin conformation capture (Hi-C) techniques has revolutionized our ability to study how 3D genome folding influences the fine-tuning of gene regulation^1,2^. Hi-C measures physical proximity between pairs of sites across the genome, resulting in a ‘contact map’ of interactions, linking gene loci with regulatory elements. These structures are now understood to play a pivotal role in gene expression^3,4^. However, despite these advances, the role of 3D genome folding in evolution remains poorly understood, largely due to the scarcity of datasets, which are typically restricted to closely-related species^5–10^, confining previous studies to narrow evolutionary lineages.

To address this limitation, we leverage Hi-C data from thousands of species (mainly from the Darwin Tree of Life project11) to more fully explore the evolutionary significance of 3D genome. By doing so, we aim to extend the basic principle established by the discovery of the double helix – that structure determines function – to higher levels of genome organization. By algorithmically restoring long-range chromatin interactions, we identified two higher-order genome architectures at the whole-genome scale, which we refer to as ‘global folding’ and ‘checkerboard’, which vary across evolutionary time. By exploring 3D genome organization across the entire tree of life, this study provides new insights into how living systems evolve intricate structures.

## RESULTS

### Two whole genome-scale genome architectures: global folding and checkerboard

To investigate the large-scale genome architectures across the tree of life, we analyzed Hi-C data from 1,025 species, spanning bacteria, fungi, plants, and animals (Fig. 1; Data. 1), representing around 3,800 million years of evolutionary history. The vast phylogenetic diversity of these species and their enormous variation in genome size (from 2.1 Mb in *Thermococcus kodakarensis* to 14.6 Gb in *Triticum aestivum*; Fig. S1) make it impossible to identify conserved synteny blocks, which are the foundation for cross-species comparison of traditional localized genome structures such as TADs and loops^5,8^. To address this limitation, we developed a two-step pre-processing approach aimed at providing a broader perspective rather than a focus on local sequences. First, we automatically scaled chromosomal maps to a comparable size. Second, we devised NormDis, a **Norm**alization algorithm to remove **Dis**tance-dependent biases, which re-weights and makes visible long-range chromatin interactions, which in Hi-C data are invariably obscured by the far more prevalent short-range contacts (see Methods and Supplementary Methods; examples shown in Fig. S2). After applying NormDis and examining the normalized Hi-C maps, it became apparent that across evolutionary history two major whole-genome-scale architectures consistently occurred. We refer to these as ‘checkerboard’ and ‘global folding’; we schematically illustrate these terms in Fig. S3 and define them below.

**Fig. 1.**
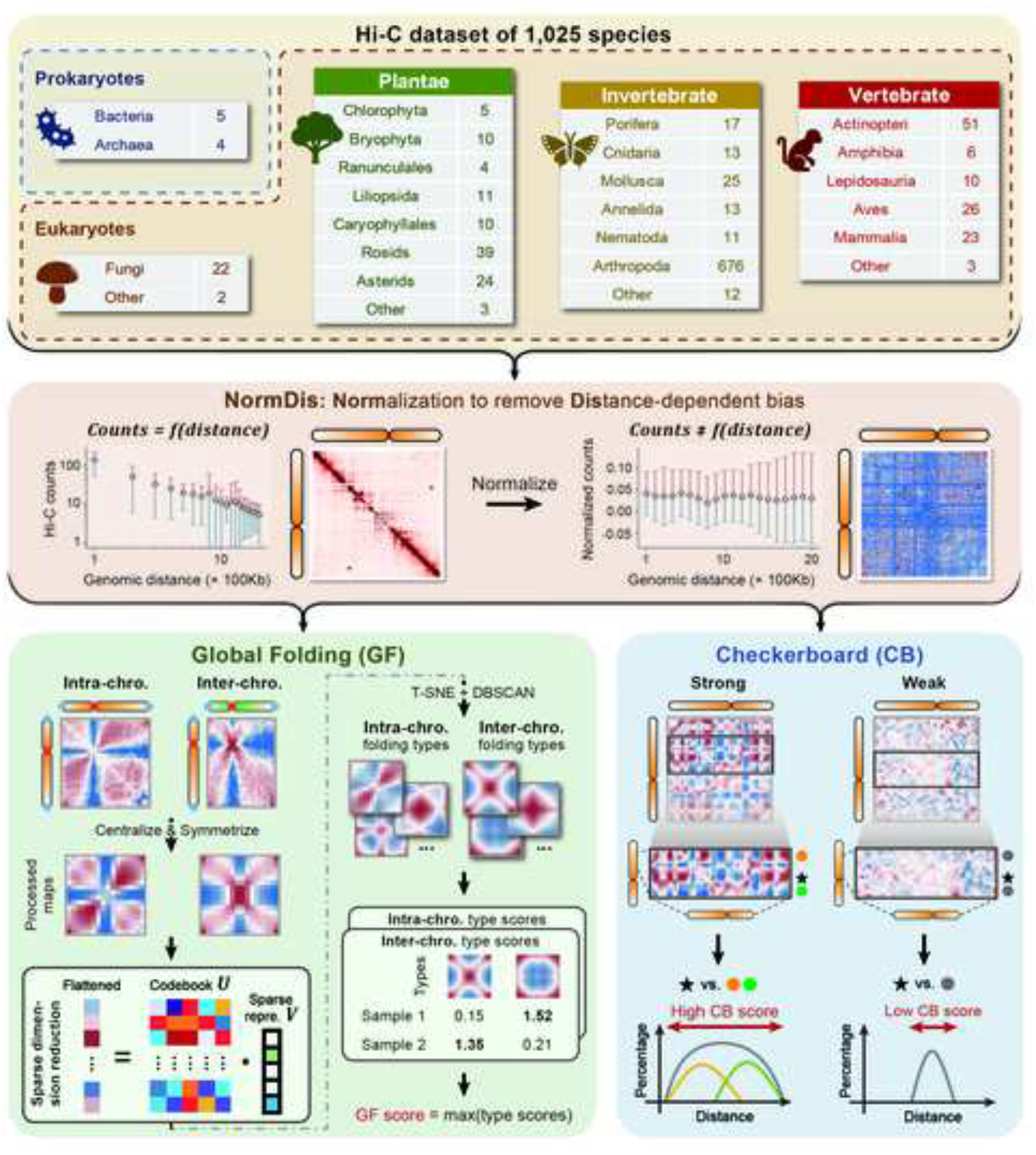
Analytical framework for deciphering higher-order genome architectures. The top panel displays the taxonomic distribution of 1,025 species, spanning prokaryotes (archaea and bacteria), fungi, plants and animals (invertebrates and vertebrates). Computational pipeline for high-order genome architectures comprises three modules: 1) NormDis, which re-weights Hi-C intearactions to remove distance-dependent biases to reveal preferred interactions (marked in red) and repelled interactions (marked in blue). 2) global folding quantification: normalized maps are pre-processed through centralization and symmetrization, and then major global folding types were discovered via unsupervised clustering; 3) checkerboard strength, which is quantified by the information entropy of interaction similarities. Higher entropy reflects stronger segregation of active and inactive compartments. Details are described in Methods.

Global folding describes how an entire set of chromosomes are spatially organized within the nucleus, with the folding of each chromosome typically (but, as discussed below, not always) anchored by three points: two at the ends (telomeres) and one in between, which strongly co-localizes with centromeres (Fig. S4). For example, in yeast (*Saccharomyces cerevisiae*), X-shaped patterns on the normalized map indicate a polarized conformation, in which center and end anchors cluster separately (Fig. S3A). However, we also observed other global folding patterns beyond ‘X’ shapes. To systematically classify these patterns, we employed a multi-step computational framework on both intra- and inter-chromosomal maps (Fig. 1; see Methods). First, we applied centralization and symmetrization to enhance topological clarity. Next, we performed feature extraction on the chromosomal maps based on sparse principal component analysis^12^, to eliminate noises and uncommon features. Finally, we categorized five intra- and five inter-global folding types through t-SNE^13^and DBSCAN^14^(Fig. 2A and S3). The global folding score (GFS) was assessed by the similarities between input normalized maps and identified folding types (documented in Data. 2 and 3).

**Fig. 2.**
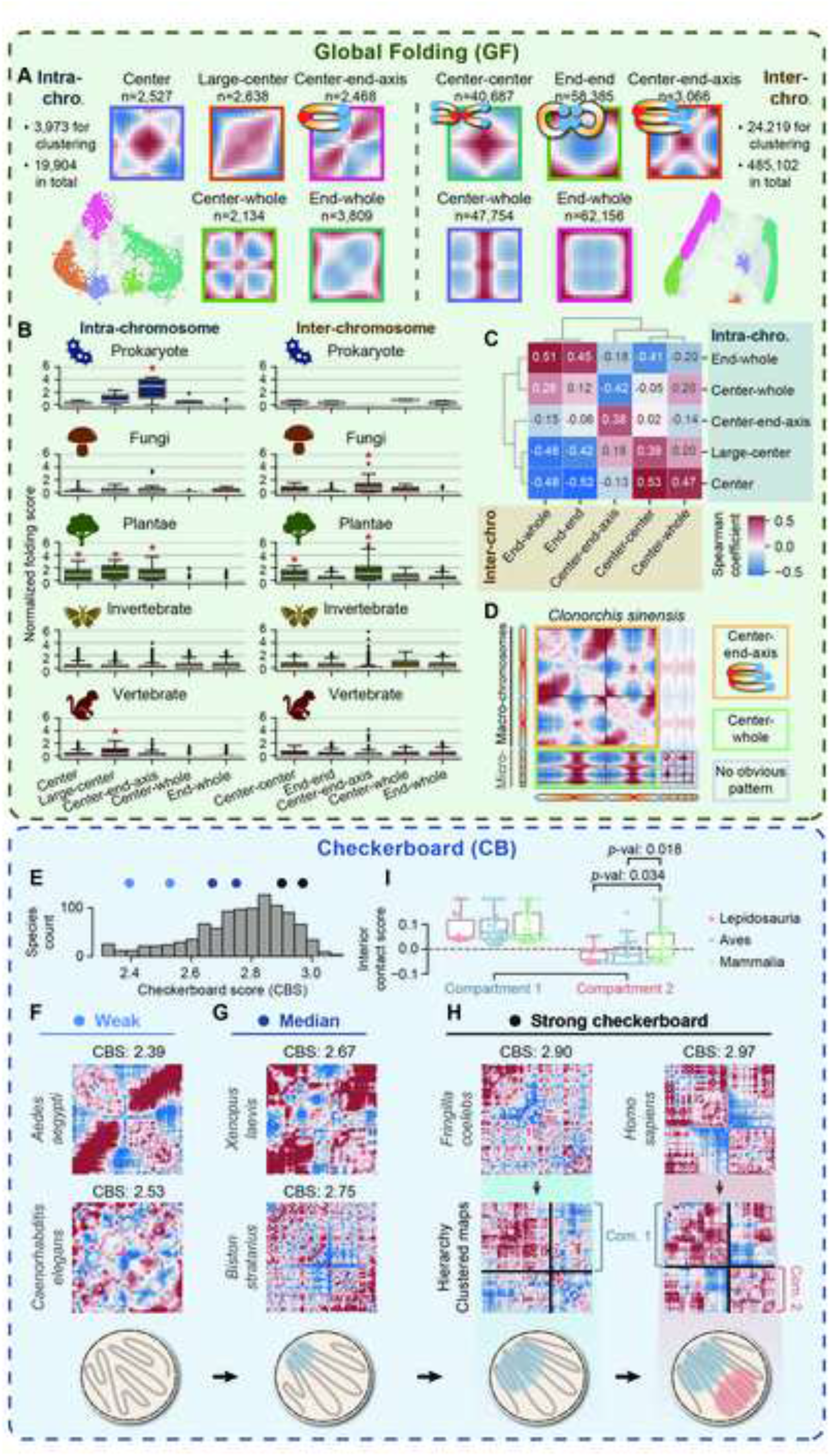
Diversity of global folding (A-D) and checkerboard patterns (E-I) across species. (A) Major global folding types. Five Intra- and five inter-chromosomal global folding types were identified through unsupervised clustering, shown by t-SNE projection. Previously characterized structures are annotated with schematics, while unlabeled ones represent novel foldings. The distribution of folding types is represented by their chromosomal map counts across all analyzed samples, reflecting their biological prevalence. (B) Enrichment of global folding across major taxonomic groups. Global folding scores for intra- and inter-chromosomal patterns across major taxonomic groups: prokaryotes (intra: n=9; inter: n=2), fungi (n=22 for both), plants (n=107), invertebrates (n=766), and vertebrates (n=126). Red asterisks (^*^) denote significant enrichment (one-tailed Wilcoxon signed-rank test, p < 0.05). (C) Interplay between intra- and inter-chromosomal folding types. Spearman correlation coefficients between folding types were calculated to quantify the association between distinct folding types. Coefficients were computed using global folding scores derived from 1,025 species. (D) Global folding polymorphism. Representative normalized map of Clonorchis sinensis demonstrates the coexistence of multiple global folding types within a single genome. (E) Distribution of checkerboard scores across all 1,025 species. Colored points pinpoint the checkerboard score for representative species. (F) Example species lacking checkerboard pattern. These species often exhibit global folding or randomized interaction patterns, reflecting relatively homogeneous interaction networks. (G) Intermediate species with mixed checkerboard and global folding/random regions. (H) Checkerboard organization. The second row exhibits hierarchically clustered contact maps, with black lines marking the segregation of compartments. The accompanying illustrations depict the spatial compartments, with different colors representing different compartments. (I) Interior contact scores of compartments, indicating their compaction levels. Positive values indicate condensed chromatin states, while negative values reflect dispersed domains.

Checkerboard patterns are the regular crisscrossed textures on normalized maps, which reflect their spatial compartmentalization (Fig. S3B). In human^2^and mice^15^, these patterns correspond to chromatin compartments, which separate euchromatic from heterochromatic regions. However, we found that checkerboard ‘strength’ varies widely across species, with some showing pronounced contrast between the different compartments comprising the checkerboard; others, by contrast, showed weaker or negligible contrast. To quantify checkerboard strength, we developed an entropy metric based on chromatin interaction polarity (Fig. 1; see Methods). Since genomic regions from the same compartment tightly interact^2^, their spatial interactions should more closely resemble each other, whereas regions from different compartments tend to repel, resulting in a lower degree of similarity. The degree of this interaction polarization, measured by information entropy, serves as a robust indicator of checkerboard ‘strength’; in effect, its degree of modularity (documented in Data. 4).

These two higher-order genome architectures, global folding and checkerboard, are remarkably conserved across mature tissues from same species (Fig. S5). This is quantitatively supported by smaller differences among tissues of given species (average standard deviation (std) of four tested species, checkerboard score (CBS): 0.06, intra-global folding score (GFS): 0.57, inter-GFS: 0.27; detailed scores are documented in Data. 5) than inter-species comparisons (std of CBS: 0.17, intra-GFS: 0.80, inter-GFS: 0.90), making them ideal for cross-species comparisons.

### Intra- and inter-chromosomal global folding

After analyzing 19,904 intra- and 485,102 inter-chromosomal maps from 1,025 species, we identified five major intra-(Center, Large-Center, Center-End-Axis, Center-Whole, and End-Whole) and five inter-chromosomal (Center-Center, End-End, Center-End-Axis, Center-Whole, and End-Whole) global folding patterns (Fig. 2A, examples in Fig. S3A). Each pattern is widely distributed, supported by a minimum of 2,134 chromosomal maps. Some of them, as labeled by the illustrations in Fig. 2A, align with previously reported structures^16^, while others represent novel architectures. For example, the newly discovered center-whole pattern is characterized by vertical or horizontal interaction signals anchored at the center, while end-whole is marked by signals at the ends. Although the biophysical architectures underlying these patterns remain unclear, they reveal the existence of structures beyond those previously identified. Other architectures may also emerge as more species data become available.

The prevalent global folding type varies among taxonomic kingdoms (Fig. 2B; examples on Fig. S6). Animals (both invertebrates and vertebrates) generally exhibit weaker global folding, with no specific type significantly enriched (minimum Wilcoxon signed-rank test *p*-value: 0.39, except for large-center pattern in vertebrates *p*-value: 0.023, *p*-values are documented in Data. 6). This indicates a relatively disordered organization at the whole-genome scale. In contrast, plants, prokaryotes (intra-chromosomal maps), and fungi (inter-chromosomal maps), strongly favor a center-end-axis architecture (maximum *p*-value: 0.031; Fig. 2B; Data. 6), which reflects a conformation in which the centromeres and telomeres cluster separately, while chromosome arms are arranged in parallel.

Despite the prevalent global folding, exceptions exist within each kingdom. For example, *Bufo bufo*, the common toad, exhibits a center-end-axis pattern, an architecture otherwise rare in vertebrates (Fig. S6). Nevertheless, closely related species tend to share similar global folding structures, as indicated by the significantly smaller differences across smaller taxonomic divisions (for example orders) than larger ones (kingdoms; mean intra-chromosomal GFS difference: 0.872 for orders and 1.195 for kingdoms, Mann-Whitney U test *p*-value: 0.005; mean inter-chromosomal GFS difference: 0.992 and 1.196, respectively, *p*-value: 0.032; Fig. S7; detailed differences are documented in Data. 7). This suggests that lineage-specific events may have shaped global folding patterns which were then stably inherited. An example is the emergence of condensin II, which resulted in the loss of global folding architectures in multiple organisms^16^. Notably, global folding appears independent of chromosomal morphology, as we found no significant correlations between GFS and chromosome length (Pearson correlation coefficient (PCC) < 0.13; Fig. S8).

We next investigate the interplay between intra- and inter-chromosomal global folding within same species. Usually, the same intra- and inter-chromosomal global folding type co-occur (Fig. 2C; average Spearman correlation coefficient, SCC: 0.40). Nevertheless, cross-type pairs are also observed, particularly those that share the same anchor points (average SCC: 0.22; examples on Fig. S9A). For example, the intra-chromosomal “center” pattern (a central chromatin domain), is often accompanied by the inter-chromosomal “center-whole”, where the center anchor interacts with an entire chromosome (SCC: 0.47; examples in Fig. S9B). These observations suggest that the anchor points play a central role in shaping global folding.

Within a single species, different chromosomes could also adopt distinct global folding patterns, a phenomenon we term global folding polymorphism. For instance, in *Clonorchis sinensis*, the first two long chromosomes display center-end-axis architectures, whereas the remaining five short ones lack large-scale folding structures (Fig. 2D). Polymorphic folding is more common in species with irregularly smaller chromosomes, referred to as micro-chromosomes in prior studies^17^(SCC between polymorphic score and chromosome length difference ratio: 0.29, *p*-value < 10^−16^; Fig. S11A). Reptiles (Lepidosauria) and birds (Aves), in which micro-chromosomes are commonly abundant^17^, exhibit the highest polymorphism scores (reptiles: 1.228; birds: 1.203; Fig. S11D and Data. 8), indicating that micro-chromosomes might contribute to the diverse folding patterns within species.

### Checkerboard score reflects the degree of genome compartmentalization

The checkerboard score (CBS) quantifies the intensity of checkered textures on normalized maps, serving as a proxy for chromatin compartmentalization. Among 1,025 species, CBS values range from weak (∼2.3) to strong (∼3) (Fig. 2E; documented in Data. 4). As with global folding patterns, checkerboard patterns do not correlate with chromosome length (PCC with length: -0.087; Fig. S12). This suggests that checkerboard organization is a biologically regulated architecture rather than a byproduct of chromosome morphology.

Species with low CBS typically exhibit alternative chromatin organizations as either strong global folding or randomized chromatin interactions. For example, *Aedes aegypti* (CBS: 2.39; Fig. 2F) displays a strong center-end-axis global folding pattern, which may restrict long-range chromatin interactions and thereby hinder the formation of checkerboard. Maps of *Caenorhabditis elegans* (CBS: 2.53) lack organized compartmentalization and instead show random spot-like interactions, probably resulting from the unrestricted thermal collisions of chromatin. Additionally, some species exhibit intermediate, transitioning chromatin architectures. For instance, the African clawed frog, *Xenopus laevis* (CBS: 2.67), and the Oak Beauty moth, *Biston stratarius* (CBS: 2.75) show mixed Hi-C patterns, where certain loci display checkerboard patterns, while others exhibit global folding or random interactions (Fig. 2G). This suggests that genomes may undergo a gradual evolutionary shift, transitioning from weaker to stronger checkerboard patterns.

Checkerboard patterns emerge from selective interactions between genomic regions, leading to a well-organized chromatin structure (Fig. 2H). For example, in human (CBS: 2.97), the genome is divided into two compartments, where loci within the same compartment interact tightly, while those in different compartments repel^2^. Interestingly, some species with slightly lower checkerboard scores, such as the chaffinch, *Fringilla coelebs* (CBS: 2.90), display an alternative “one compartment” architecture (Fig. 2H), in which a single compartment dominates, while the other regions remain less structured. The “one compartment” architecture appears more prevalent in reptiles (Lepidosauria) and birds (Aves) than in mammals (Mammalia) (Mann Whitney U test *p*-value of reptiles: 0.034; birds: 0.018; Fig. 2I). This suggests that in these species, genomes might be capable of assembling only a single chromatin hub, reflecting an intermediate evolutionary stage preceding the well-established A/B compartments.

### Evolution of genome architectures along the tree of life

To investigate the evolutionary dynamics of genome architectures, we analyzed 21 taxonomic groups across prokaryotes, fungi, plants and animals (Fig. 3). Each group includes Hi-C data from at least four species, ensuring statistical robustness and minimizing the influence of outliers (numbers shown in Fig. 1A).

**Fig. 3.**
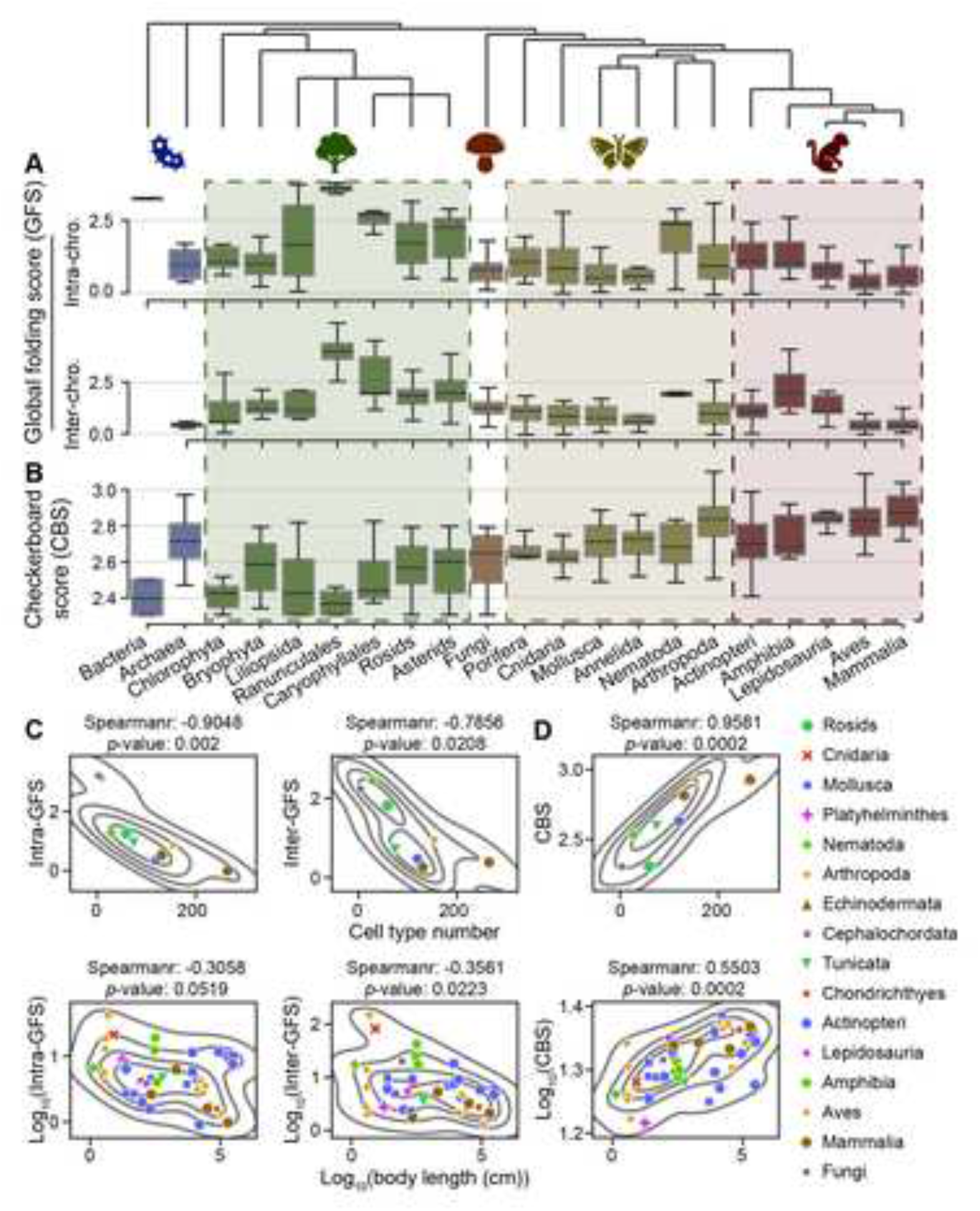
Evolutionary trends of high-order architectures and their correlations to species complexity. (A) Intra- and inter-chromosomal global folding scores (GFS) across 21 phylogenetically ordered taxonomic groups (group sizes shown in Fig. 1A). (B) Checkerboard scores (CBS) for the same taxonomic groups. (C) Global folding scores show negative correlation with species complexity proxies: cell type number (CTN; n=8) and body length (n=26). (D) Checkerboard scores significantly correlate with species complexity proxies: cell type number (CTN; n=8, p=0.0002) and body length (n=26, p=0.0002), underscoring their role in facilitating biological complexity.

Global folding strength does not follow a consistent evolutionary trend (Fig. 3A; detailed scores are documented in Data. 9). In plants, both intra- and inter-chromosomal global folding scores (GFSs) peak at Ranunculales, a basal eudicot lineage (median intra-GFS 3.60 vs. 0.95 for all species; inter: 4.02 vs. 1.04). After this peak, GFSs gradually decline. In vertebrates, GFSs reach a lower peak at amphibians (Amphibia; intra GFS: 0.99; inter: 2.02), decreasing further both in birds (Aves; 0.29; 0.38) and mammals (Mammalia; 0.54; 0.38). Among invertebrates, only roundworms (Nematoda; 2.38; 1.93) exhibit strong GFSs, while others lack pronounced global folding patterns. Overall, global folding strength fluctuates without a clear evolutionary trend, suggesting that it is not shaped by a universal evolutionary force.

We next examined whether GFS correlates with organismal complexity, using a direct measure of complexity, cell type number (CTN)^18,19^as well as an indirect one, body length^20^(a proxy for long-term effective population size, itself a strong correlate of complexity; values are documented in Data. 10). We observed a negative correlation between GFS and CTN (SCC of intra-GFS: -0.90, *p*-value: 0.02; inter: -0.78, *p*-value: 0.02), and also a weaker but still negative correlation with body length (intra: -0.36, *p*-value: 0.002; inter: -0.31, *p*-value: 0.051; Fig. 3C), suggesting that species with higher complexity tend not to be associated with strong global folding patterns.

In contrast to GFS, checkerboard score exhibits a clear evolutionary trend. In both invertebrate and vertebrate animals, as well as in eudicots, CBS steadily increases over evolutionary time (Fig. 3B; documented in Data. 11). In addition, CBS is strongly and significantly associated with organism complexity as evidenced by the strong correlations with CTN (SCC: 0.95; *p*-value: 0.0002) and body length (SCC: 0.55; *p*-value: 0.0002; Fig. 3D). As stronger checkerboard patterns reflect greater levels of chromatin compartmentalization, these findings are consistent with the hypothesis that gene expression becomes more precisely coordinated in species with greater complexity.

### Distinct evolutionary trajectories of genome architectures

To explore the evolutionary trajectory of genome architectures, we traced both global folding and checkerboard patterns of extant species and inferred their ancestral states. By mapping species onto a two-dimensional matrix based on global folding and checkerboard score, we identified a spectrum of genome organizations, ranging from disordered maps (e.g., *Sphagnum tenellum*), strong global folding (e.g., *Papaver bracteatum*), strong checkerboard (e.g., *Homo sapiens*), and mixed architectures (e.g., *Verbascum thapsus*) (Fig. 4A). Generally, GFS negatively correlates with CBS (intra-chromosomal SCC: - 0.26, *p*-value < 10^−16^; inter SCC: -0.34, *p*-value < 10^−16^, shown in Fig. S13). This supports the hypothesis that global folding physically obstructs checkerboard formation, likely due to conflicted spatial constraints.

**Fig. 4.**
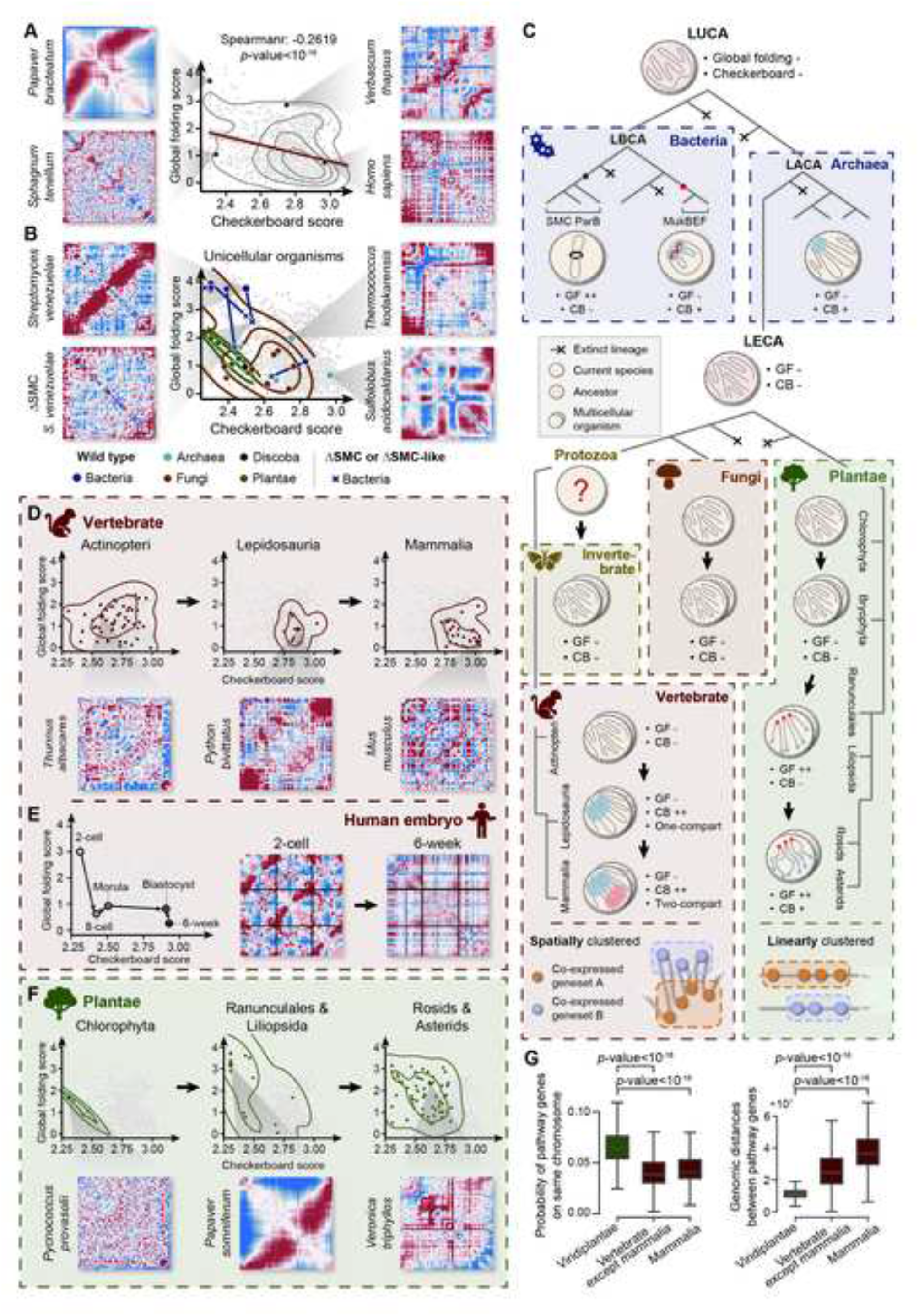
Evolutionary and developmental trajectories of 3D genome architecture. (A) Negative correlation between checkerboard score and intra-chromosomal global folding score across 1,025 species. The correlation with inter-global folding score is shown in Fig. S13. Example species illustrate the spectrum of architectures: Sphagnum tenellum (random), Papaver bracteatum (strong global folding), Homo sapiens (strong checkerboard), and Verbascum thapsus (mixed architectures). (B) Unicellular organisms (including bacteria, archaea, algae, fungi, and Discoba) mostly exhibit random architecture without both architectures. Left: normalized maps of representative bacteria and its mutant (S. venezuelae) show that SMC knockout disrupts global folding. Right: example archaea with coarsen checkerboard patterns. (C) Evolutionary trajectory of 3D genome architectures. Schematic of genome architecture evolution from the disordered structures of common ancestors to diverse architectures of extant species: bacteria mostly adopt global folding; archaea have developed rudimentary checkerboard; each lineage of eukaryotes has forged its own paths, where animals exhibit checkerboard patterns along with spatial gene clustering, and plants displays global folding patterns with linear gene clustering. (D) Evolution of vertebrate genome architecture. Progressive strengthening of checkerboard architectures is observed in vertebrates. Representative groups are shown with exemplified maps: Actinopteri (weak checkerboard), Lepidosauria (intermediate checkerboard with one compartment) and Mammalia (strong checkerboard with two compartments). Other groups are shown in Fig. S14 and S15. (E) Genome architectural transition during early human embryogenesis: strong global folding of 2-cell stage to checkboard patterns of 6-week embryo. (F) Evolution of plant genome architecture. Divergent patterns appear in major plant clades: Chlorophyta (random map), Ranunculales & Liliopsida (dominant global folding) and Rosids & Asterids (persistent global folding with weak checkerboard). (G) Lineage-specific genomic arrangements of pathway genes. Pathway genes of plants (n=784 pathways) adopt linear clustering as indicated by the higher probability of locating on the same chromosome and shorter intra-chromosomal distance. Genes of vertebrates (n=2087 for vertebrates except Mammalia, n=2463 for Mammalia) exhibit a more scattered distribution. Statistical significance was assessed using Mann-Whitney U tests.

To infer the genome architecture of the last eukaryotic common ancestor (LECA), we analyzed 14 unicellular eukaryotes (including five algae and eight fungi) and also 17 earliest multicellular animal groups, Porifera. Most of these early life forms exhibit random chromatin interaction maps (30 of 31), lacking strong global folding or checkerboard patterns (median GFS: 0.79, median CBS: 2.50 for unicellular eukaryotes, Fig. 4B; median GFS: 1.07, median CBS: 2.64 for Porifera, Fig. S14 and S15). Taken together, these observations suggest that LECA possessed a disordered genome structure with weak global folding and weak checkerboard patterns (Fig. 4C).

In bacteria and archaea, genome architectures vary. Bacteria either exhibit strong global folding (4 of 5 analyzed bacteria; average GFS: 3.26; Fig. 4B), or a partial checkerboard (*Escherichia coli*; CBS: 2.83; Fig. S16). These divergent architectures are diminished or disrupted by the knockout of structural proteins (see Supplementary File), indicating that the last bacterial common ancestor (LBCA), which lacked such proteins, had an unstructured genome architecture, similar to that in knockout bacteria (Fig. 4C). Archaea lack notable global folding patterns (average GFS: 1.00), but exhibit a spectrum of checkerboard patterns from weak to strong (Fig. 4B). As with bacteria, we conjecture that the last archaeal common ancestor (LACA) also lacked strong genome architectures. Notably, the checkerboard patterns in archaea are coarser than in mammals, with larger interaction domains (median size: 4.96% of chromosome length vs. 3.17% of mice, Mann Whitney U test *p*-value: 0.0009, and 3.14% of human, *p*-value: 0.0007; Fig. S17). Similarly, we inferred the genome architecture of the last universal common ancestor (LUCA), from which the fundamental prokaryotic domains (archaea and bacteria) diverged^21^, to lack both global folding or checkerboard architectures (Fig. 4C).

From the random architectures of the common ancestors, animals and plants have evolved distinct spatial organizations. In both vertebrates and invertebrates, checkerboard patterns, indicative of functional compartmentalization, appear to increase over evolutionary time (Fig. 4D; Fig. S14 and S15). The trajectory begins with the disordered structures in fishes, transitions through “one-hub” architectures in reptiles and birds, and culminates at the A/B compartments in mammals. This may reflect an increased capacity to spatially cluster (and thereby coordinate the activity of) genes otherwise scattered throughout the genome.

Interestingly, we observed a similarly progressive emergence of checkerboard patterns during human early embryo development. Chromatin undergoes a disruption of global folding and a stepwise reconstruction of checkerboard architectures from the 2-cell stage (intra-GFS: 3.01, inter-GFS: 3.45, CBS: 2.31) to 6-week embryo (intra-GFS: 0.24, inter-GFS: 1.04, CBS: 2.93) (Fig. 4E; Fig. S18). The construction of checkerboard mirrors the evolutionary trend in animals, which suggests that the checkerboard pattern is a fundamental architectural feature required for gene regulation, enabling the emergence of increasing complexity.

Unlike animals, plants exhibit a markedly different evolutionary trajectory. During plant evolution, global folding surges dramatically, around the divergence of Liliopsida and Ranunculales (Fig. 4F). While checkerboard patterns increase modestly in eudicots (Fig. 4F, S14 and S15), they remain significantly weaker than in animals (median CBS of Rosids: 2.56, Mann Whitney U test *p*-value 2.5 × 10^−11^; Asterids: 2.60, *p*-value 2.4 × 10^−9^ compared to mammals: 2.88; Fig. S19). The dominance of global folding may physically affect checkerboard formation, constraining the long-range spatial compartmentalization of plant genomes, as seen in animals.

Despite their weaker checkerboard patterns, plants still require coordinated gene expression, particularly for their complex secondary metabolic pathways^22^. Instead of spatial clustering functionally associated genes, it is plausible that plants adopt an alternative strategy. For example, in opium poppy, *Papaver somniferum*, genes involved in the noscapine and morphine biosynthesis pathways are linearly adjacent on the genome, forming a localized gene cluster^23^. Consistent with this being a more widespread phenomenon, we found that genes within the same pathways are more proximal in plants than in vertebrates, as evidenced by a significantly higher probability of being on the same chromosome (median probability: 6.3% in plants compared to 3.9% in vertebrates besides mammals and 4.1% in mammals; Mann Whitney U test *p*-value < 10^−16^) and closer genomic distances (median distance between genes on same chromosome: 1.1 × 10^7^bp in plants compared to 2.4 × 10^7^bp in vertebrates and 3.6 × 10^7^bp in mammals; *p*-value < 10^−16^; Fig. 4G). This supports the hypothesis that in contrast to the spatial compartmentalization of genomes observed in animals, plant genomes have a greater degree of linear gene clustering.

## DISCUSSION

### Evolution of higher-order genome architectures

Our analysis of Hi-C data across 1,025 species suggests general principles governing 3D genome evolution, that originating from the random genome architectures of common ancestors, two organized, low-entropy genome architectures evolve, which we refer to as global folding and checkerboard. In prokaryotes, the majority of bacteria (4 of 5) exhibit global folding, whereas a majority of archaea (3 of 4) display coarse checkerboard patterns. This dichotomy extends to eukaryotes: plants evolved pronounced global folding, while invertebrate and vertebrate animals progressively strengthened checkerboard organization. The sustained organization of these architectures, energetically unfavorable yet prevalent, implies strong selective forces and specialized regulatory maintenance.

### Protein complexity facilitates architectural sophistication

The evolution of genome architectures is paralleled with increasing complexity of structure-maintenance machinery. In bacteria, the center-end-axis global folding pattern is regulated by a single SMC protein, suggesting its relatively simple regulatory mechanisms. In contrast, global folding in *Arabidopsis thaliana* is controlled by a more complex, two-step regulation involving multiple proteins^24^. Similarly, species exhibiting the strongest checkerboard patterns demonstrate advanced mechanisms, such as the specialized architectural proteins in *Drosophila* species^25,26^and the specific N-terminal of CTCF in mammals^26,27^. These observations refute the notion of genome architectures as passive byproducts, instead positioning them as actively engineered systems shaped by specialized protein-mediated mechanisms.

### Global folding and cellular plasticity

Although global folding lacks a correlation with species complexity, its association with cellular plasticity offers a compelling functional hypothesis. Plants, which exhibit the strongest global folding, retain widespread cellular totipotency^28^, and human 2-cell stage embryos (highly plastic stem cells) similarly display evident global folding. This parallels with prokaryotic systems, where disrupting global folding patterns (e.g., via SMC knock out in *B. subtilis*^29^or *C. crescentus*^30^) impairs chromosome segregation and cell cycle progression, as also seen in yeast mutants^31^. Taken together, we propose that global folding may serve as a scaffold for maintaining cellular pluripotency, with potential implications for regenerative medicine and developmental biology.

### Checkerboard and cellular specialization

The modular organization of functionally related genes is crucial for coordinated regulation, serving as a basis for cellular specialization. Two common mechanisms for achieving modularity are observed: linear genomic clustering or spatial clustering via checkerboard architectures. Linear clustering is particularly prevalent in species with random or global folding patterns, which lack fine-tuned spatial compartmentalization, as exemplified by operons in prokaryotes^32^and gene clusters in plants^23^. This arrangement may enable more rapid and dynamic response to fluctuating external stimuli, such as unpredictable weather, with less time and energetic expense; they would not need to be spatially coordinated first. This could be particularly advantageous for sessile organisms like plants^33^. However, linear clustering does impose evolutionary constraints, favoring genetic innovation through polyploidization, despite its rarity and associated risks of genome instability^34^.

In contrast, checkerboard architecture offers a spatially mediated mechanism, which we found to positively correlate with species complexity. By spatially organizing scattered genes into close proximity, checkerboard provides greater flexibility in gene regulation, which, in turn, facilitates the increasingly diverse cell types. Nevertheless, the transition towards checkerboard accompanies the loss of pluripotency. Interestingly, besides animals, we also observed weak but notable checkerboard-like organization in some eudicots, albeit constrained by their dominant global folding architecture (Fig. 4F, illustrated in Fig. 5). This indicates that besides linear clustering, some eudicots have also evolved a capacity to spatially cluster co-expressed genes. For example, studies of *Arabidopsis thaliana* have shown that co-regulated genes can be brought together through 3D topologies, even over long chromosomal distance^35^. These findings indicate that, relative to global folding, checkerboard structures are a more flexible genome architecture, the use of which over evolution have facilitated the increases of organism complexity.

**Fig. 5.**
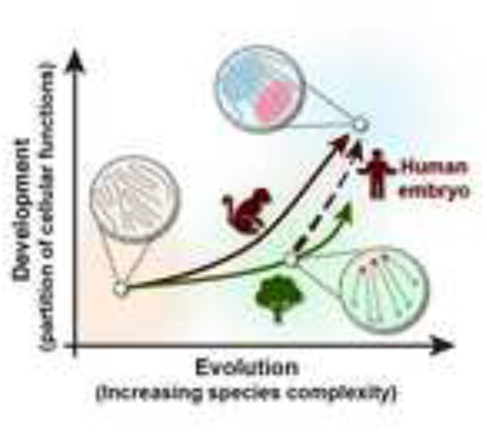
Trade-offs between two high-order genome architectures during evolution or development. The horizontal axis represents species complexity at maturity—the maximum attainable complexity of an organism. The vertical axis reflects developmental complexity, which is the degree of functional partitioning during ontogeny. While evolutionary and developmental complexity are inherently related and are not suitable for being represented as independent axes, we employ this simplified framework to illustrate the trajectories of 3D genome architecture during evolution and development. Animal evolution (solid red curve) exhibits progressive reliance on checkerboard architecture, driving the increase of species complexity but reducing developmental plasticity. Plant evolution (solid green curve) shows sustained global folding, maintaining cellular totipotency. Human embryogenesis (dashed red curve) recapitulates evolutionary transitions of genome architectures: early embryos prioritize global folding (2-cell stage), followed by checkerboard emergence (8-cell stage to 6-week embryo). The distinct evolutionary and developmental trajectories reflect a universal trade-off between genome architectures: checkerboard enables functional specialization at the cost of plasticity, while global folding preserves adaptability while limiting regulatory sophistication.

### Trade-off between plasticity and specialization

The evolution of species complexity is a testament to the ingenuity of life. The distinct trajectories of 3D genome organization underscore the potential trade-offs between global folding (associated with cellular plasticity) and checkerboard organization (linked to specialization), through which each evolutionary lineage or embryonic developmental trajectory has forged its own path (Fig. 5).

Overall, our findings expand the axiom “structure determines function” to high-order genome architectures, which shape high-level biological processes, such as cellular specialization or plasticity. This bridges evolutionary biology and developmental biology, offering insights for regenerative medicine (e.g., enhancing cellular plasticity) and synthetic biology (e.g., engineering gene regulation networks). Future studies dissecting the interplay between these architectures in diverse lineages will further unravel how life’s complexity is encoded in the 3D foldings of the genome.

## RESOURCE AVAILABILITY

### Lead contact

Requests for further information and resources should be directed to and will be fulfilled by the lead contact, Kai Ye.

### Materials availability

This study did not generate new unique reagents.

### Data and code availability

- This paper analyzes existing, publicly available data, accessible at Darwin Tree of Life project or NCBI. Detailed sources for each species are listed in Supplementary File 1.
- All original code has been deposited at https://github.com/xjtu-omics/HiArch and is publicly available at [DOI] as of the date of publication.
- Any additional information required to reanalyze the data reported in this paper is available from the lead contact upon request.

## ACKNOWLEDGMENTS

We thank the Darwin Tree of Life project consortium for sharing the Hi-C data used in this study.

## AUTHOR CONTRIBUTIONS

Conceptualization: K.Y. and Y.C. Methodology: Y.C. Investigation: Y.C. and Y.L. Formal Analysis: Y.C. Funding Acquisition: K.Y. Writing – Original Draft: Y.C. Writing – Review & Editing: K.Y., S.B., X.Y. Supervision: K.Y.

## SUPPLEMENTAL INFORMATION

A brief list (index) of the <u>supplemental files</u> you are including should appear in the main-text manuscript under a “supplemental information” heading. The files themselves should be uploaded separately, not as a part of this manuscript.

The primary file is typically a PDF, called “Document S1,” and should be listed first. Include a short list of the PDF’s contents, e.g., “Figures S1–S6” (see below), but **do not include their titles or legends here**; for supplemental figures, the titles and any legends should appear only in the supplemental PDF (but for papers publishing in the journal *Cell*, see the **NOTE** below). Additional non-PDF supplemental files (e.g., Excel files, video files, large datasets in ZIP formats) should have their titles and any legends listed here on their own lines.

Example (for journals other than *Cell*):

**Document S1. Figures S1–S6, Tables S1 and S2, and supplemental references Table S3. Traces of Edman sequencing of rhBMPs, related to Figure 2 Video S1. Stages of cell mitosis, related to Figure 3**

Mitosis is the process by which a cell divides to make two genetically identical cells.

In the example above, the author has uploaded three supplemental files: the primary supplemental PDF (Document S1), one Excel file, and one video. The PDF (Document S1) contains six supplemental figures; the figures as well as their full titles and legends are in the PDF. Also in this PDF are two supplemental tables, which can appear easily in PDF format, and additional references for works cited within the same PDF. One additional table has been uploaded separately as an Excel file. Finally, the author has uploaded a video file, for which a brief legend is included.

**NOTE:** For the journal *Cell* only, supplemental figures are treated just like main figures, with the image files uploaded separately and individually and the titles and legends appearing in this main text file, after the titles and legends for the main figures (see “figure titles and legends” section below). The supplemental figures will be included as part of the main text. If a *Cell* paper has supplemental figures but no other supplemental items, then there is no “Document S1” PDF, and there is no supplemental information index. Otherwise, the index might look like this example:

**Document S1. Tables S1–S5 and supplemental references Table S6. Traces of Edman sequencing of rhBMPs, related to Figure 2**

In the *Cell*-only example above, the author has uploaded two supplemental files: a primary supplemental PDF (Document S1), which contains five supplemental tables and a list of supplemental references, and one larger table as an Excel file. Because they are publishing in *Cell*, any supplemental figures will have been uploaded separately as individual files, and the titles and legends for those supplemental figures will be included in their main-text manuscript.

## STAR★METHODS

## KEY RESOURCES TABLE

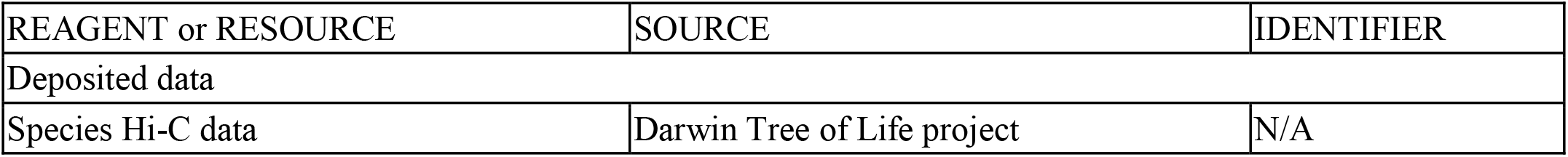

| REAGENT or RESOURCE | SOURCE | IDENTIFIER |
| --- | --- | --- |
| Deposited data |  |  |
| Species Hi-C data | Darwin Tree of Life project | N/A |

| REAGENT or RESOURCE | SOURCE | IDENTIFIER |
| --- | --- | --- |
| Deposited data |  |  |
| Species Hi-C data | Darwin Tree of Life project | N/A |

## EXPERIMENTAL MODEL AND STUDY PARTICIPANT DETAILS

### METHOD DETAILS

#### Pre-processing of Hi-C matrix

A Hi-C dataset comprising 1,517 species was collected. The tree of life was downloaded from NCBI Taxonomy. Species that could not be found in the taxonomy dataset were excluded, resulting in a total of 1,475 species. The sources and detailed information, including species names and NCBI species indices, are listed in Supplementary Table 1.

Hi-C maps are capable of reflecting the quality of genome assembly (examples shown in Figure S20). We manually checked the assembly quality and removed 450 species and 710 chromosomes with low-quality assembles (documented in Supplementary Table 1). At last, 1,025 species with high-quality assembles are kept.

To normalize the distinct genomic lengths among species, Hi-C maps are automatically scaled based on two criteria: 1) to make the number of bins per chromosome approximately the same, and 2) to ensure sufficient contacts at both intra- and inter-chromosomal scales. Notably, chromosomes with extremely sparse contacts, which typically result from the absence of certain chromosomes (e.g., sex chromosomes), are removed.

#### NormDis: Normalization to remove proximity effects

The NormDis algorithm is applied to eliminate distance-dependent biases on Hi-C maps, which is introduced by the polymeric nature of DNA molecules. The main idea is to make Hi-C contacts distance-independent by normalizing contacts by expected contact number,

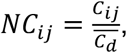

where *C*_*ij*_is the raw Hi-C contact of row *i* and column *j, NC*_*ij*_is the normalized contact, 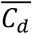 is the expected contact of distance *d* = |*i* − j|.

Previous studies showed that Hi-C contacts associate with genomic distances in a log-log correlation^36^. Based on this, we tried to calculate 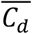 by linear fitting of logarithms of genomic distance and Hi-C contacts (Figure S21), calculated by:

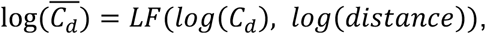

where *LF* represents the linear fitting optimized by the ordinary least squares. However, this linear correlation is observed only in a narrow interval of genomic distances (e.g., 100 kb to 10 Mb in *Acropora millepora*, as shown in Figure S21). Beyond this interval, deviations emerge, as shown at the distal regions of the *A. millepora* contact map. These deviations disrupt linear fitting accuracy, generating regional overestimations and underestimations that collectively impair normalization precision.

Diagonal mean values provide more accurate expected values for normalization,

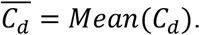

However, near the matrix corners, the shorter diagonal length increases sensitivity to outliers (Figure S21), often resulting in normalization errors.

Consequently, we developed a scaled diagonal method that aggregates long-distance diagonals to ensure computational robustness while preserving accuracy, by

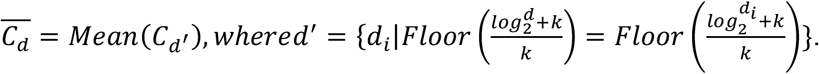

where *k* controls the number of diagonals to be aggregated. Notably, based on the linear nature of chromosome, expected contacts should be theoretically decreasing monotonically with genomic distances. However, some species, for example *Acropora millepora*, exhibit anomalously elevated long-range contacts, likely reflecting its unique global folding patterns (Figure S21). To preserve these signals, we enforce monotonic decay in the expected contact values.

Long-distance contacts are exposed after NormDis. To test whether these contacts reflect real physical structures within nuclei, we conducted a DNA FISH experiment, labeling centromeres and telomeres of opium poppy (*Papaver somniferum*; Figure S2B). Normalized maps of poppy display a special pattern, including the anti-diagonal lines on intra-maps and X-shape on inter-maps. These patterns indicate that centromeres and telomeres of long chromosome arms independently cluster at the two side of nuclei, while the short-arm telomeres freely float in between. This structure is evident by the reconstructed structures and the DNA FISH images, which indicates the consistence of normalized maps and real physical structures.

#### Detection and measurement of global folding Preprocessing for global folding

To unveil the global folding patterns from the normalized maps, a preprocessing pipeline is applied (Figure S22A; detailed in Supplementary Methods). Notably, maps sharing the same global folding type may exhibit distinct visual patterns, due to variations in the genomic positioning of their center anchors. It is exemplified in *Papaver bracteatum* (Figure S22A), where two separate maps from different chromosomes nevertheless preserve the same anti-diagonal center-end-axis pattern. To address this, we developed CenterFinder, a tool to identify center anchors (Figure S22B). It offers two modes tailored to different types of global folding:

1. The ‘Inter row sum’ mode identifies center anchors by detecting local maxima in inter-chromosomal contact row sums, making it optimal for anchors exhibiting prominent inter-chromosomal interactions.
2. The “Intra X” mode identifies center anchors by locating the anti-diagonal midpoints, specifically designed for intra-chromosomal center-end-axis patterns.

Typically, anchor with the strongest signal is designated as the default center anchor, but it can be manually adjusted. The mode used and manually re-chosen anchors are documented in Supplementary Table 12 and 13.

#### Global folding pattern mining

Global folding pattern mining extracts biologically representative patterns from high-noise maps via robust statistical analysis (Figure S22C). We initially filtered for strong global folding maps by selecting the top 5% of inter- and top 20% of intra-chromosomal maps exhibiting minimal dissimilarity between average-pooled and symmetrized maps, effectively excluding maps lacking strong folding patterns.

After the above pre-processes and selection, maps still contain considerable noises, which will hinder subsequent pattern mining. To address this issue, we apply sparse representation techniques to reduce the dimension of each map, as well as to reduce noise. Specifically, we assume that each processed map can be approximately reconstructed as a linear combination of a small set of features. Formally, for the *i*-th sample, *x*_*i*_∈ *R*^256^, we assume that

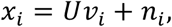

where *U* ∈ *R*^256×*k*^ denotes the dictionary matrix, shared across all maps, and randomly initiated; *k* is the element number of the dictionary; *v*_*i*_∈ *R*^*k*^ is a *k* dimensional representation of *x*_*i*_; and *n*_*i*_denotes the corresponding noise. It should be noted that *x*_*i*_represents distinct sparse patterns, thus it is reasonable to assume that *v*_*i*_can be low-dimensional (*i*.*e*., *k* is a small number) and sparse. We can simultaneously perform sparse representation on all maps, by solving the following sparse principal component analysis model:

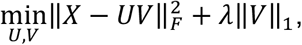

where *X* = [*x*_1_, *x*_2_, … *x*_*N*_] represents *N* input maps; *V* = [*v*_1_, *v*_2_, … *v*_*N*_] represents the sparse representation for all maps; and *λ* controls the trade-off between reconstruction accuracy and sparsity of *V*. Both the dictionary *U* and the sparse representation *V* will be jointly optimized using a bi-level iterative algorithm. After optimized sparse representation *V* is achieved, we can view *v*_*i*_as noise-reduced k-dimension representations of *x*_*i*_.

With noise-reduced *k*-dimension representations, we then apply t-SNE^13^to visualize and DBSCAN^14^to cluster. The average pattern of each cluster is designated as the representative pattern for each global folding types. In total, we identified five types for intra-chromosome and five for inter-chromosome (Figure 2A).

Although the dictionary matrix in sparse dimension reduction is randomly initiated, we re-run the pattern mining and still got the same types (Figure S23), indicating the robustness of our procedures.

We then compute the similarities of maps to identified types, serving as the global folding scores (GFS), by

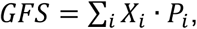

where *X*_*i*_is the *i*-th value from map, *P*_*i*_is the *i*-th value from type. Higher global folding score denotes higher alignment to folding type. GFSs for 1,025 species are documented in Supplementary Table 2 and 3.

#### Measurement of checkerboard strength

The checkerboard pattern, displayed as the crisscrossed textures on normalized maps, signifies the physical separation into genomic compartments. Checkerboard ‘strength’ reflects the degree of separation, characterized by two components: 1) the tight connections within the interior compartment, and 2) repelled contacts between different compartments. Accordingly, we measure the checkerboard strength by firstly computing the similarities between genomic bins to reveal the connections (high similarities) or repulsions (low similarities), and then measuring the strength of connections and repulsions (Figure S24).

The “cosine” distance metric is utilized for assessing the similarities,

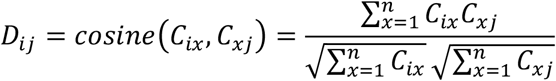

where *C*_*ij*_is the raw contact of bin *i* and bin *j, cosine* is the cosine distance, *D*_*ij*_is the distance between bin *i* and bin *j*. Lower distances indicate more similar connections, whereas higher distances reflect weak or repelled contacts between bins. Maps exhibiting pronounced checkerboard patterns show a bimodal distribution of similarity values (high and low; Figure S24), resulting in a high information entropy,

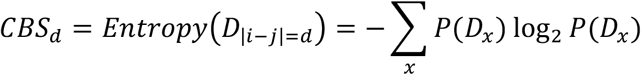

where *D*_|*i*−*j*|=*d*_ is the cosine distances with genomic distance *d*. In contrast, random maps exhibit intermediate similarities, reflecting neither preferential attraction nor repulsion between genomic bins, and result in lower entropies.

Even without a checkerboard (Figure S24), map with strong global folding can produce high entropy in whole-map calculations due to opposing short-distance and long-distance similarities. We addressed this by calculating distance-stratified entropies and averaging them into the checkerboard score,

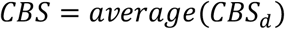

where *CBS*_*d*_are the distance-stratified entropies. Given the distance-dependence of this analysis, we omitted long-range interactions to avoid distortion from diminishing diagonal lengths. 15% chromosome length is set as the threshold for this long-distance removal. This threshold does not affect the entropy or subsequent results, as consistent results are obtained for thresholds of 10% or 20% (Figure S25). CBSs for 1,025 species are documented in Supplementary Table 4.

## Supplementary Methods

Dataset comprising Hi-C experiments of 1,517 species from various kingdoms is collected. The tree of life was downloaded from NCBI Taxonomy. Species that could not be found in taxonomy were excluded, resulting in a total of 1,475 species. The sources and detailed information, including species names and NCBI species indices, are listed in Supplementary Table 1.

### Pre-processing of Hi-C matrix

To scale chromosomal maps from diverse species to comparable sizes, Hi-C maps are automatically scaled by NormDis, which takes sparse matrix format as input. It first removes chromosomes with extremely low Hi-C interaction signals (empty signal percent > 99%), as well as short assembly contigs (distinguished from chromosomes by the naming prefixes). Next, a rough concatenation is performed to ensure the maximum chromosomal size not exceeding 600. The roughly concatenated matrix then undergoes iterative concatenation to ensure sufficient contacts (>95%) at whole-genome scale. If this threshold remains unsatisfied after maximum concatenation (10 iterations), the Hi-C data will be treated as low-coverage and excluded from analysis. Additionally, chromosomes with bin sizes <20 were excluded. This process ensures that only sufficiently large and well-covered chromosomal data are retained for further processing.

After normalization, poorly assembled chromosomes become readily apparent, as Hi-C maps are capable of reflecting assembling quality^45^. Assembly errors could produce false patterns that might be misinterpreted as real physical structure signals. For example, the duplication in the second chromosome of *Andrena bucephala* can be misinterpreted as the center-whole global folding (Figure S25). We manually checked the assembly quality and removed 450 species and 710 chromosomes with low-quality assembles. At last, 1,025 species with high-quality assembles are kept. Additionally, chromosomes were sorted in descending order of length, for better identification. This step ensures a cleaner and more structured dataset for downstream analysis.

### NormDis: Normalization to remove proximity effects

The NormDis algorithm is applied to eliminate distance-dependent biases on Hi-C maps, which is introduced by the polymeric nature of DNA molecules. The main idea is to make Hi-C contacts distance-independent by normalizing contacts by expected contact number,

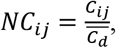

where *C*_*ij*_ is the raw Hi-C contact of row *i* and column *j, NC*_*ij*_ is the normalized contact,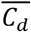is the expected contact of distance *d* = |*i* − *j*|.

Here, we developed a scaled diagonal method that aggregates long-distance diagonals to ensure computational robustness while preserving accuracy, by

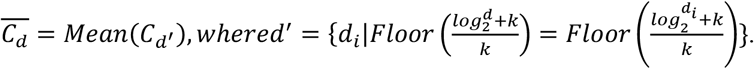

After normalization, wavelet denoising (from skimage.restoration with VisuShrink algorithm^46^and ‘db3’ wavelet function) is applied to maps to reduce noise. Subsequently, z-normalization is applied to reveal preferential or repelled contacts over background. Additionally, contact values exceeding ±3σ were truncated at the 3-sigma boundaries.

### Detection and measurement of global folding

#### Preprocessing for global folding

To unveil the global folding patterns in Hi-C maps, a four-step preprocessing pipeline is applied to each map, including spatial filtering, centralization, average pooling, and symmetrization (Figure S21A). Spatial filtering was initially applied to attenuate small-scale noise. In detail, a median filter (with kernel size = 10% map size) is applied with scipy.ndimage.uniform_filter. To mitigate extreme values, we performed winsorization by truncating the top and bottom 5% of interaction values.

Maps sharing the same global folding type may exhibit distinct visual patterns, due to variations in the genomic positioning of their center anchors. It is exemplified in *Papaver bracteatum* (Figure S22A), where two separate maps from different chromosomes nevertheless preserve the same anti-diagonal center-end-axis pattern. To address this, we developed CenterFinder, a tool to identify center anchors (Figure S22B). It offers two modes tailored to different types of global folding:

1. The ‘Inter row sum’ mode identifies center anchors by detecting local maxima in inter-chromosomal contact row sums, making it optimal for anchors with prominent inter-chromosomal interactions. In detail, the top 33.3% of interactions are initially selected from filtered maps, as anchor points often correspond to the high-interaction regions (Figure S21B). All maps then underwent z-normalization to normalize signal intensities across samples. Anchor positions were identified by detecting local maxima in row sums of resulting maps. To attenuate outliers, row sums underwent spatial convolution (scipy.ndimage.convolve with 10% chromosomal-length kernel, ‘reflect’ mode). This smoothing process enhances the robustness of anchor detection. However, not all local maxima correspond to true center anchors. For example, due to the inherent noise in Hi-C maps, subdominant maxima near the primary peak may be falsely detected as anchors. To address it, we eliminated the subdominant maxima, which were identified if the difference between its height and the lowest intervening point was less than 3% of the average value. Additionally, we removed maxima that located at the ends of chromosome, as these represent end anchors rather than center anchors. This was accomplished by excluding anchors that did not exhibit a decreasing signal toward the chromosome end. Finally, the remaining anchor with the highest value was automatically identified as the center anchor. This algorithm is not designed to identify the most accurate anchors, but rather to filter out incorrect candidates. Further manual verification is recommended for optimal results.
2. The ‘Intra X’ mode identifies center anchors by locating the anti-diagonal midpoint, specifically designed for center-end-axis patterns. We performed convolution on filtered maps using anti-diagonal sliding kernels to identify the anti-diagonal midpoints (Figure S21B). The local maxima in convolution output served as anchor candidates. Unlike the ‘inter row sum’ mode, the detected anti-diagonal pattern of ‘intra x’ mode is not common on Hi-C maps, thereby inherently avoiding noises. Therefore, the anchor filtering steps of the ‘inter row sum’ mode become unnecessary and are omitted. The peak with the highest value was automatically identified as the center anchor.

The center anchors of center-related types—center-end-axis and center-center for intra-chromosomal types, and center-center, center-end-axis, and center-whole for inter-chromosomal types—located near centromeres in all 10 centromere-annotated species (Figure S4). This observation highlights the biological relevance of centromeres as central points in chromosomal organization. Centromere positions are obtained from UCSC table browser and other studies^31,47^.

Notably, for some species without obvious global folding patterns, the “third” mode, ‘no anchor’ is available to bypass the detection of center anchors, thereby speeding up the computation. Importantly, omitting anchor detection does not affect the calculation of global folding strength, as their maps lack strong global folding patterns regardless of where the anchors are. For detailed information, the specific mode used and any manually re-selected anchors are documented in Supplementary Table 12 and 13.

We then applied average pooling using PyTorch’s torch.nn.AdaptiveAvgPool2d to downsample the maps to 16×16 matrices. If center anchors are provided, the pooling process ensures that anchors are positioned at center.

Considering the symmetrical organization of chromosomes from the center centromere to the two-sided telomeres, we symmetrize the maps accordingly. For intra-chromosomal maps, we average the raw maps with their centrally rotated counterparts to ensure the central rotational symmetry. For inter-chromosomal maps, we average the raw maps with those rotated by 90, 180, and 270 degrees to achieve comprehensive symmetry.

### Global folding pattern mining

After the above pre-processes and selection, maps still contain considerable noises, which will hinder subsequent pattern mining. To address this issue, we apply sparse representation techniques to reduce the dimension of each map, as well as to reduce noise. Specifically, we assume that each processed map can be approximately reconstructed as a linear combination of a small set of features. Formally, for the *i*-th sample, *x*_*i*_ ∈ *R*^256^, we assume that

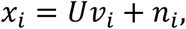

where *U* ∈ *R*^256×*k*^ denotes the dictionary matrix, shared across all maps, and randomly initiated; *k* is the element number of the dictionary; *v*_*i*_ ∈ *R*^*k*^ is a *k* dimensional representation of *x*_*i*_; and *n*_*i*_ denotes the corresponding noise. It should be noted that *x*_*i*_ represents distinct sparse patterns, thus it is reasonable to assume that *v*_*i*_ can be low-dimensional (*i*.*e*., *k* is a small number) and sparse. We can simultaneously perform sparse representation on all maps, by solving the following sparse principal component analysis model:

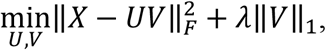

where *X* = [*x*_1_, *x*_2_, … *x*_*N*_] represents *N* input maps; *V* = [*v*_1_, *v*_2_, … *v*_*N*_] represents the sparse representation for all maps; and *λ* controls the trade-off between reconstruction accuracy and sparsity of *V*.

It is worth noting that the above formulation directly imposes sparsity on *V*, which may potentially lead to suboptimal solutions. Therefore, we introduce an auxiliary variable *W* to serve as a sparse representation of *V*. The inclusion of *W* decouples the sparsity constraint from the reconstruction term, enabling explicit sparsity control via an *l*_1_-norm regularization while promoting structured and interpretable representations. Then to solve the proposed optimization problem, we employ a bi-level optimization strategy, i.e., to update each variable iteratively while keeping others fixed. The optimization process consists of three key steps: updating *V*, updating *W*, and updating U.

1. Updating *V*. Given fixed *U* and *W*, we solve the following subproblem for update *V*:

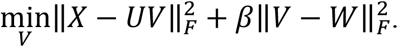 This is a least-squares problem with a closed-form solution:

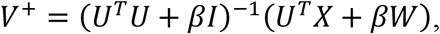

where *I* represents the identity matrix.
2. Updating *W*. Given *V*, the sparse representation *W* is updated by solving:

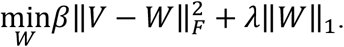 The optimization can be efficiently solved using the element-wise soft-thresholding function:

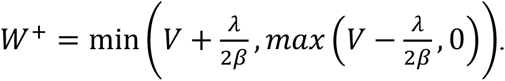
3. Updating *U*. Finally, the dictionary *U* can be updated by solving:

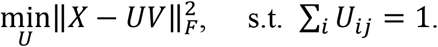 The closed-form solution is given by:

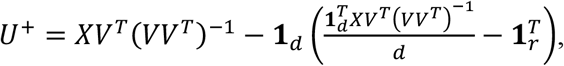

where **1**_*d*_ and **1**_*r*_ are all-ones vectors. The above updates are performed iteratively:

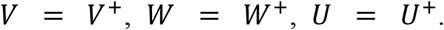

The process continues until convergence, which is typically determined by monitoring changes in *V*, e.g. when ‖ *V*^+^ − V‖ falls below a predefined threshold. By jointly optimizing *V, W* and *U*, we obtain a structured sparse representation that effectively captures the underlying patterns in *X*. The optimized *w*_*i*_ provides a denoised, low-dimensional representation, that can be viewed as noise-reduced k-dimension representations of *x*_*i*_.

Notably, transposition of inter-chromosomal maps (swapping chromosomal positions between x and y axes) should theoretically preserve global folding patterns. However, due to their inherent asymmetries, different folding configurations may emerge. To resolve this, we implemented selective transposition, applying only when the transposed map better matched the current folding pattern. This approach ensures consistent sparse representations across inter-chromosomal maps.

With noise-reduced k-dimension representations, we then apply T-SNE^19^(perplexity=700 for intra and 4,000 for inter) to visualize and use DBSCAN^20^(eps=1.2, min_samples=130 for intra and 1, 620 for inter) for clustering the maps. The average pattern of each cluster is designated as the representative pattern for each global folding types. In total, we identified five types for intra-chromosome and five for inter-chromosome.

Although the dictionary matrix in sparse dimension reduction is randomly initiated, we re-run the pattern mining and still got the same patterns (Figure S22), indicating the robustness of our procedures.

### Calculation of global folding scores

We then compute the similarities of processed maps to patterns as the global folding scores (GFS), by

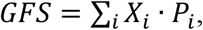

where *X*_*i*_ is the *i*-th value from map, *P*_*i*_ is the *i*-th value from pattern. Higher global folding score denotes higher alignment to pattern. GFSs for 1,025 species are documented in Supplementary Table 2 and 3.

Notably, the distributions of GFSs vary across different types due to inherent variations in type strength. To enable equitable comparisons between types, GFSs are z-normalized. The maximum normalized value among types is then set as the integral score for global folding strength.

We aim at avoiding setting an arbitrary threshold to determine the exact global folding types for each map, as this would result in the information loss regarding their strength. However, it is sometimes useful for describing the prevalence of types among species by quantifying how many species are included. To this end, we set the top 33.3% probability for standard normalization (0.43) as the threshold, solely for the purpose of numerical calculation, for example in Figure 2A and Figure S10.

### Measurement of checkerboard strength

The “cosine” distance metric is utilized for assessing the similarities,

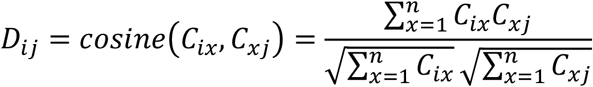

where *C*_*ij*_ is the raw contact of bin *i* and bin *j*, *cosine* is the cosine distance, *D*_*ij*_ is the distance between bin *i* and bin *j*. oower distances indicate more similar connections, whereas higher distances reflect weak or repelled contacts between bins. Maps exhibiting pronounced checkerboard patterns show a bimodal distribution of similarity values (high and low; Figure S24), resulting in a high information entropy,

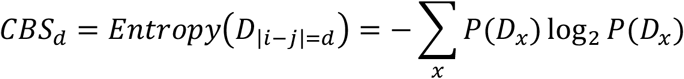

where *D*_|*i*−*j*|=*d*_ is the cosine distances with genomic distance *d*. To discretize the probability distribution for entropy calculation, we partition the distances into 30 uniformly spaced intervals over the range [0, 1.6], where each interval has equal width of 0.053 (1.6/30). In contrast, random maps exhibit intermediate similarities, reflecting neither preferential attraction nor repulsion between genomic bins, and result in lower entropies.

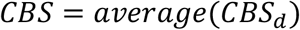

where *CBS*_*d*_ are the distance-stratified entropies. Given the distance-dependence of this analysis, we omitted long-range interactions to avoid distortion from diminishing diagonal lengths. 15% chromosome length is set as the threshold for this long-distance removal. This threshold does not affect the entropy or subsequent results, as consistent results are obtained for thresholds of 10% or 20% (Figure S25). CBSs for 1,025 species are documented in Supplementary Table 4.

Notably, maps with sizes smaller than 50 are excluded from this analysis, as their limited dimensions are insufficient to support the formation of complex crisscrossed checkerboard patterns. This ensures that only maps with adequate size are considered for meaningful analysis.

Our algorithm of checkerboard score is simple and interpretable as it is based on the physical structures of checkerboard. Still, complicated algorithms might be meaningful to get more accurate measurements. The checkerboard scores show a significant association with organism complexity, reflecting the evolving ability of spatially gathering co-expressed genes. However, species in this analysis are primarily metazoans, due to the limited data of organismal complexity. Further studies are needed to fully assess this correlation.

## Supplementary materials

### Construction of the tree of life and classification of species

To construct the tree of life, species names were first transferred to NCBI species index, documented in Supplementary Table 1. The phylogenetic tree was then generated using the ete3 Python toolkit’s built-in NCBI taxonomy database.

To analyze phylogenetic related species, we selected 21 representative taxonomic groups spanning plants, invertebrates, and vertebrates. Each group includes Hi-C data for at least four species to minimize the influence of outliers. These groups were systematically arranged according to their phylogenetic positions in the tree of life. For taxonomic groups occupying identical topological positions in the phylogeny, we ordered them according to established evolutionary conventions. For example, oiliopsida, a phylum from monocots, was placed before Ranunculales, a basal phylum of eudicots. Caryophyllales, another basal phylum of eudicots, was positioned before Rosids and Asterids. Orders between Rosids and Asterids is biologically uninformative as they are sister clades. This same phylogenetic principle applies to another sister group, Annelida and Mollusca. Nematoda was placed before Arthropoda, which is considered the most complex groups among invertebrates.

### Different tissues from same species

To examine the conservation of global folding and checkerboard across mature tissues, we collected poppy (*Papaver somniferum*) Hi-C data of stem, leaf and root from Xiaofei et al.^48^, *Arabidopsis thaliana* Hi-C data of flower, leaf, seedling from GSE155503, GSE201839 and GSE145769, human Hi-C data of H1-hESC, HFF and gm12878 from 4DN data portal, mouse Hi-C data of thymocyte, mature olfactory sensory neurons (OSNs) and B-cell from 4DN data portal. All calculations (NormDis, global folding, and checkerboard scores) employed default parameters, except for global folding detection modes. Specifically, we used the ‘intra-x’ mode for poppy, while applying the ‘no anchor’ mode for human and mouse.

### Interplay between intra- and inter-global folding types

To investigate the interplay between intra- and inter-chromosomal global folding, we performed the Spearson correlations between intra- and inter-global folding types scores on all 1,025 analyzed species, by scipy.stats.spearsonr with default parameters.

### Global folding polymorphism

Global folding polymorphism is a phenomenon where chromosomes from a same species adopt distinct global folding structures. To quantify the strength of this polymorphism, we developed an analysis pipeline to measure the degree of polymorphism, by using the standard deviation (STD) as the metric. Specifically, to reduce the impact of outliers, we removed outlier chromosomal maps with top and bottom 10% global folding scores. We then calculated the STDs for each folding type. The average STD was set as the polymorphic score for each species.

We prioritized inter-chromosomal maps over intra-maps in calculation, as the greater abundance of inter-maps offers more robust results. Additionally, species lacking strong global folding (with a strongest global folding score < 0.43, corresponding to the top 33.3% probability of standard distribution) will be assigned a polymorphic score of zero.

To investigate the correlation between polymorphism with chromosome length range, we conducted the Spearson correlation between polymorphic scores with chromosome length difference ratio, calculated by (*max*(*chrolength*) − *min*(*chrolength*))/*max*(*chrolength*).

### Assess the compaction degree of chromatin compartments through hierarchy clustering

To identify chromatin compartments, we employed hierarchical clustering to divide the genome into two compartments. Hierarchy clustering is an easier method with clearer vision, compared to current algorithms for compartments^7,49^. Nevertheless, it is only suitable for small-size maps, for example the scaled maps in our analysis, due to its complexity.

In detail, we calculated the “cosine” distances between genomic bins as the inputted distance maps to hierarchy clustering. It is conducted by scipy.cluster.hierarchy with method=“ward” and optimal clustering on. In our analysis, the hub with higher interior contacts is identified as the first hub.

### Correlation with organism complexity

We collected the species complexity proxies to analyze their correlations to high-order architectures. Morphological features, the body lengths, are collected from Bush *et al*.^3^. Cell type numbers are collected from Chen *et al*.^24^. The correlations are calculated through scipy.stats.spearmanr, with default parameters. We removed outliers by the 3-sigma rule, to avoid them influencing the correlation analysis.

### Global folding analysis for circular chromosomes

The sequential ends of circular chromosomes linearly connect to each other, resulting in the long-distance interactions at the upper right and lower left corners of intra-chromosomal maps. To eliminate their influence in global folding detection, we masked these two corners of circular chromosomes. Specifically, the normalized interaction values of points with Manhattan distances to the upper right and lower left corners less than 5 are set to zero. Circular chromosome is common in prokaryotes. Maps of all prokaryotes—except for *Streptomyces venezuelae*, which has linear chromosomes—are processed with this mask.

### Measure the checkerboard sizes

Besides the checkerboard strength, other aspects of checkerboard also exhibit species specificity. For example, the checkerboard sizes of *S. acidocaldarius* and mammals are obviously different (Figure S17). We developed a pipeline to measure the sizes of compartmental domains. It firstly converts the normalized maps to adjacency maps, by calculating the “cosine” distances between genomic bins. Compartmental domains are exposed by this conversion, given that domains are made up of bins with similar interactions. Next, we identified domains by merging nearby bins of low distances (cosine distance < 0.5). Small domains with size smaller than 3 are removed.

### Embryo development

We collected the human early embryo development Hi-C data from Chen *et al*.^50^. The NormDis, checkerboard, and global folding calculations were performed using default parameters, with the exception of adjusted center anchors. Specifically, we utilized the centromere locations obtained from the UCSC Table Browser as the center anchors.

### Proximity analysis of pathway genes

We assess the proximity between pathway genes, to support the hypothesis that genes are more likely to linearly cluster in plants than in animals. We collected the gene annotation from Gene Ontology (GO) dataset. Only pathways marked as biological processes (aspect is “P” in.goa file) are utilized for analysis. Genome annotations were downloaded from NCBI dataset. To avoid influenced by the outliers, only pathways involving more than 5 genes are analyzed. Meanwhile, to facilitate calculation, pathways with more than 500 genes are excluded.

### Significance and correlation analysis

Mann-whitney U test and Wilcoxon signed-rank test is conducted by mannwhitneyu and wilcoxon from scipy. Spearman correlation and Pearson correlation is conducted by spearmanr from scipy and corrcoef from numpy.

## Supplementary figures

**Figure S1.**
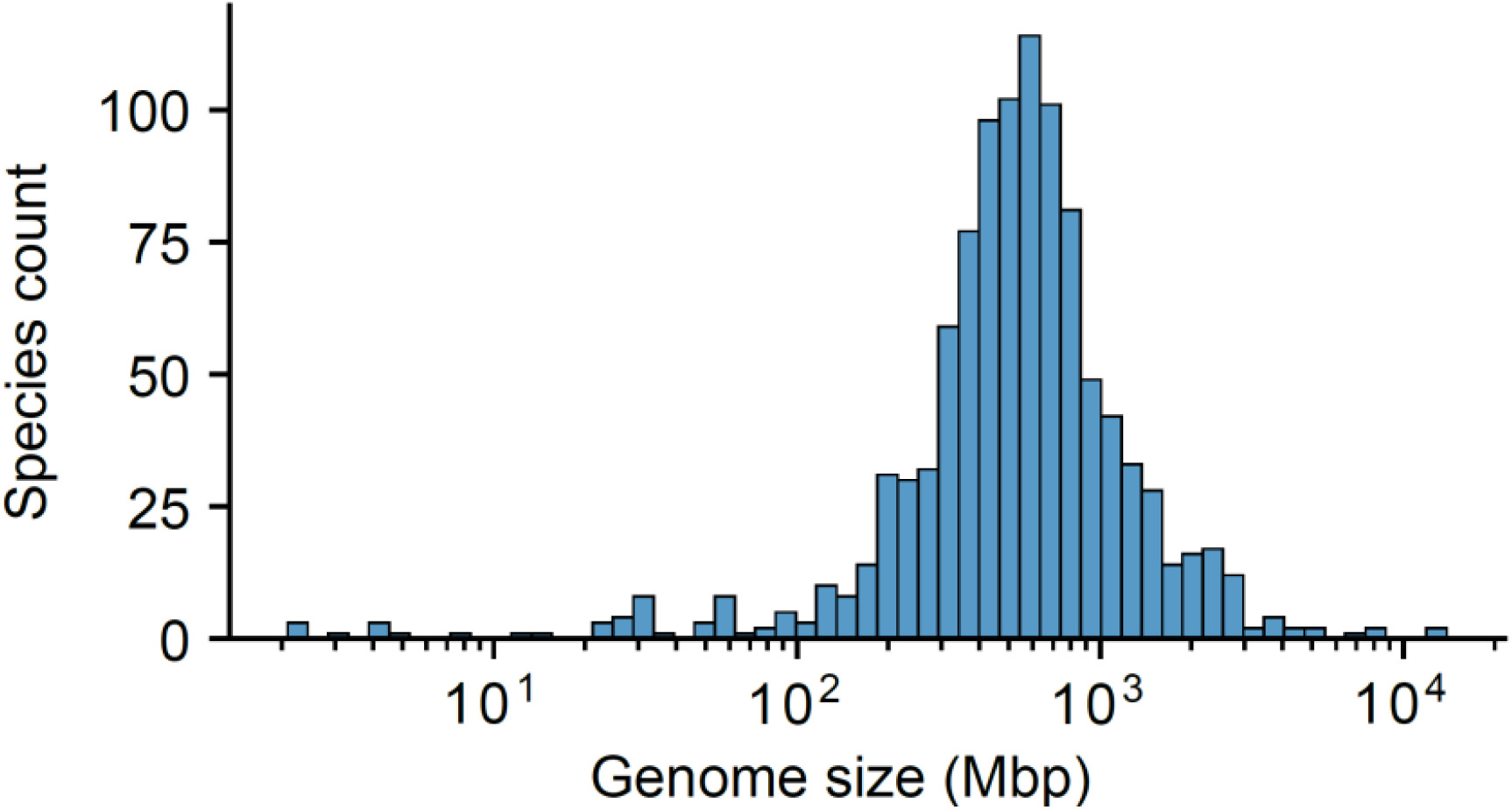
Variation in Genome Sizes. Genome sizes are enormously varied among analyzed 1,025 species, from 2.1 Mb in *Thermococcus kodakarensis* to 14.6 Gb in *Triticum aestivum*.

**Figure S2.**
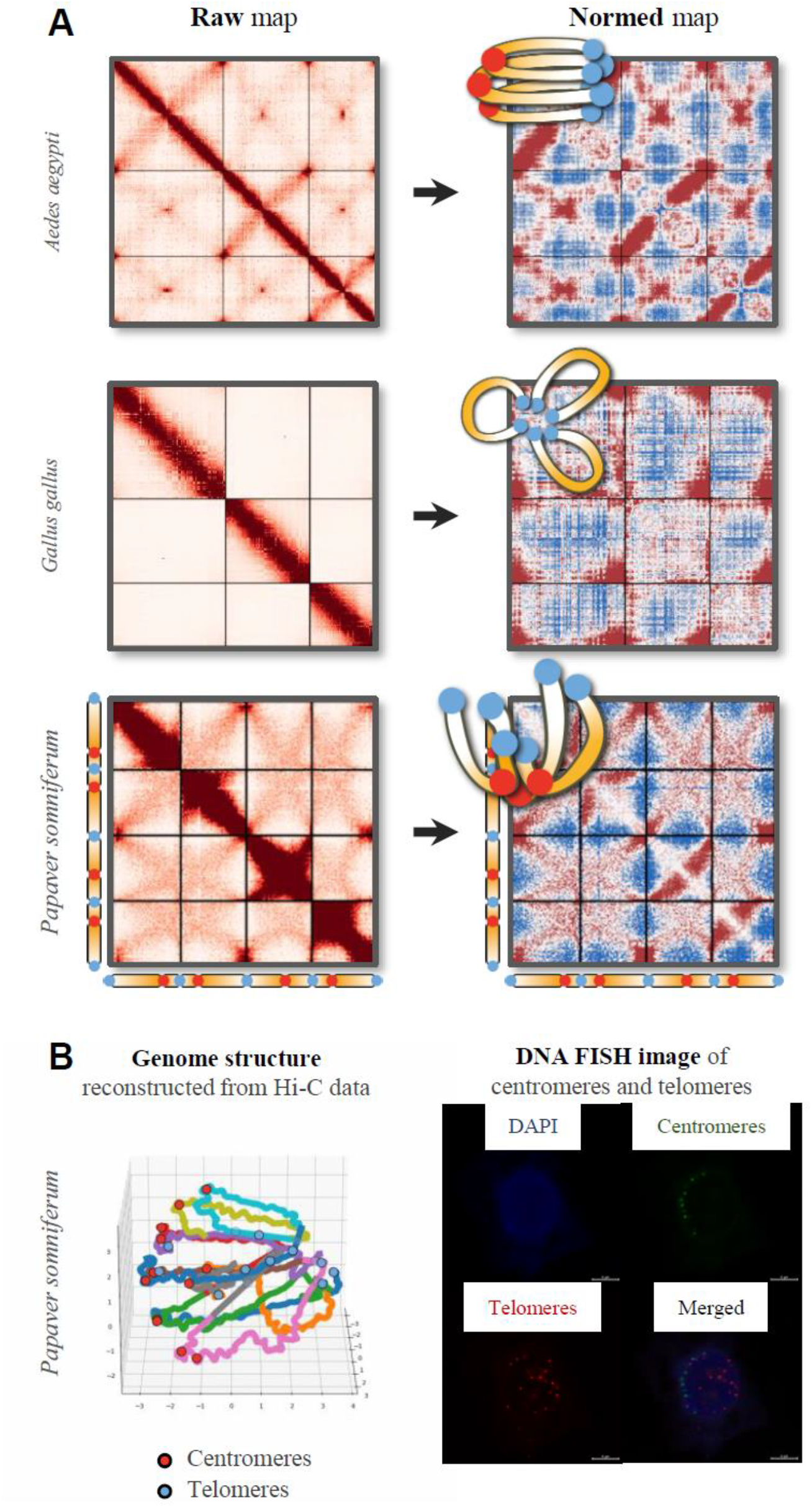
Examples and Integrative Validation of NormDis. A. Raw (left) and NormDis-normalized (right) Hi-C maps of three example species, with their exposed large-scale structures illustrated at the upper left corner. Normalization enhances long-distance interactions and suppresses short-distance interactions. B. NormDis-predicted architecture of *Papaver somniferum*, reconstructed genome structure by Pastis^51^with centromeres and telomeres labeled by colored dots, and DNA FISH images with centromeres and telomeres labeled. The concordance between NormDis predictions, polymer models, and FISH data demonstrates robust cross-validation of genome architecture inference methods in non-model species.

**Figure S3.**
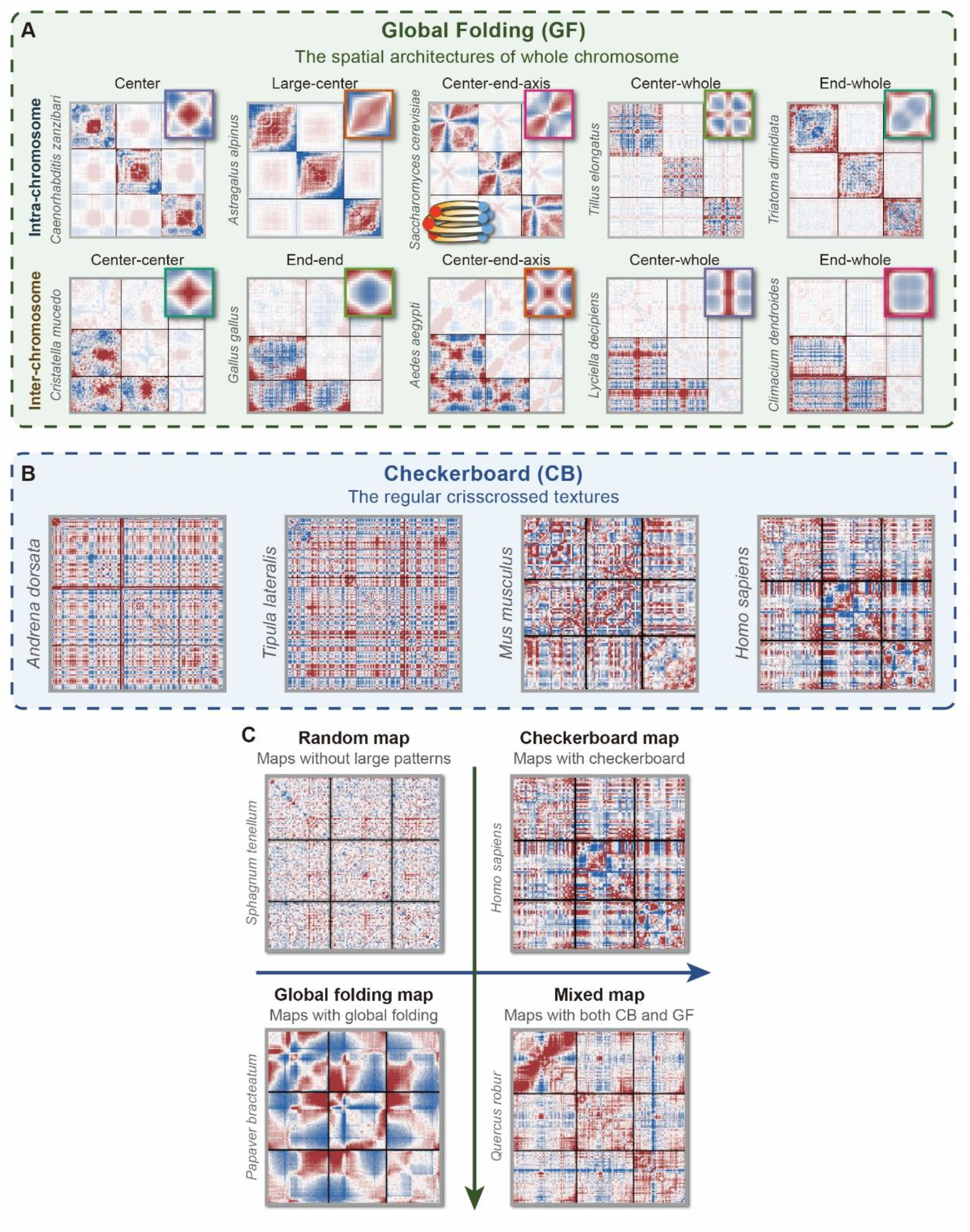
Diversity and Coexistence of High-Order Genome Architectures. A. Spectrum of global folding patterns. Global folding reflects karyotype-scale spatial organization of chromosomes. Unsupervised clustering of Hi-C maps from 1,025 species identified five intra- and five inter-chromosomal types. Representative species for each type are shown, with each displaying its first three chromosomes. The average pattern for each type is plotted in the upper right corners. Genome configuration of yeast (*Saccharomyces cerevisiae*) is illustrated as an example for global folding. B. Checkerboard patterns are the regular crisscrossed textures on maps, which indicate chromatin compartments. Four species with strong checkerboard organization are displayed, with each displaying its first three chromosomes. C. Architectural independence and coexistence. Maps without any high-order architectures (named as random maps), maps with only checkerboard or global folding patterns, and maps with both global folding and checkerboard patterns are shown, displaying the first three chromosomes of each.

**Figure S4.**
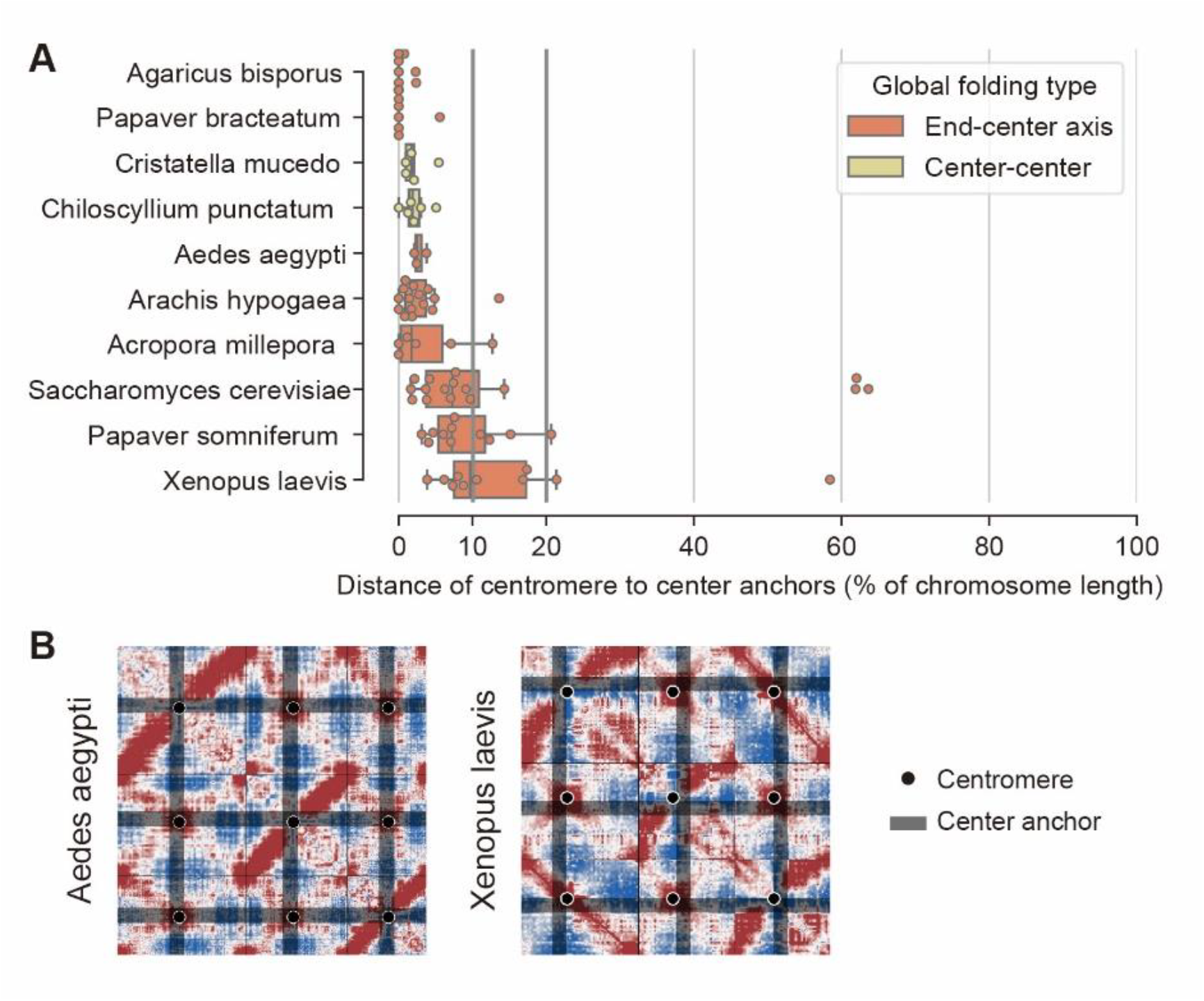
Coupling of Center Anchors and Centromeres. A. Center anchors locate near centromeres, as indicated by the ratio of the distance between center anchors and centromeres corresponds to chromosome length, analyzed in species exhibiting center-associated inter-chromosomal global folding (end-center-axis and center-center types). B. Positions of centromeres and center anchors on two example species, displaying the first three chromosomes of each. Centromeres are marked by the black dots, and the positions of center anchors are marked by the black lines.

**Figure S5.**
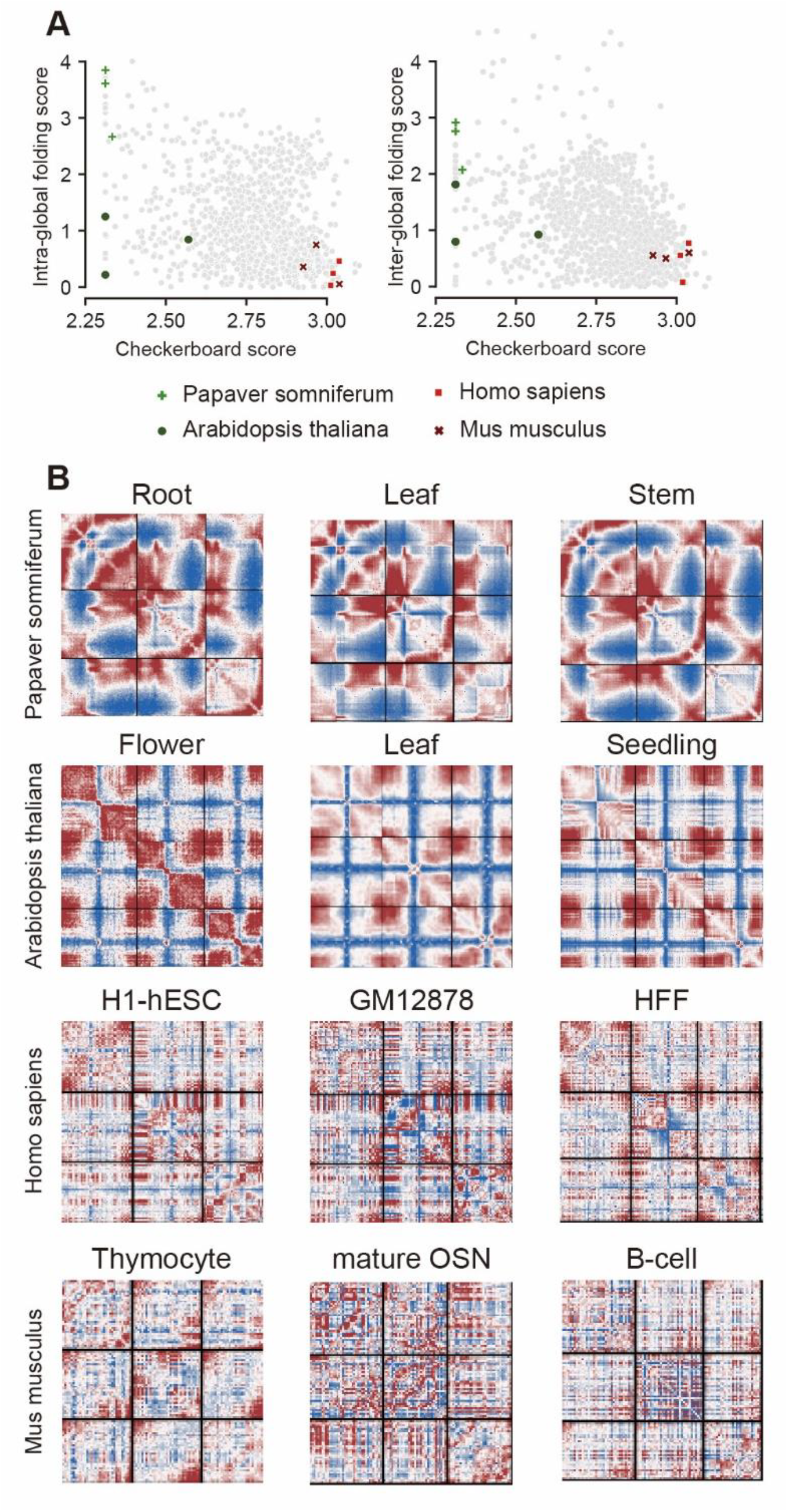
Conservation of high-order genome architectures across mature tissues. A. Architectural conservation across tissues. Scatter plot of global folding scores (GFS) and checkerboard scores (CBS) across tissues from four species, with 1,025 species as background. Detailed scores are documented in Supplementary Table 5. B. Normalized contact maps for exemplified tissues across four species, showing conserved global folding and checkerboard patterns within species. Black lines demarcate chromosome boundaries.

**Figure S6.**
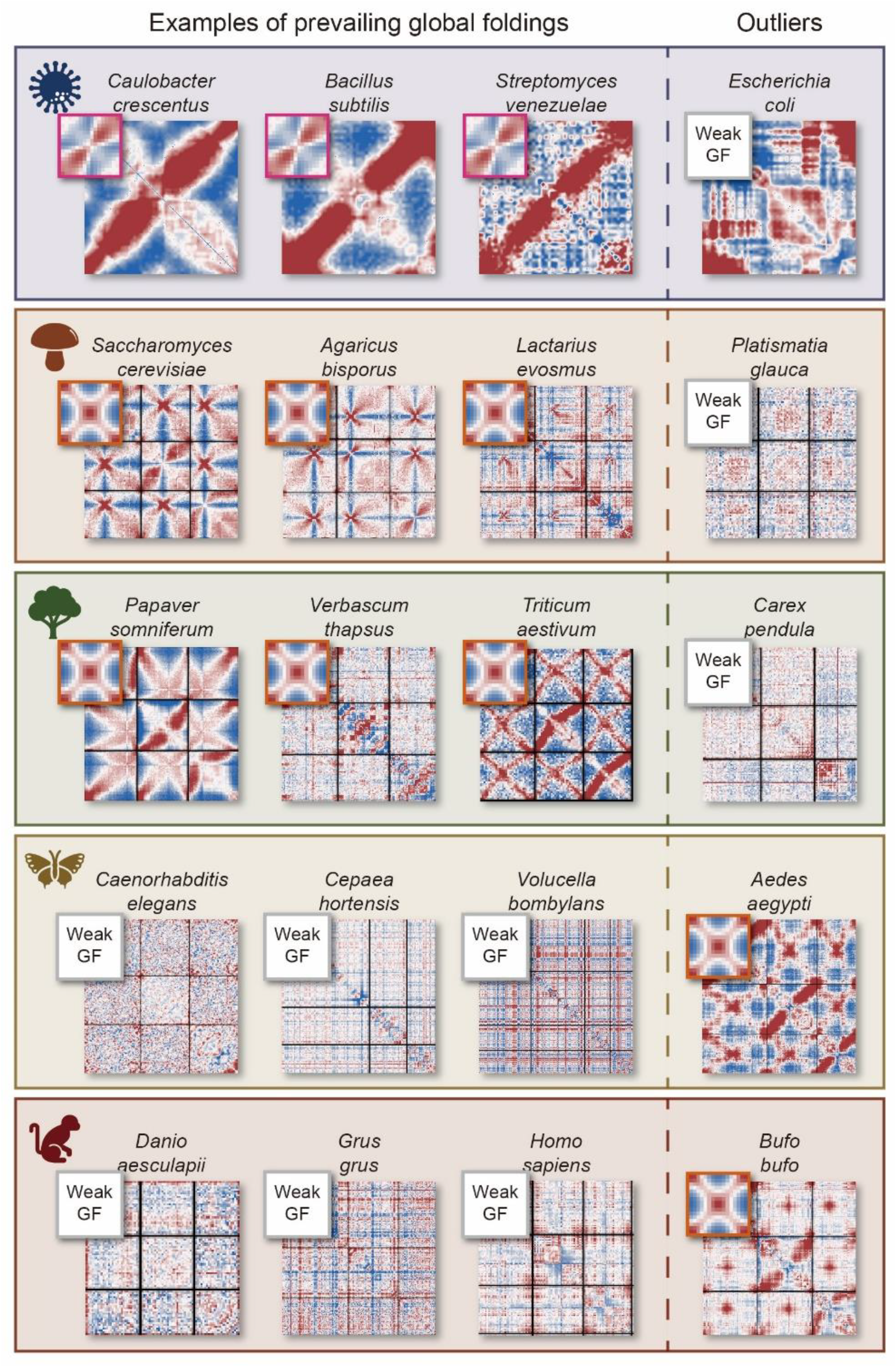
Taxonomic Prevalence and Exceptions in Global Folding. Left three panels: taxonomically dominant patterns, examples of global folding types within major taxonomic groups. Right panel: outlier species exhibiting non-canonical folding types.

**Figure S7.**
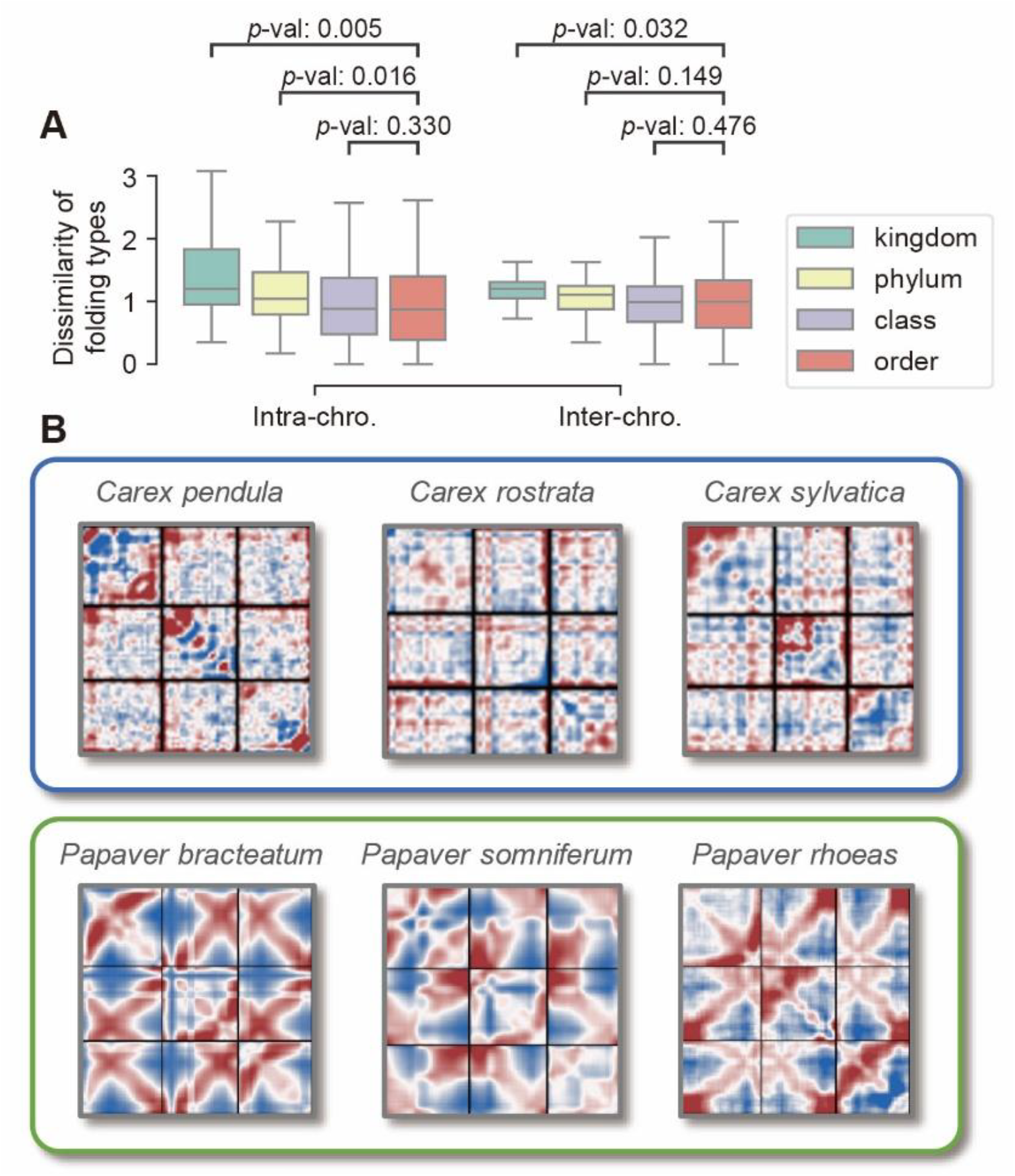
Phylogenetic Conservation and Divergence in Global Folding. A. Architectural variation is restricted by phylogenetic relatedness. Standard deviations (SDs) of global folding scores decrease with finer taxonomic resolution. *P*-values are calculated by the Mann-Whitney U test with a greater alternative hypothesis, which includes 3 kingdoms, 11 phyla, 15 classes, and 33 orders. The reduced sample sizes resulting from finer taxonomic divisions likely explain the non-significant differences between closely related taxonomic divisions (for example between class and order). B. Exemplified lineage-specific global folding. Two lineages from the same kingdom (plants) exhibit distinct architectures. The upper species from *Carex* genus lacks strong global folding patterns, while the lower species from *Papaver* genus exhibit center-end-axis patterns.

**Figure S8.**
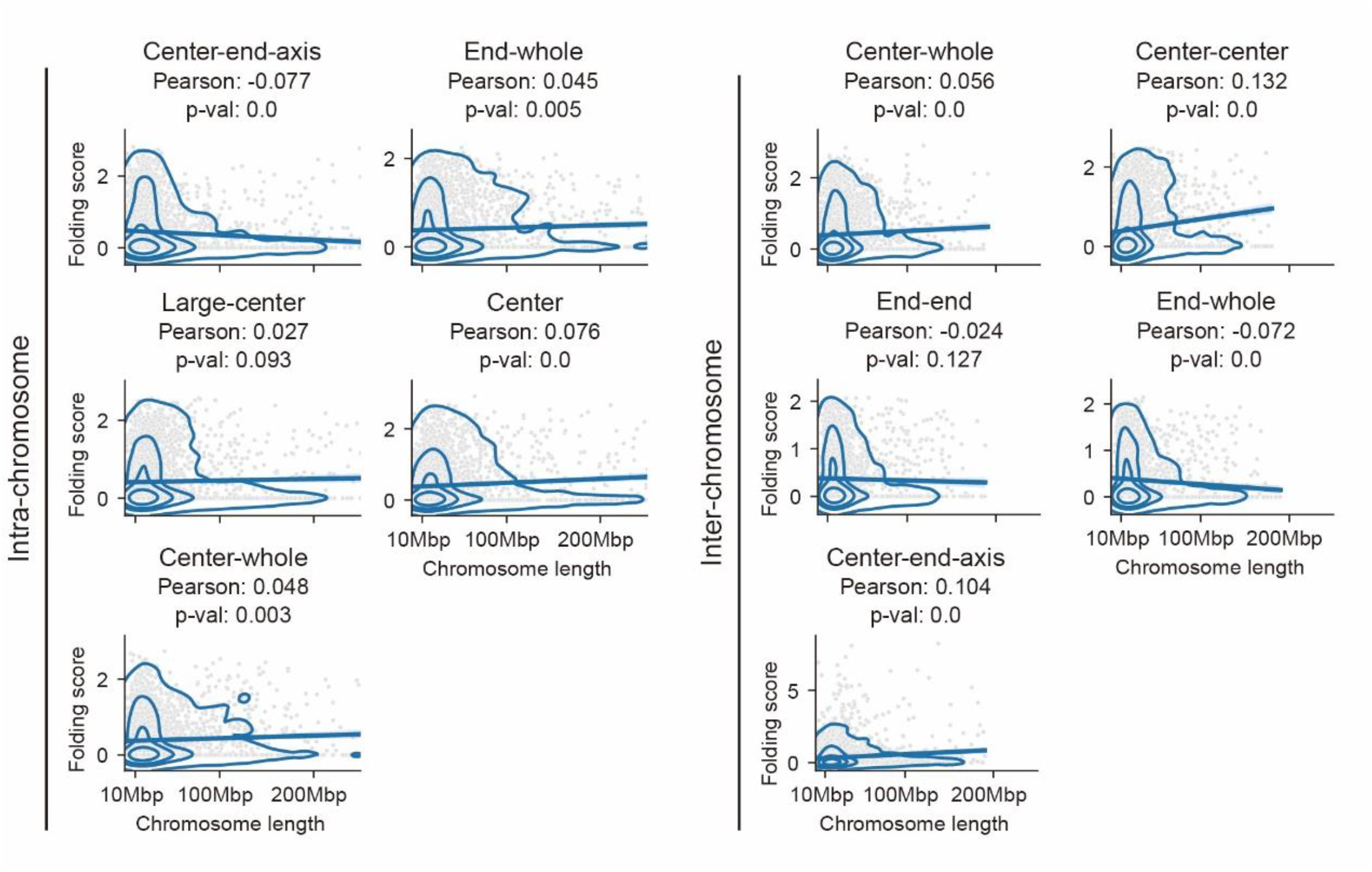
Global Folding Is Uncoupled from Chromosomal Morphology. Pearson correlation analysis reveals no significant association (|r| < 0.13 for all folding types) between global folding scores and chromosomal length. 20,000 randomly sampled chromosomes are shown for each correlation. Results are stratified by five intra-(top row) and five inter-chromosomal (bottom row) global folding types.

**Figure S9.**
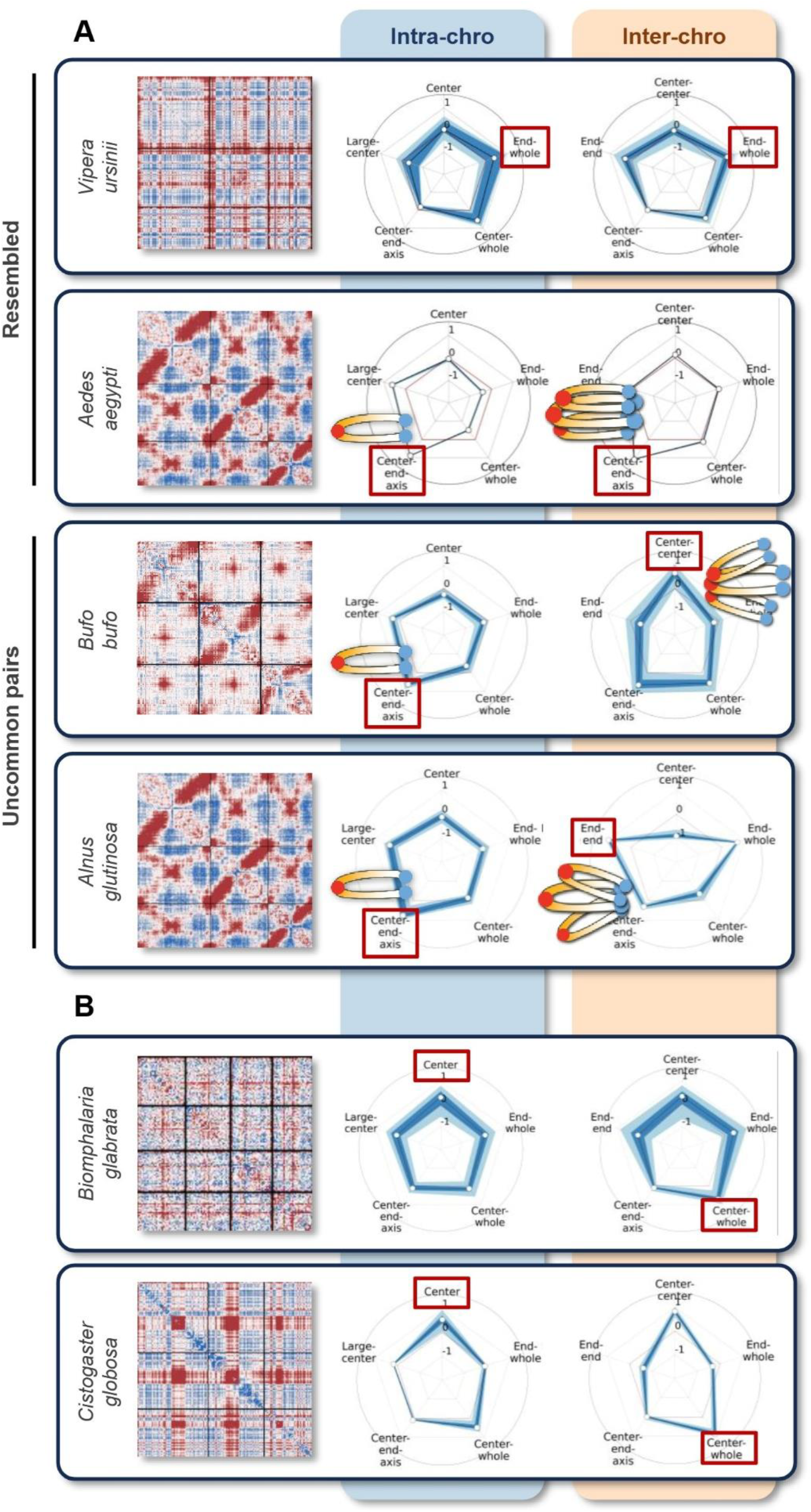
Variety in global folding type interplay. A. Variety in the interplay between intra- and inter-global folding types. The upper two panels exhibit species with resembled global folding pairs. The lower two panels exemplify the uncommon pairs. Normalized maps are shown on left for each species, displaying the first three chromosomes. Rader maps exhibiting the intra- and inter-global folding type strength are shown on right. Illustrations for center-end-axis related types are displayed. The strongest types are marked by the red rectangles. Although the three illustrated species exhibit the same intra-global folding type, center-end axis, the inter-global folding types differs. *Aedes aegypti* displays the typical parallel arrangements of chromosomes, whiles *Bufo bufo* exhibits stronger center anchors clustering but weak end anchors clustering, and *Alnus glutinosa* exhibits the opposite. This indicates the variety in the interplay between intra- and inter-global folding type. B. Example species of different global folding types but sharing the same anchor, displaying the first three or four chromosomes of each. The number of species belonging to each pair is shown in Figure S8. The strongest types are marked by the red rectangles.

**Figure S10.**
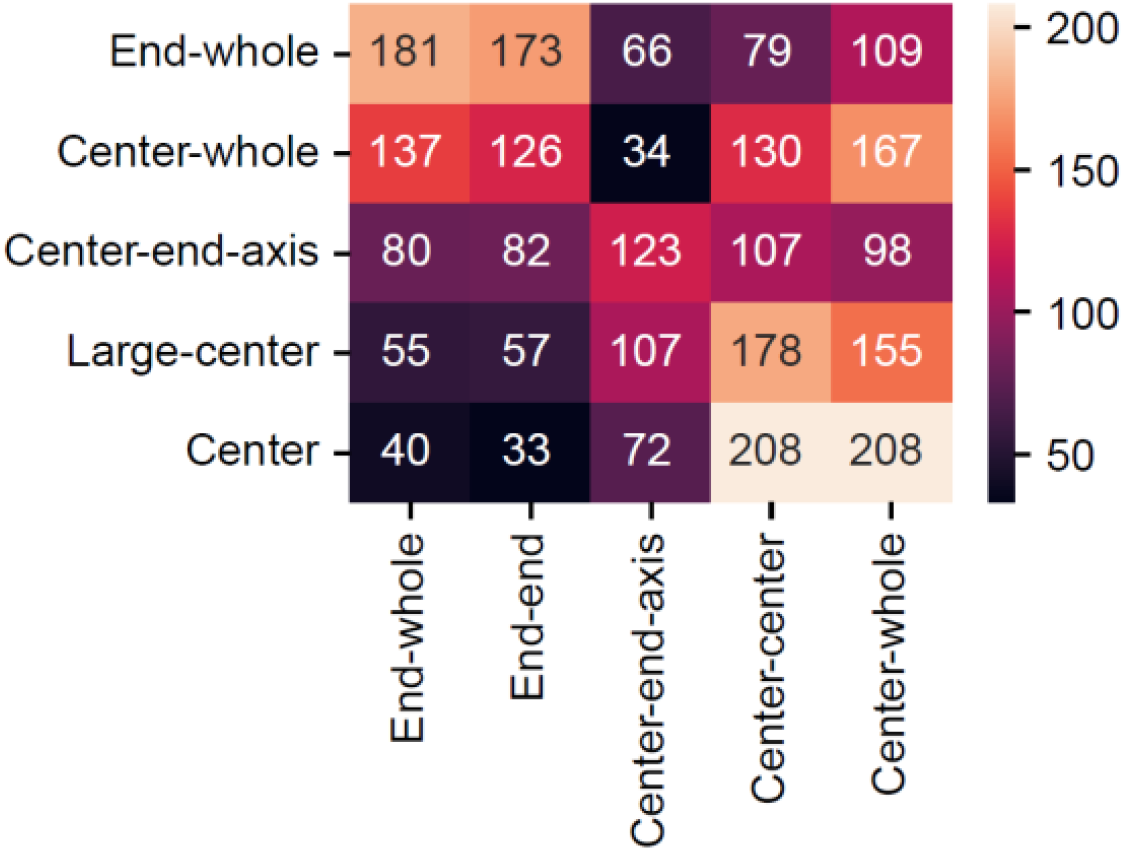
Variety in global folding type interplay (pair number). Analysis of 1,020 species reveals widespread diversity in global folding pairs. Rows: five intra-chromosomal folding types; columns: five inter-chromosomal folding types. Numbers represent species counts for each pairwise combination.

**Figure S11.**
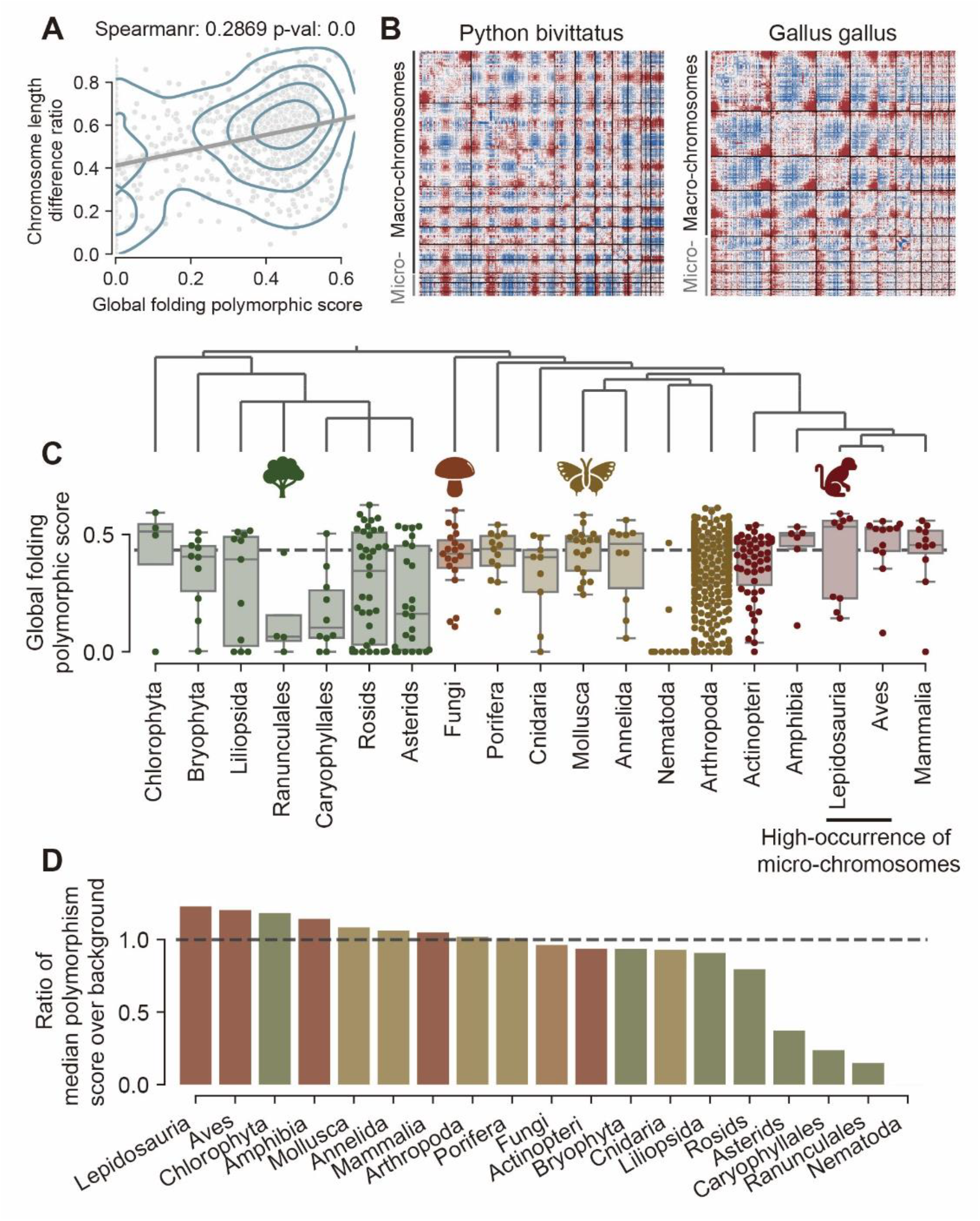
Prevalence of Global Folding Polymorphism. A. Chromosomal length variation associates with folding polymorphism, indicated by the correlation between global folding polymorphic scores and chromosome length heterogeneity across 1,025 species. B. Representative species with global folding polymorphism in addition to Figure 2D. C. Global folding polymorphic score of main taxonomical groups, indicating the broad distribution of polymorphism. D. Sorted ratios of median polymorphic score over background (marked by the black line). Detailed data is shown in Supplementary File 5.

**Figure S12.**
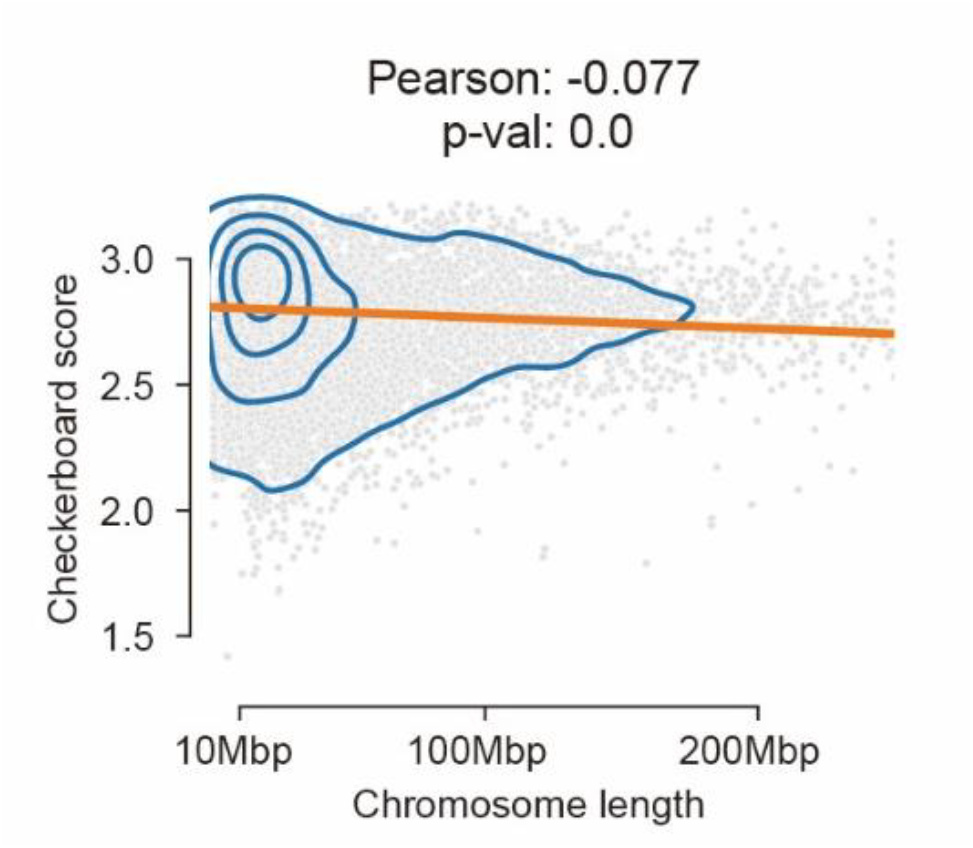
Checkerboard Architecture Is Uncoupled from Chromosomal Morphology. Correlations between chromosomal length and checkerboard score (r = -0.077). 20,000 chromosomes are randomly sampled for efficient plotting.

**Figure S13.**
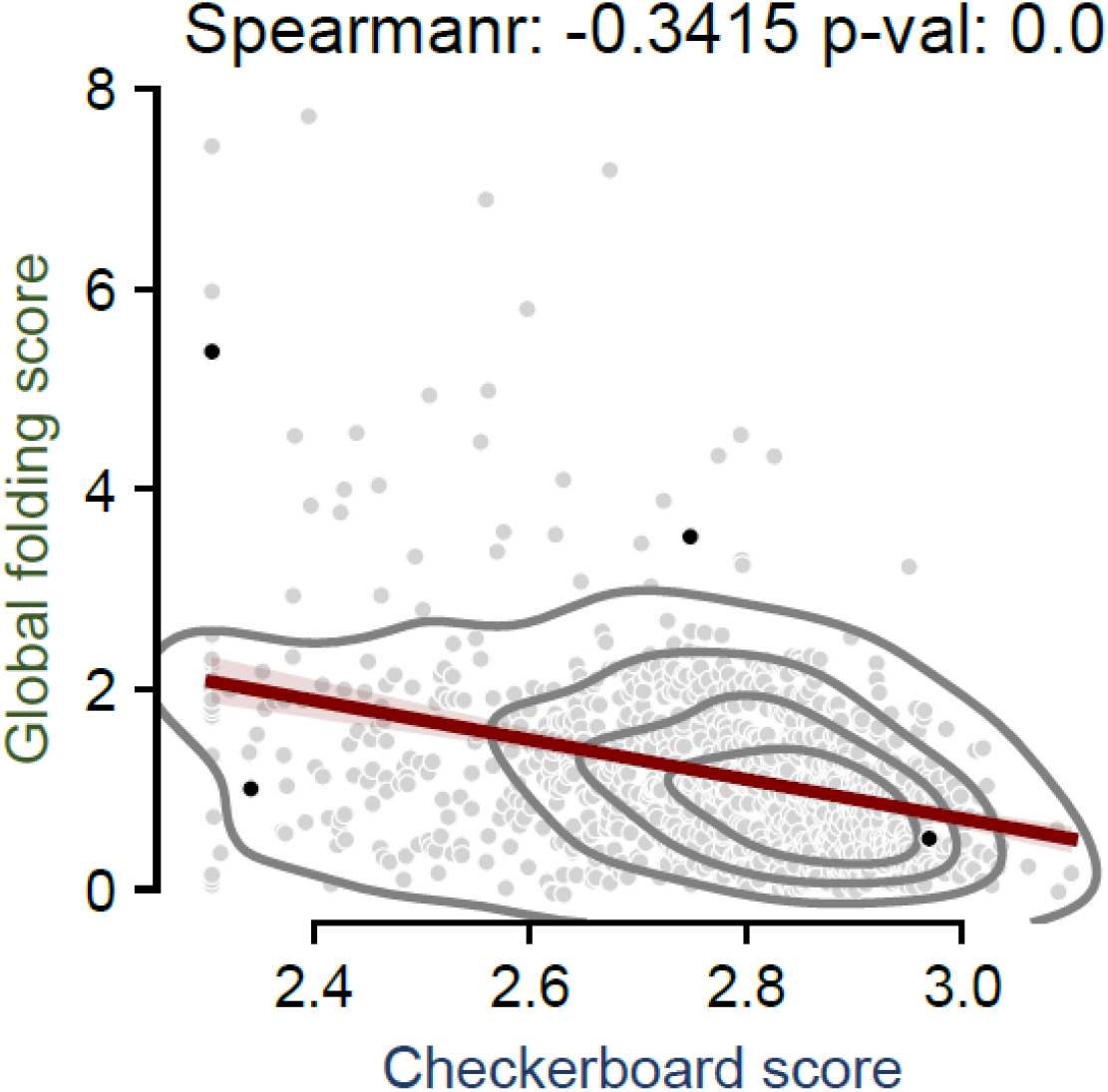
Negative correlation between checkerboard and inter-chromosomal folding. Checkerboard scores exhibit a negative correlation with inter-chromosomal global folding scores across 1,025 species (Spearman ρ = −0.3415, p < 1×10−15). Correlation with intra-global folding score is shown in Figure 5A. The four example species shown in Figure 5A are marked by the black dots in this plot. This suggests the competing relationship between checkerboard and global folding.

**Figure S14.**
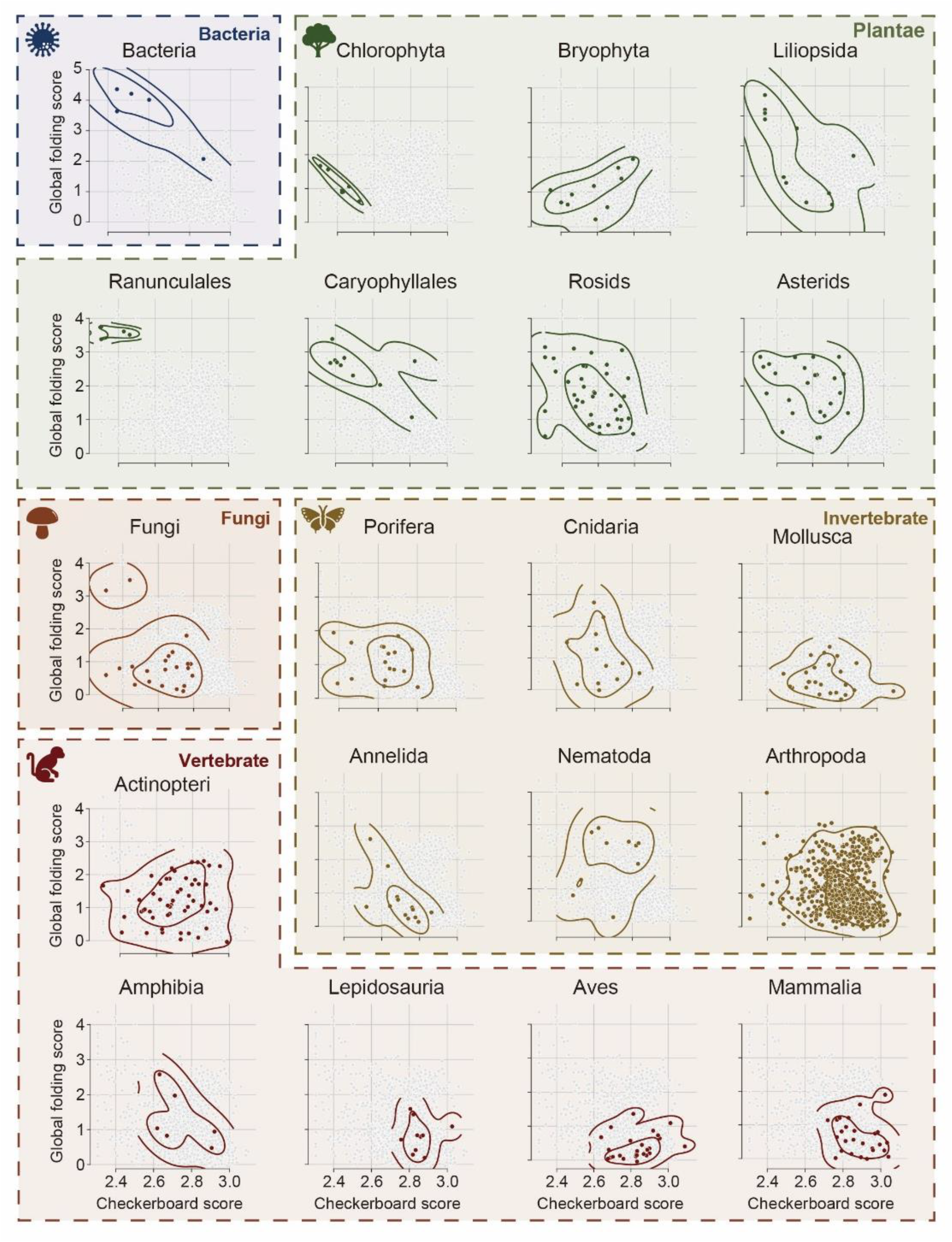
The evolutionary trajectory of 3D genome as shown by checkerboard score and intra-chromosomal global folding scores. Scatter plot of checkerboard scores versus intra-global folding scores for 21 taxonomic groups (group sizes shown in Fig. 1A), overlaid on the background distribution of 1,025 species (gray points).

**Figure S15.**
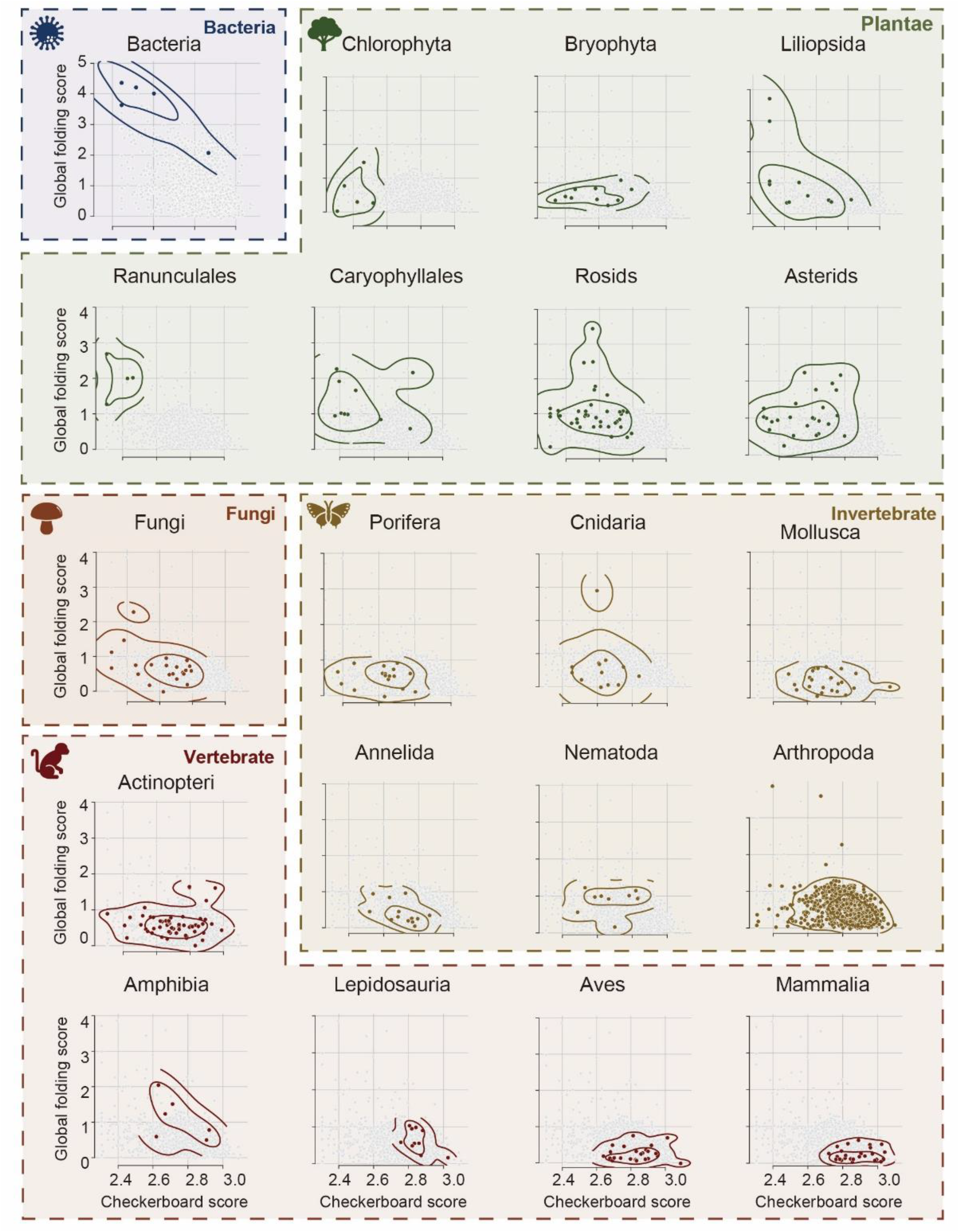
The evolutionary trajectory of 3D genome as shown by checkerboard score and inter-chromosomal global folding scores. Scatter plot of checkerboard scores versus inter-global folding scores for 21 taxonomic groups (group sizes shown in Fig. 1A), overlaid on the background distribution of 1,025 species (gray points).

**Figure S16.**
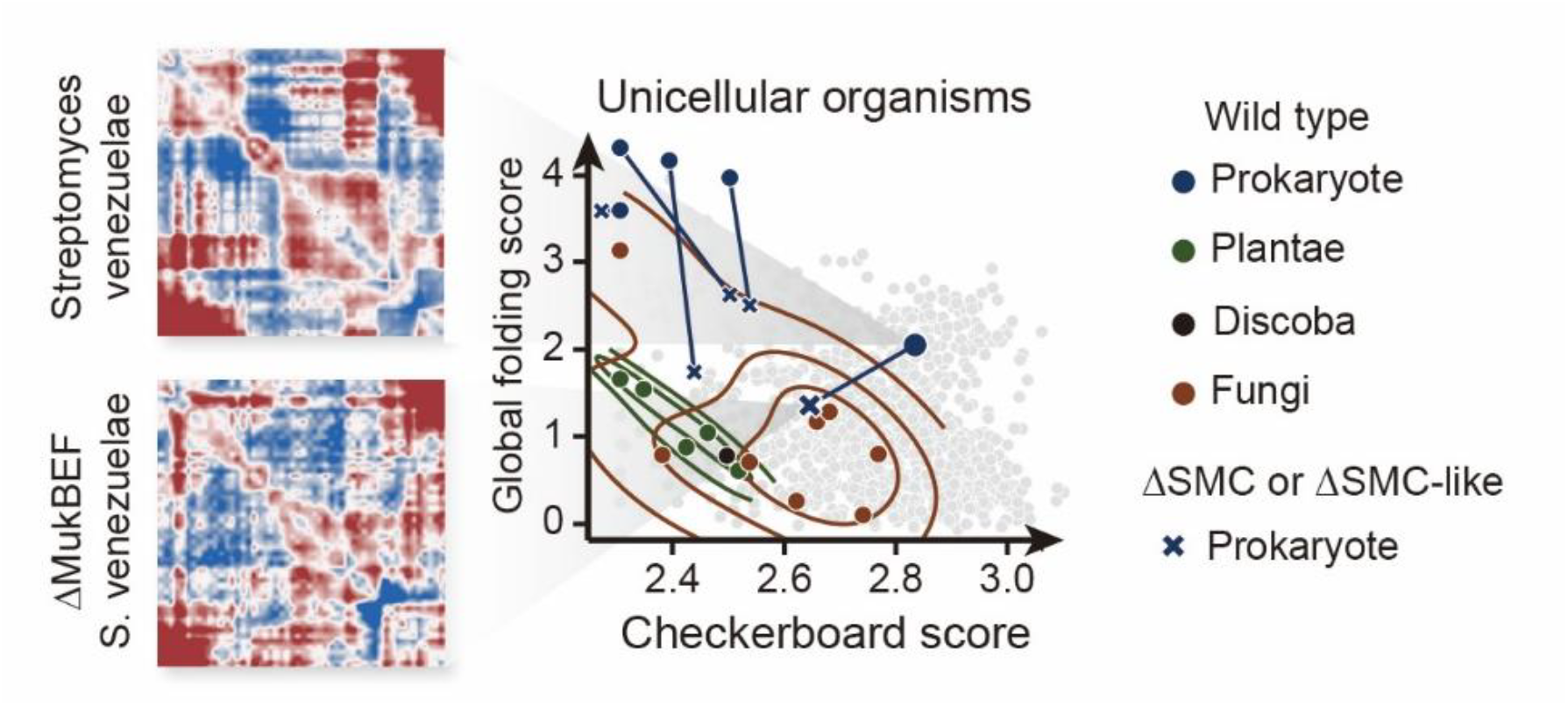
MukBEF Drives Checkerboard Patterns in *E. coli*. Normalized Hi-C maps of *Escherichia coli* (wild-type) and its MukBEF knockout mutant. Wild-type exhibits the interaction stripes reflecting spatial compartmentalization, while the mutant shows weaker patterns.

**Figure S17.**
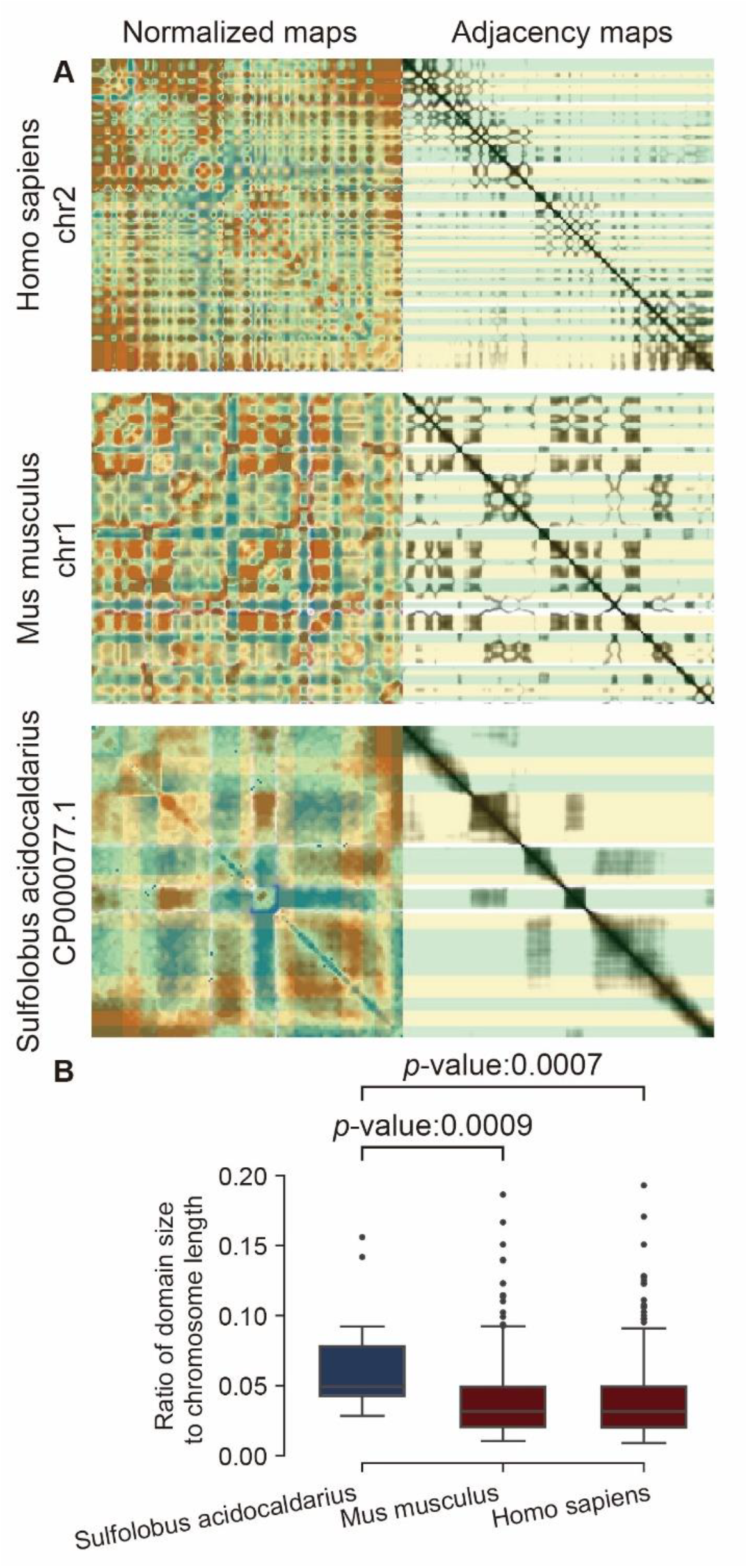
Checkerboard Domain Sizes among Species. A. Comparative visualization of checkerboard domain sizes. Normalized Hi-C maps (left) and adjacency matrices (right; by cosine distance) from *Homo sapiens* (Chr. 2), *Mus musculus* (Chr. 1), and *Streptomyces venezuelae*. Compartment domains are marked by the yellow or green boxes. B. Quantitative scaling of compartment size. Distribution of compartment-to-chromosome length ratios for human, mouse and *S. venezuelae*, as measured by their ratio to chromosome length. 13 domains are found in *S. venezuelae* (median ratio 4.96%), 385 in mice (3.17%; Mann Whitney U test *p*-value: 0.0009) and 415 in human (3.14%; 0.0007).

**Figure S18.**
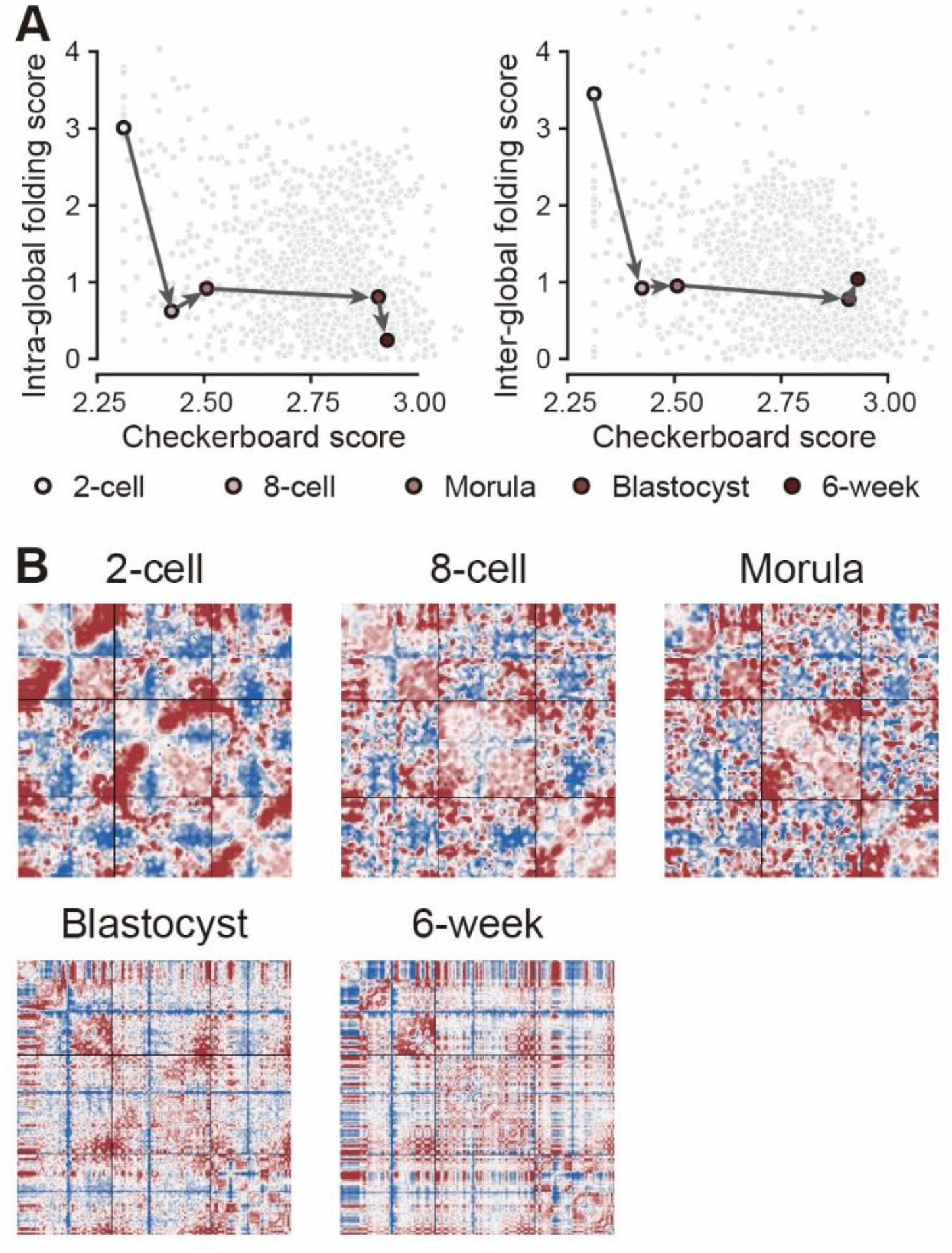
Architectural Transition During Human Embryogenesis. A programmed transition of high-order architectures is observed during human embryogenesis. It indicates that global folding maintains genome plasticity for totipotency, while checkerboard architecture enables cell fate determination through spatially coordinated gene regulation. A. Developmental trajectory of 3D genome reorganization. Scatter plots trace the progression from global folding to checkerboard patterns during early human embryogenesis. The developmental order is indicated by the arrows, with each stage is labeled below. B. Normalized maps of human early embryo development, displaying the first three chromosomes of each.

**Figure S19.**
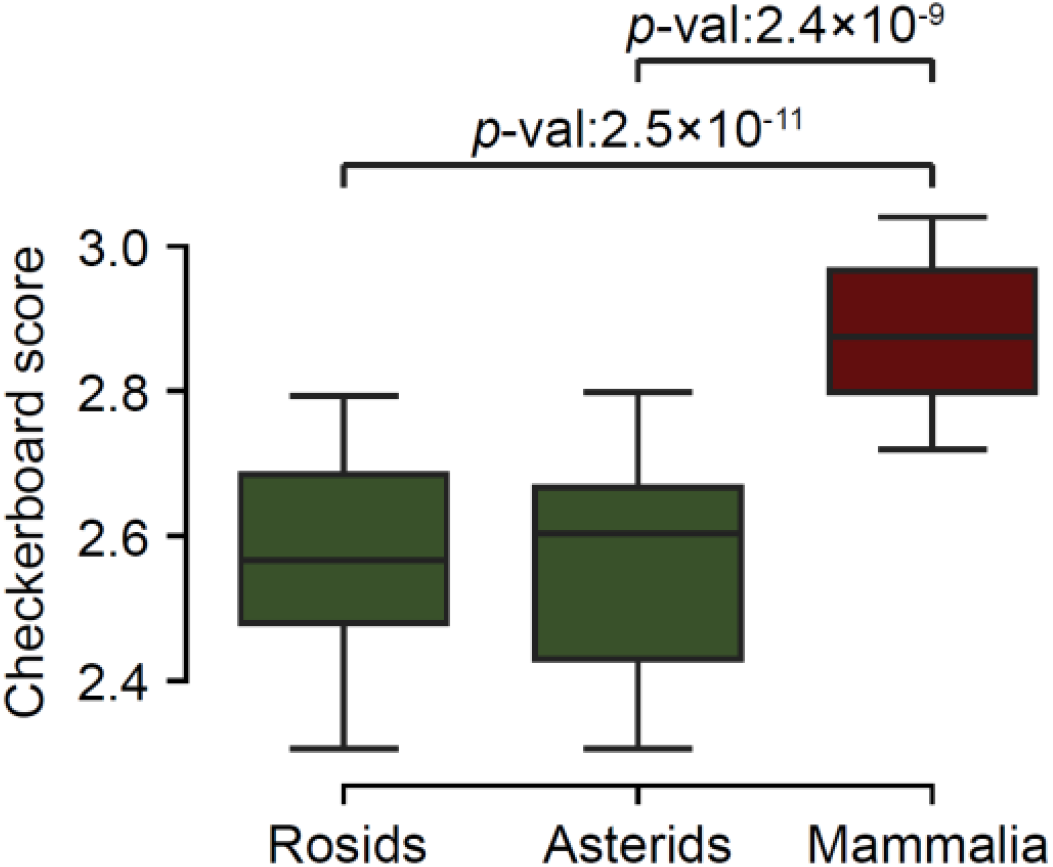
Checkerboard Strength across Plant and Animal Lineages. Boxplot of checkerboard scores between Rosids (n=39), Asterids (n=24) and Mammalia (n=23). *P*-value is calculated by Mann-whitney U test. Checkerboard scores differ significantly between Mammalia (median CBS = 2.87) and eudicot clades (Rosids: 2.57, Asterids: 2.60).

**Figure S20.**
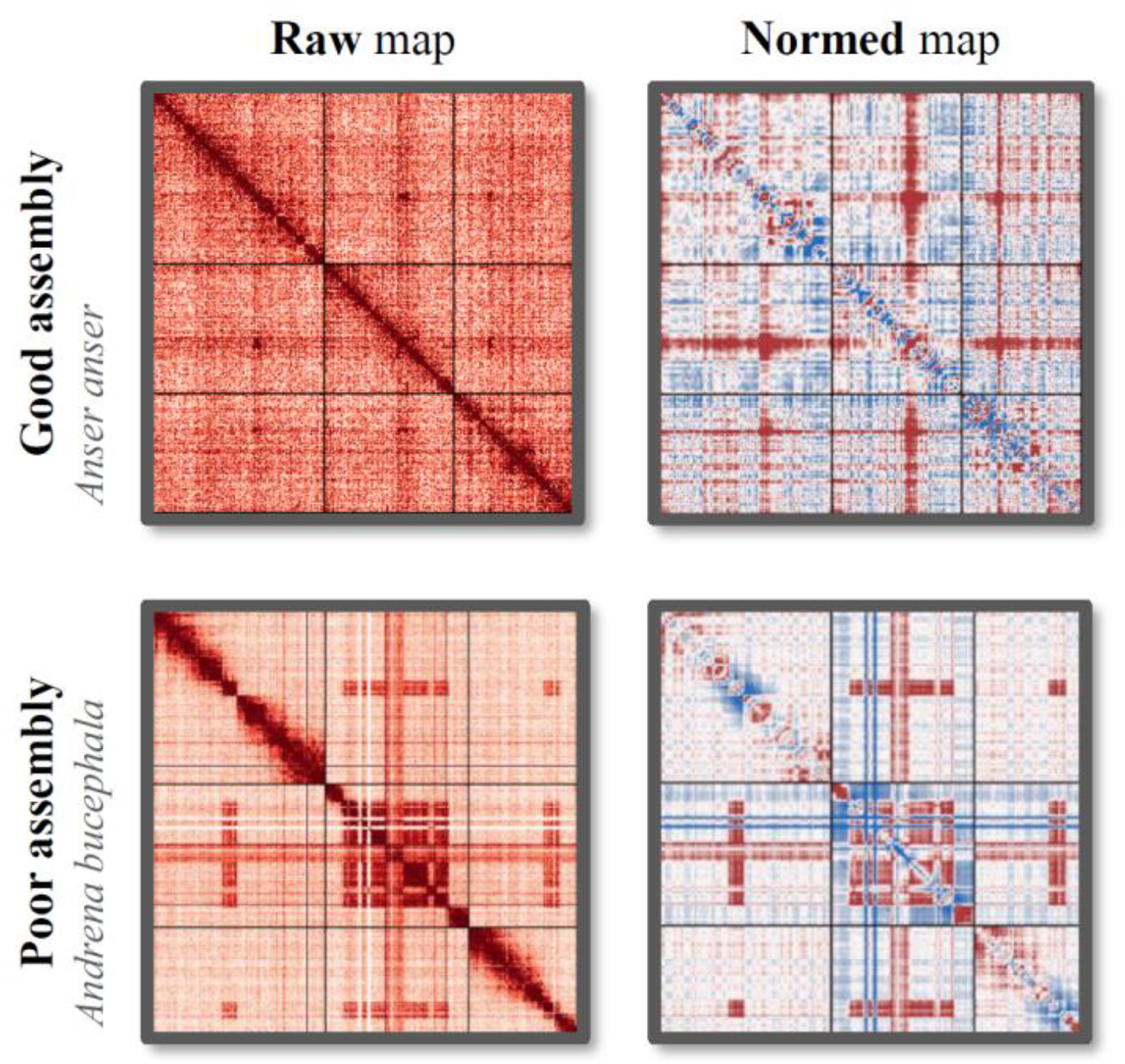
Assembly Quality Assessment Based on Hi-C. Top: Example species of high-quality assemblies. Species with center-whole global folding pattern exhibit gradual interaction decay from interaction-dense genomic bins without sharp transition. Bottom: Example species of poor assemblies. oow-quality assemblies display sharp artifactual interaction spikes at erroneous scaffold junctions. High-quality Hi-C assemblies are prerequisite for resolving true chromatin architectures, as poor assemblies introduce topological biases.

**Figure S21.**
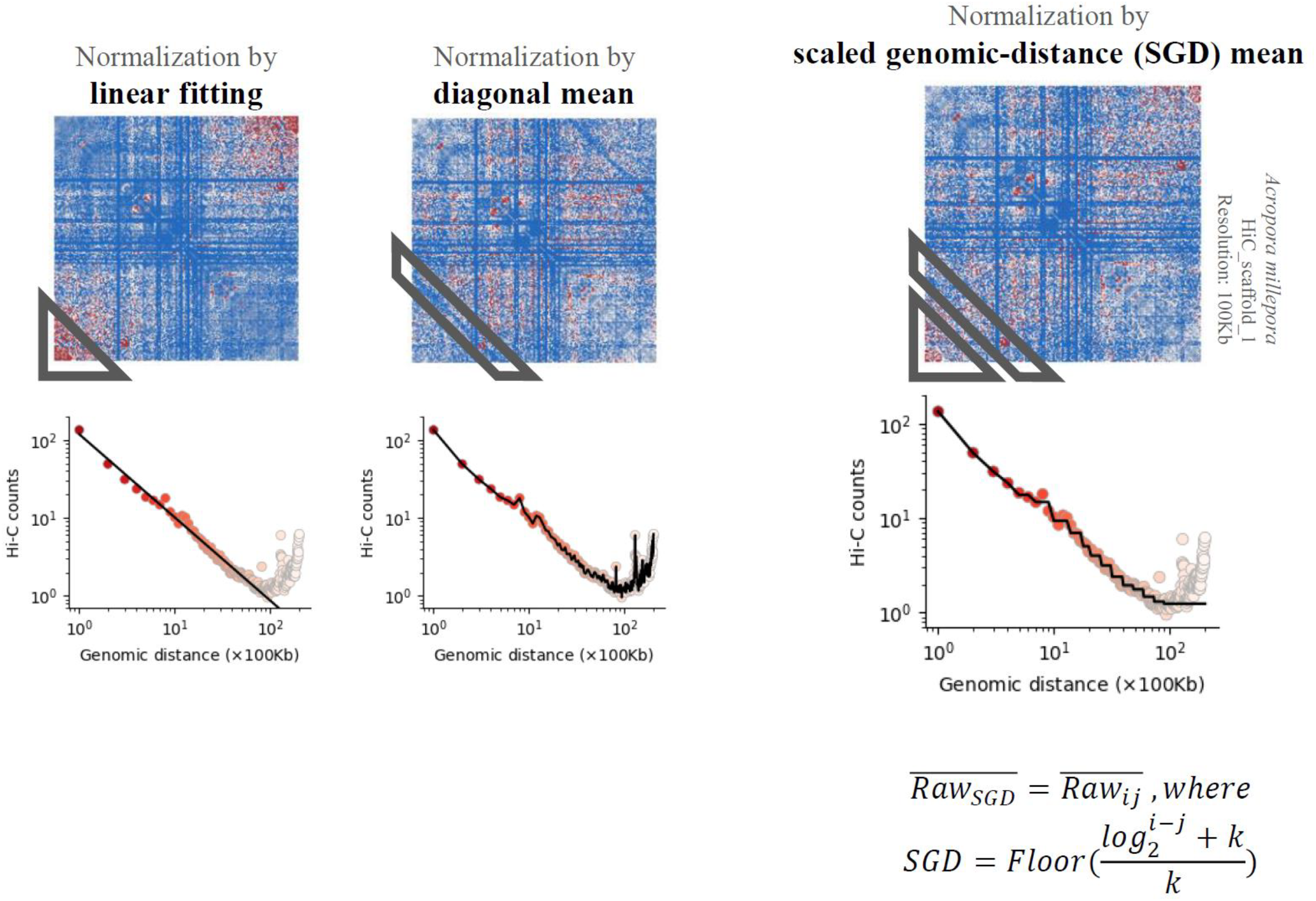
Comparative performance of Hi-C normalization strategies. Top: Normalized maps of *Acropora millepora* (Chromosome 1) processed by three normalization strategies. Bottom: Interaction frequencies (Red dots) and expected frequencies (black lines). NormDis with SGD normalization strategy exhibit most accuracy in removing distance biases, avoiding overcorrection (linear fitting) or outlier susceptibility (diagonal mean).

1. Linear fitting: Artificially elevates long-distance interactions, marked by the gray triangle at the lower left corner.
2. Diagonal mean: Distorted by a long-distance outlier, marked by the gray quadrilateral.
3. NormDis: Corrects both biases, preserving true biological signals. Detailed information is described in Methods.

**Figure S22.**
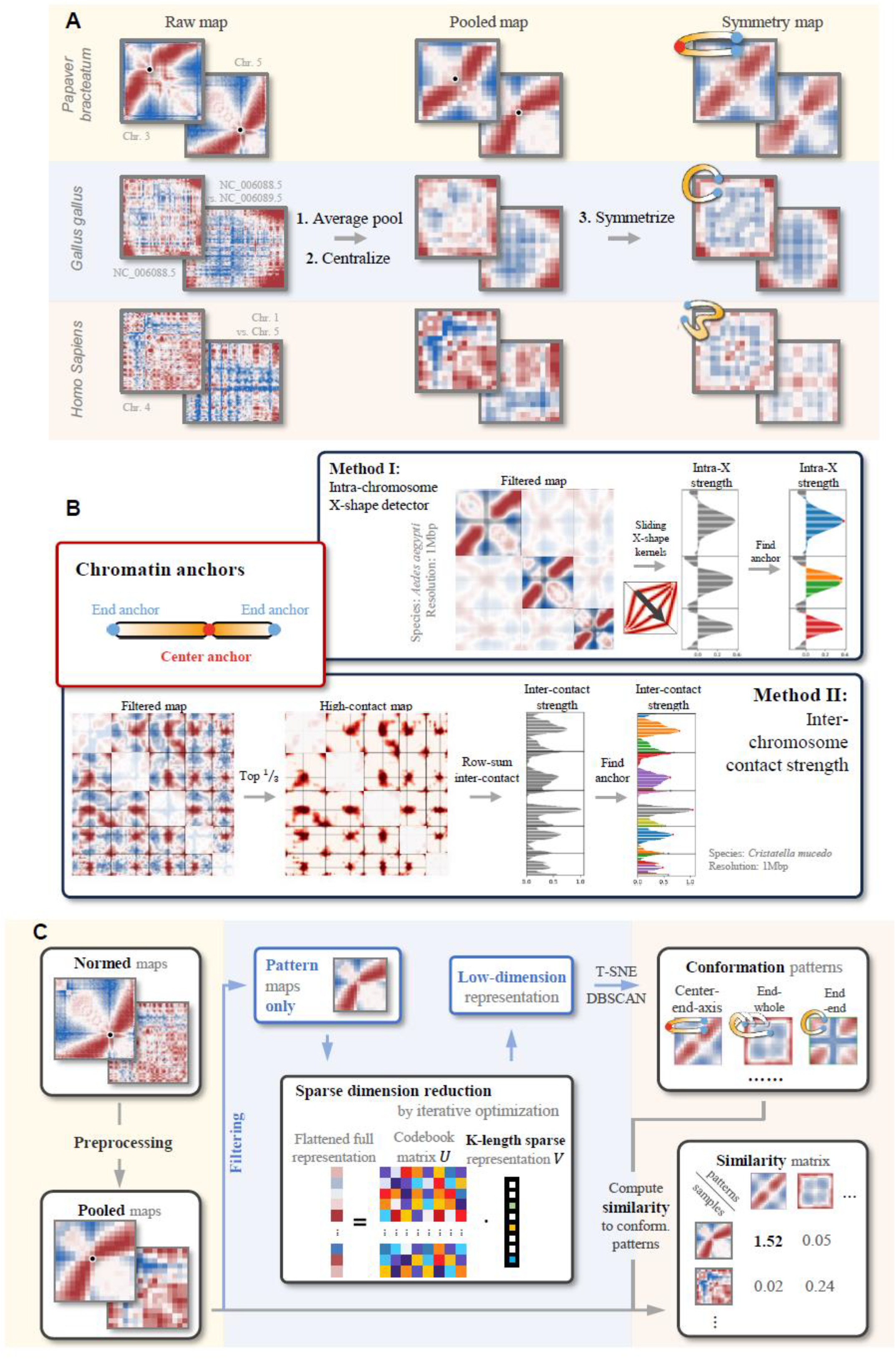
Systematic Pipeline for Global Folding Pattern Mining. A. Preprocessing for large-scale pattern enhancement, including average pooling, centralization and symmetrization, detailed in Methods. Representative examples include *Papaver somniferum* (center-anchor), *Gallus gallus* (end-anchor) and *Homo sapiens* (no obvious global folding). B. CenterFinder algorithm for center anchor detection. Dual-mode computational framework identifies folding centers: “Intra X” mode is optimized for center-end-axis patterns, while “Inter row sum” mode is optimized for anchors with elevated inter-chromosomal contacts. C. Overview of global folding pattern mining. Four-stage analytical workflow includes training set curation, pre-processing, pattern mining and quantification of global folding. This pipeline resolves global folding architectures, enabling cross-species comparison of large-scale genome organization.

**Figure S23.**
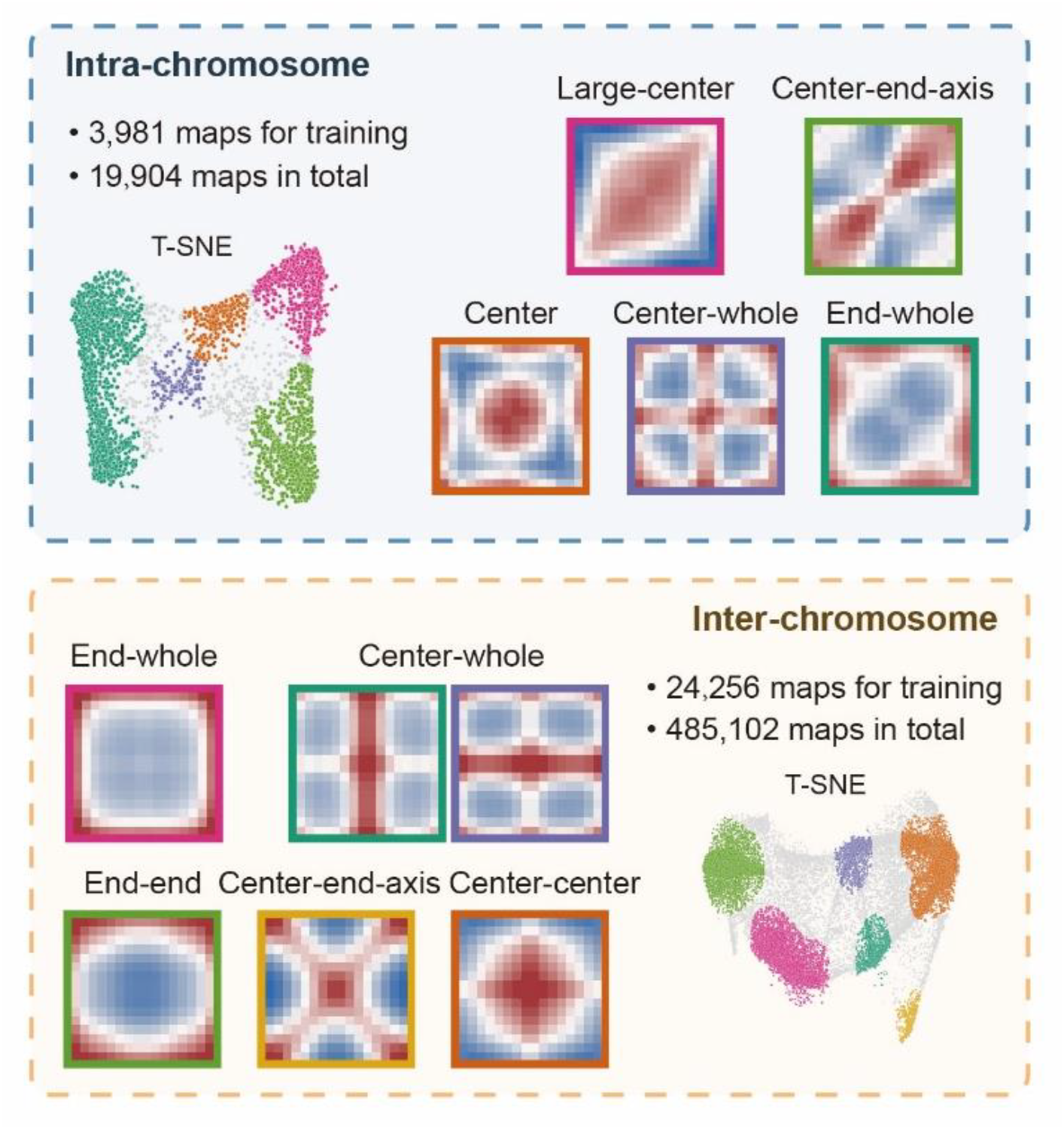
Robust Reproducibility of Global Folding Pattern Mining. Global folding patterns were re-identified through independent runs with randomized dictionary matrix initialization and automated training set selection. The conserved recovery of global folding patterns across randomized runs validates our pipeline’s algorithmic stability and insensitivity to initial conditions.

**Figure S24.**
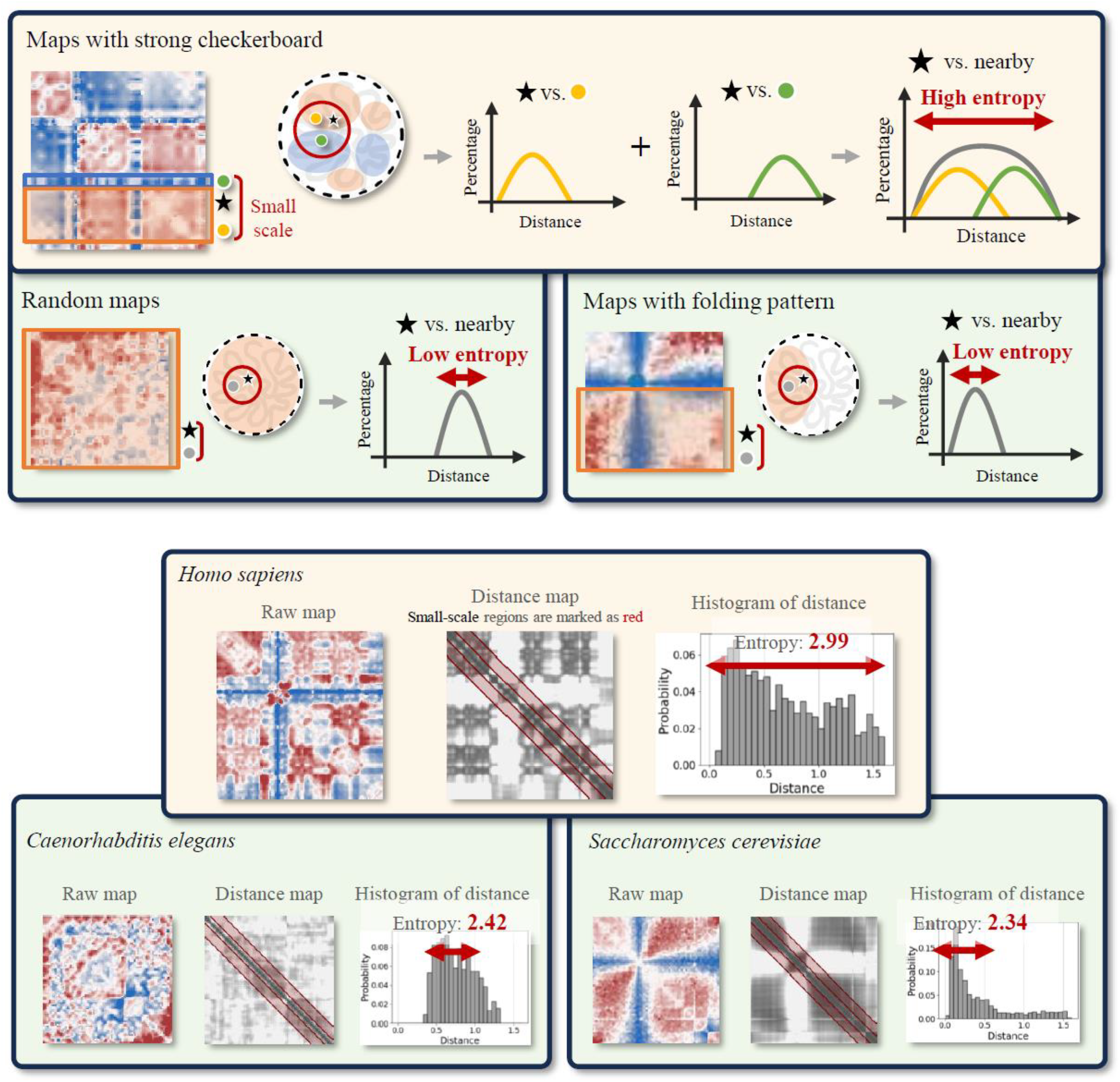
Quantitative Framework for Checkerboard Patterns. Illustrations for methods of quantifying checkerboard patterns. The upper panel represents three chromatin organization paradigms: checkerboard-dominant, random and global folding-dominant. Illustrations of the underlying physical structures are shown in black edged circles, with colored circles denoting the compartments. The lower panel provides three example species, representing the three types of maps: checkerboard-dominant (*Homo sapiens*, high entropy 2.99), global folding-dominant (*Saccharomyces cerevisiae*, low entropy 2.34) and random (*Caenorhabditis elegans*, low entropy 2.42). This entropy-based approach objectively quantifies compartmentalization strength.

**Figure S25.**
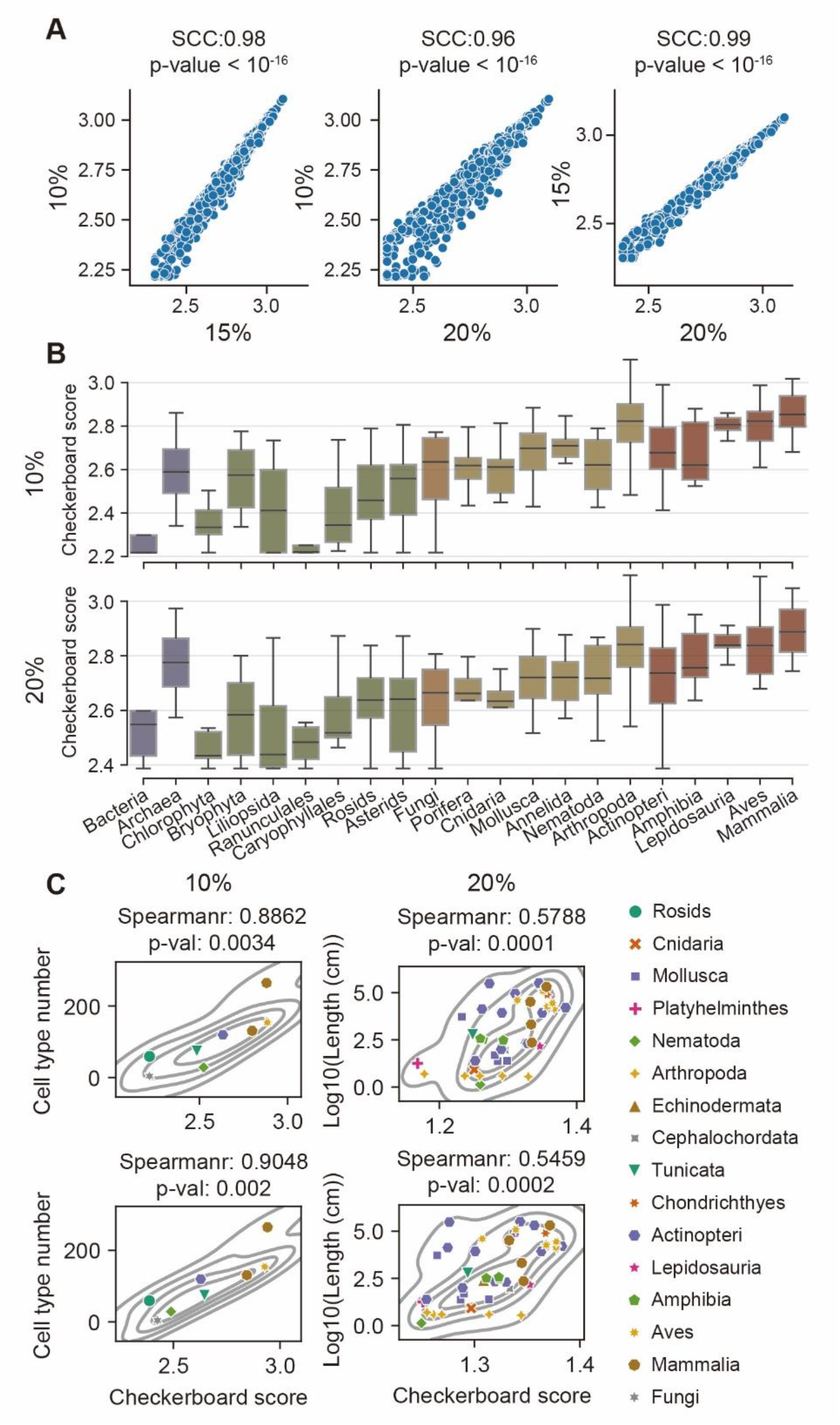
Threshold-Independent Robustness of Checkerboard Quantification. A. High consistency across long-distance thresholds of checkerboard quantification. Checkerboard scores exhibit strong correlation between 10%, 15%, and 20% thresholds across 1,025 species: 10% vs. 15% (Spearman correlation coefficient, SCC = 0.98, *p* <10e-16), 10% vs. 20% (SCC: 0.96, *p* <10e-16), and 15% vs. 20% (SCC: 0.99, *p* <10e-16). B. Consistency of checkboard scores in taxonomic groups. Checkerboard scores for 21 taxonomic groups remain concordant across thresholds. Results of 15% interaction similarity thresholds across 1,025 species are shown in Figure 3. C. Consistency of correlations between checkboard scores and species complexity. Correlations of 15% interaction similarity thresholds across 1,025 species are shown in Figure 3. The threshold invariance of checkerboard quantification validates its reliability for cross-species comparisons.

## Supplementary Tables

**Supplementary Table 1**

Hi-C dataset information and processing details.

**Supplementary Table 2**

Intra-global folding scores (GFS) of 1,025 species.

**Supplementary Table 3**

Inter-global folding scores (GFS) of 1,025 species.

**Supplementary Table 4**

Checkerboard scores (CBS) of 1,025 species.

**Supplementary Table 5**

Global folding scores (GFS) and checkerboard scores (CBS) for mature tissues from same species.

**Supplementary Table 6**

Global folding scores (GFS) and significance of Wilcoxon signed-rank test for the main five taxonomic group.

**Supplementary Table 7**

Differences of global folding scores across different taxonomic levels.

**Supplementary Table 8**

Global folding polymorphism scores for 21 taxonomic group.

**Supplementary Table 9**

Global folding scores (GFS) and significance of Wilcoxon signed-rank test for 21 taxonomic group.

**Supplementary Table 10**

Species complexity as proxied by cell type number (CTN) or body length (kg).

**Supplementary Table 11**

Checkerboard scores (CBS) and significance of Wilcoxon signed-rank test for 21 taxonomic group.

**Supplementary Table 12**

CenterFinder modes used for each species.

**Supplementary Table 13**

Manually re-chose center anchors for each species.

