## Supplementary file for "The evolution of high-order genome architecture revealed from 1,000 species"

${NC}_{ij}=\frac{C_{ij}}{\overline{C_{d}}}$,

where $C_{ij}$ is the raw Hi-C contact of row $i$ and column $j$, ${NC}_{ij}$ is the normalized contact, $\overline{C_{d}}$ is the expected contact of distance $d=\left| i-j \right|$.

Here, we developed a scaled diagonal method that aggregates long-distance diagonals to ensure computational robustness while preserving accuracy, by

$\overline{C_{d}}=Mean\left( C_{d^{'}} \right), whered^{'}=\left\{ d_{i} \right|Floor\left( \frac{{log}_{2}^{d}+k}{k} \right)=Floor\left( \frac{{log}_{2}^{d_{i}}+k}{k} \right)\}$.

where $k$ controls the number of diagonals to be aggregated. Notably, based on the linear nature of chromosome, expected contacts should be theoretically decreasing monotonically with genomic distances. However, some species, for example *Acropora millepora*, exhibit anomalously elevated long-range contacts, likely reflecting its unique global folding patterns (Figure S21). To preserve these signals, we enforce monotonic decay in the expected contact values.

$\min_{U,V}\left\| X-UV \right\|_{F}^{2}+\lambda\left\| V \right\|_{1}$,

where $X=\left[ x_{1},x_{2},\cdots x_{N} \right]$ represents $N$ input maps; $V=\left[ v_{1},v_{2},\cdots v_{N} \right]$ represents the sparse representation for all maps; and $\lambda$ controls the trade-off between reconstruction accuracy and sparsity of $V$.

It is worth noting that the above formulation directly imposes sparsity on $V$, which may potentially lead to suboptimal solutions. Therefore, we introduce an auxiliary variable $W$ to serve as a sparse representation of $V$. The inclusion of $W$ decouples the sparsity constraint from the reconstruction term, enabling explicit sparsity control via an $l_{1}$-norm regularization while promoting structured and interpretable representations. Then to solve the proposed optimization problem, we employ a bi-level optimization strategy, i.e., to update each variable iteratively while keeping others fixed. The optimization process consists of three key steps: updating $V$, updating $W$, and updating $U$.

1. Updating $V$. Given fixed $U$ and $W$, we solve the following subproblem for update $V$:

$\min_{V}\left\| X-UV \right\|_{F}^{2}+\beta\left\| V-W \right\|_{F}^{2}$.

This is a least-squares problem with a closed-form solution:

$V^{+}=\left( U^{T}U+\beta I \right)^{-1}\left( U^{T}X+\beta W \right)$,

where $I$ represents the identity matrix.

2. Updating $W$. Given $V$, the sparse representation $W$ is updated by solving:

$\min_{W}\beta\left\| V-W \right\|_{F}^{2}+\lambda\left\| W \right\|_{1}$.

The optimization can be efficiently solved using the element-wise soft-thresholding function:

$W^{+}=\min\left( V+\frac{\lambda}{2\beta}, max\left( V-\frac{\lambda}{2\beta}, 0 \right) \right)$.

3. Updating $U$. Finally, the dictionary $U$ can be updated by solving:

$\min_{U}\left\| X-UV \right\|_{F}^{2}$,	s.t. $\sum_{i} U_{ij}=1$.

The closed-form solution is given by:

$U^{+}=XV^{T}\left( VV^{T} \right)^{-1}-\boldsymbol{1}_{d}\left( \frac{\boldsymbol{1}_{d}^{T}XV^{T}\left( VV^{T} \right)^{-1}}{d}-\boldsymbol{1}_{r}^{T} \right)$,

where $\boldsymbol{1}_{d}$ and $\boldsymbol{1}_{r}$ are all-ones vectors.

The above updates are performed iteratively:

The “cosine” distance metric is utilized for assessing the similarities,

$$D_{ij}=cosine\left( C_{ix}, C_{xj} \right)=\frac{\sum_{x=1}^{n} C_{ix}C_{xj}}{\sqrt{\sum_{x=1}^{n} C_{ix}}\sqrt{\sum_{x=1}^{n} C_{xj}}}$$

where $C_{ij}$ is the raw contact of bin $i$ and bin $j$, $cosine$ is the cosine distance, $D_{ij}$ is the distance between bin $i$ and bin $j$. Lower distances indicate more similar connections, whereas higher distances reflect weak or repelled contacts between bins. Maps exhibiting pronounced checkerboard patterns show a bimodal distribution of similarity values (high and low; Figure S24), resulting in a high information entropy,

To investigate the correlation between polymorphism with chromosome length range, we conducted the Spearson correlation between polymorphic scores with chromosome length difference ratio, calculated by ${(max\left( chrolength \right)-min(chrolength))}/{max\left( chrolength \right)}$.

### Supplementary figures

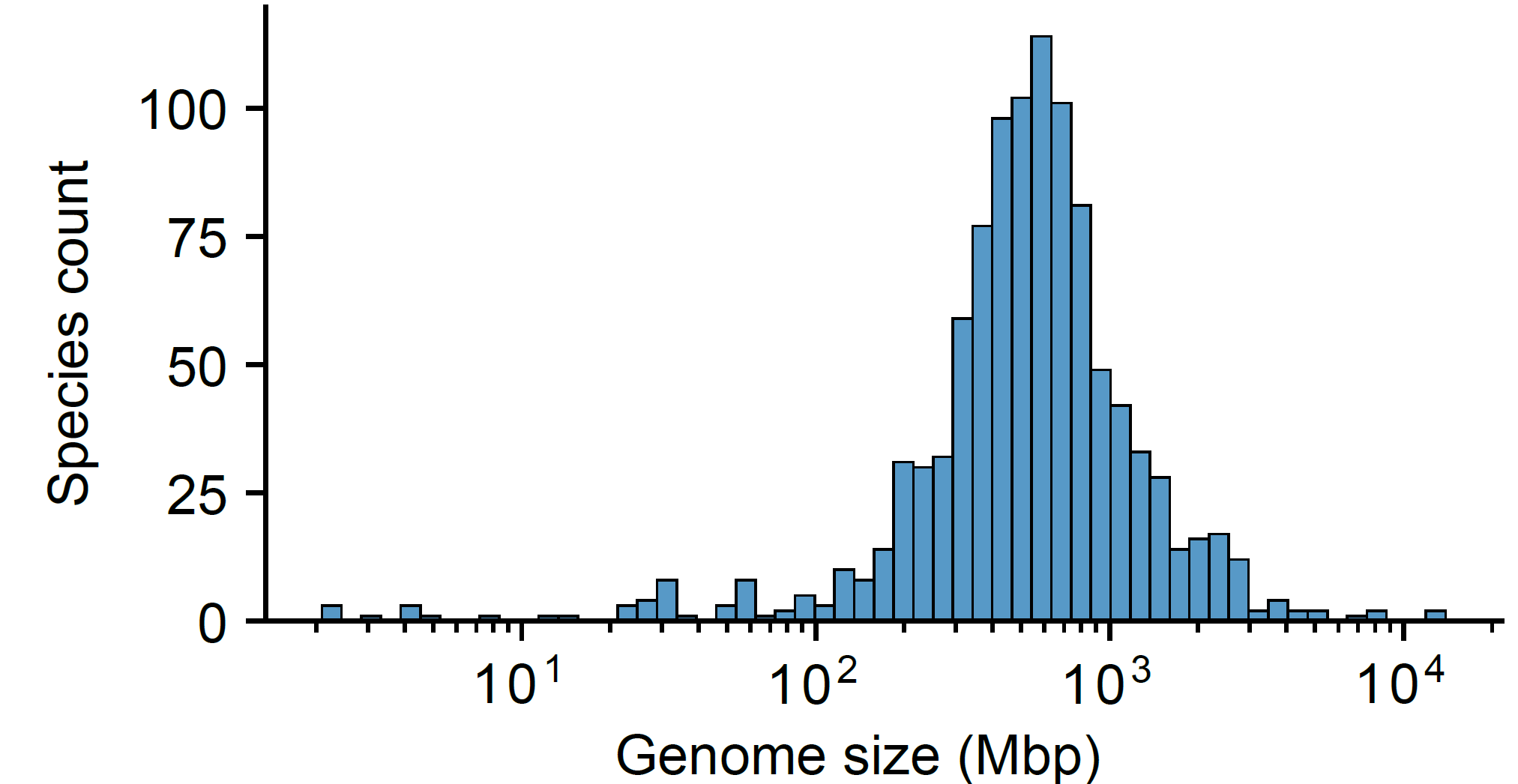

#### Figure S1 Variation in Genome Sizes

Genome sizes are enormously varied among analyzed 1,025 species, from 2.1 Mb in *Thermococcus kodakarensis* to 14.6 Gb in *Triticum aestivum*.

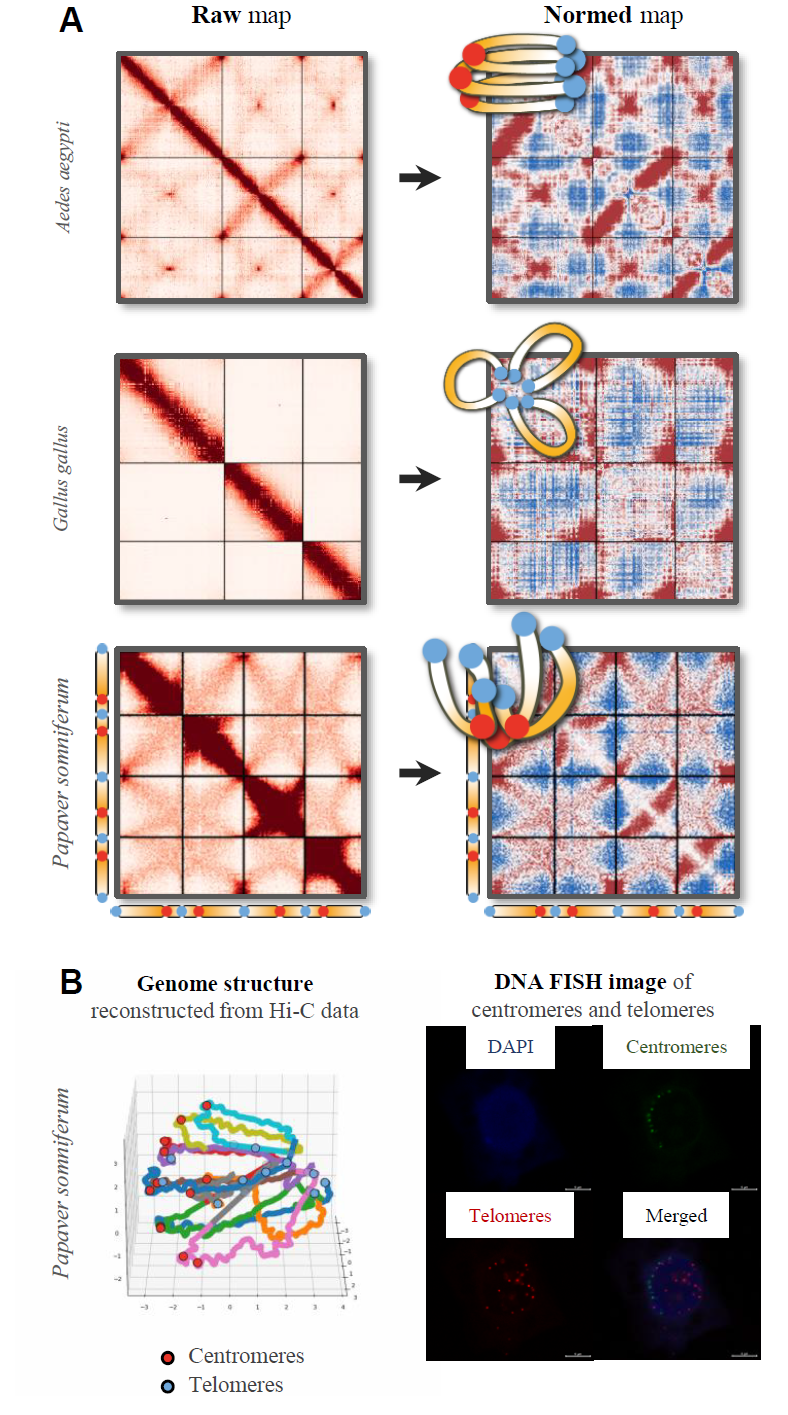

#### Figure S2 Examples and Integrative Validation of NormDis

A. Raw (left) and NormDis-normalized (right) Hi-C maps of three example species, with their exposed large-scale structures illustrated at the upper left corner. Normalization enhances long-distance interactions and suppresses short-distance interactions.
B. NormDis-predicted architecture of *Papaver somniferum*, reconstructed genome structure by Pastis^51^ with centromeres and telomeres labeled by colored dots, and DNA FISH images with centromeres and telomeres labeled. The concordance between NormDis predictions, polymer models, and FISH data demonstrates robust cross-validation of genome architecture inference methods in non-model species.

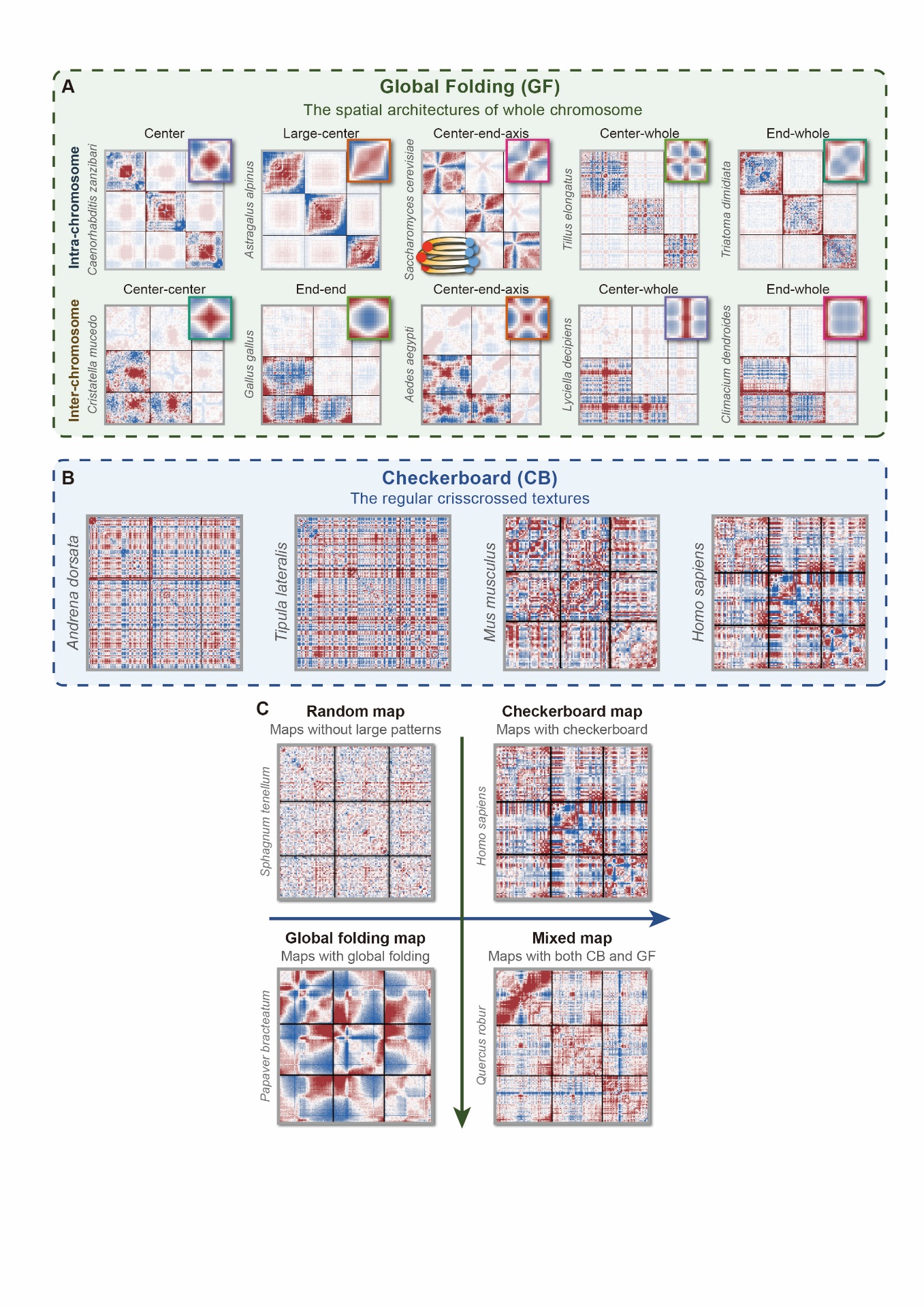

#### Figure S3 Diversity and Coexistence of High-Order Genome Architectures.

A. Spectrum of global folding patterns. Global folding reflects karyotype-scale spatial organization of chromosomes. Unsupervised clustering of Hi-C maps from 1,025 species identified five intra- and five inter-chromosomal types. Representative species for each type are shown, with each displaying its first three chromosomes. The average pattern for each type is plotted in the upper right corners. Genome configuration of yeast (*Saccharomyces cerevisiae*) is illustrated as an example for global folding.
B. Checkerboard patterns are the regular crisscrossed textures on maps, which indicate chromatin compartments. Four species with strong checkerboard organization are displayed, with each displaying its first three chromosomes.
C. Architectural independence and coexistence. Maps without any high-order architectures (named as random maps), maps with only checkerboard or global folding patterns, and maps with both global folding and checkerboard patterns are shown, displaying the first three chromosomes of each.

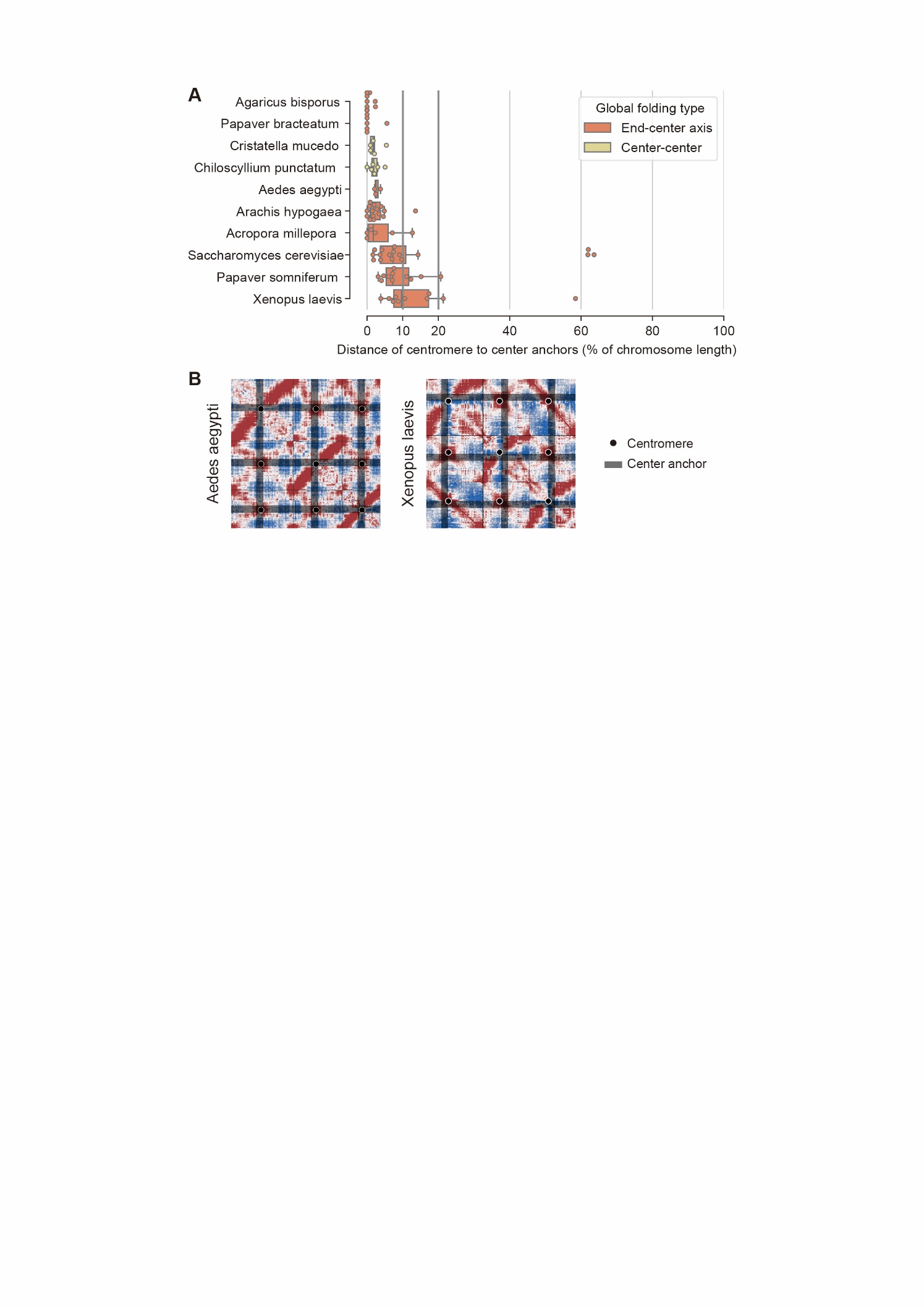

#### Figure S4 Coupling of Center Anchors and Centromeres.

A. Center anchors locate near centromeres, as indicated by the ratio of the distance between center anchors and centromeres corresponds to chromosome length, analyzed in species exhibiting center-associated inter-chromosomal global folding (end-center-axis and center-center types).
B. Positions of centromeres and center anchors on two example species, displaying the first three chromosomes of each. Centromeres are marked by the black dots, and the positions of center anchors are marked by the black lines.

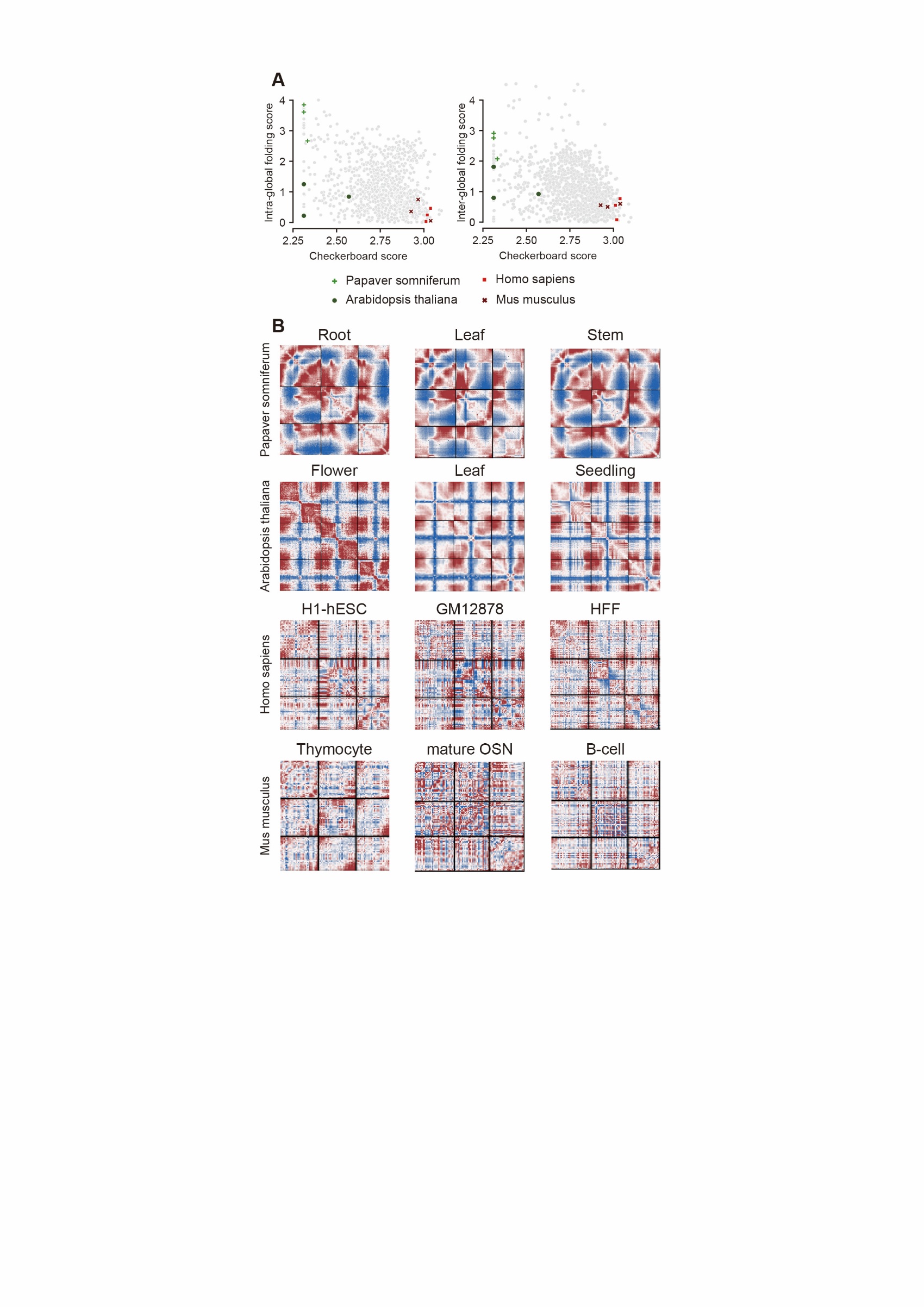

#### Figure S5 Conservation of high-order genome architectures across mature tissues

A. Architectural conservation across tissues. Scatter plot of global folding scores (GFS) and checkerboard scores (CBS) across tissues from four species, with 1,025 species as background. Detailed scores are documented in Supplementary Table 5.
B. Normalized contact maps for exemplified tissues across four species, showing conserved global folding and checkerboard patterns within species. Black lines demarcate chromosome boundaries.

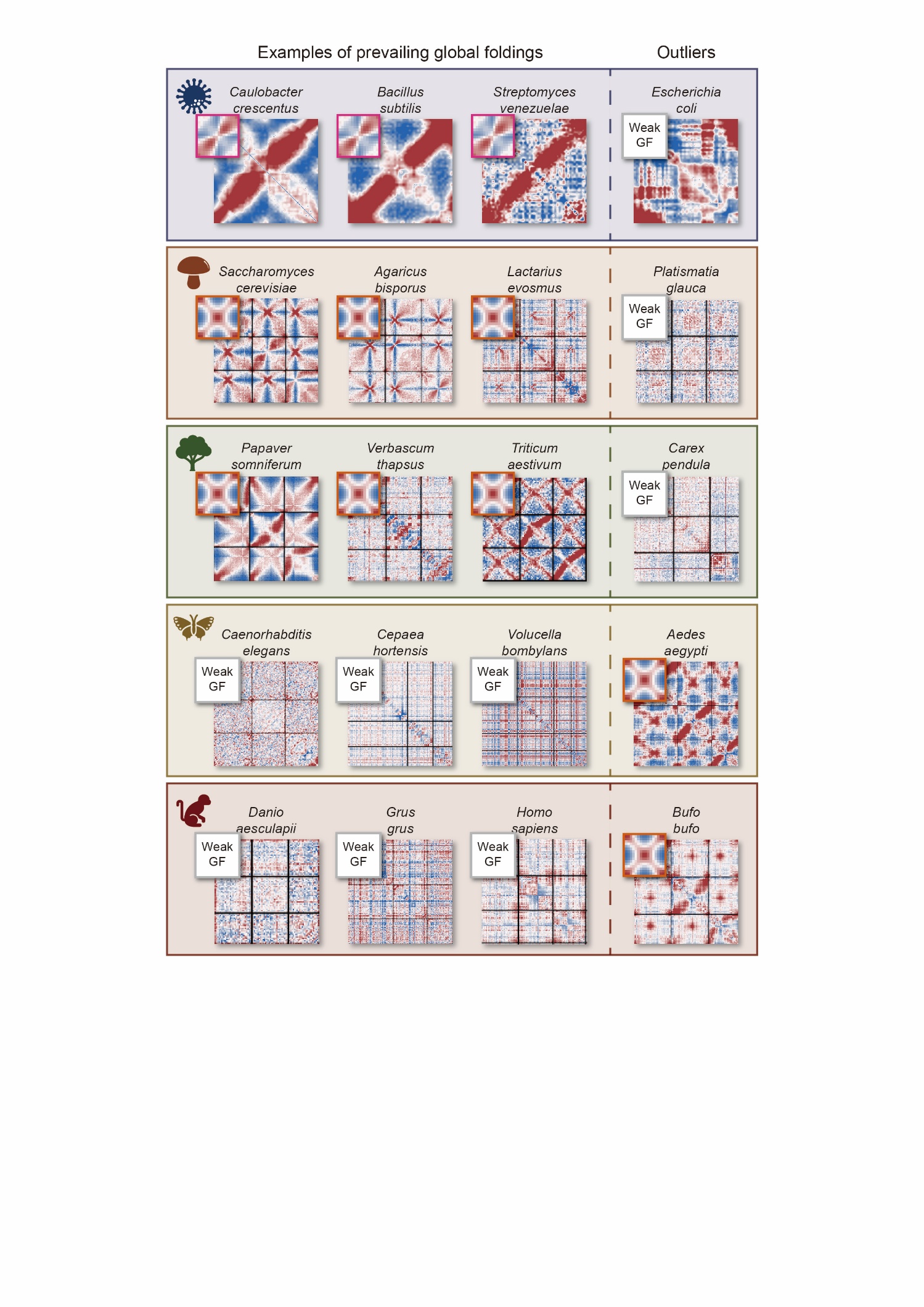

#### Figure S6 Taxonomic Prevalence and Exceptions in Global Folding.

Left three panels: taxonomically dominant patterns, examples of global folding types within major taxonomic groups. Right panel: outlier species exhibiting non-canonical folding types.

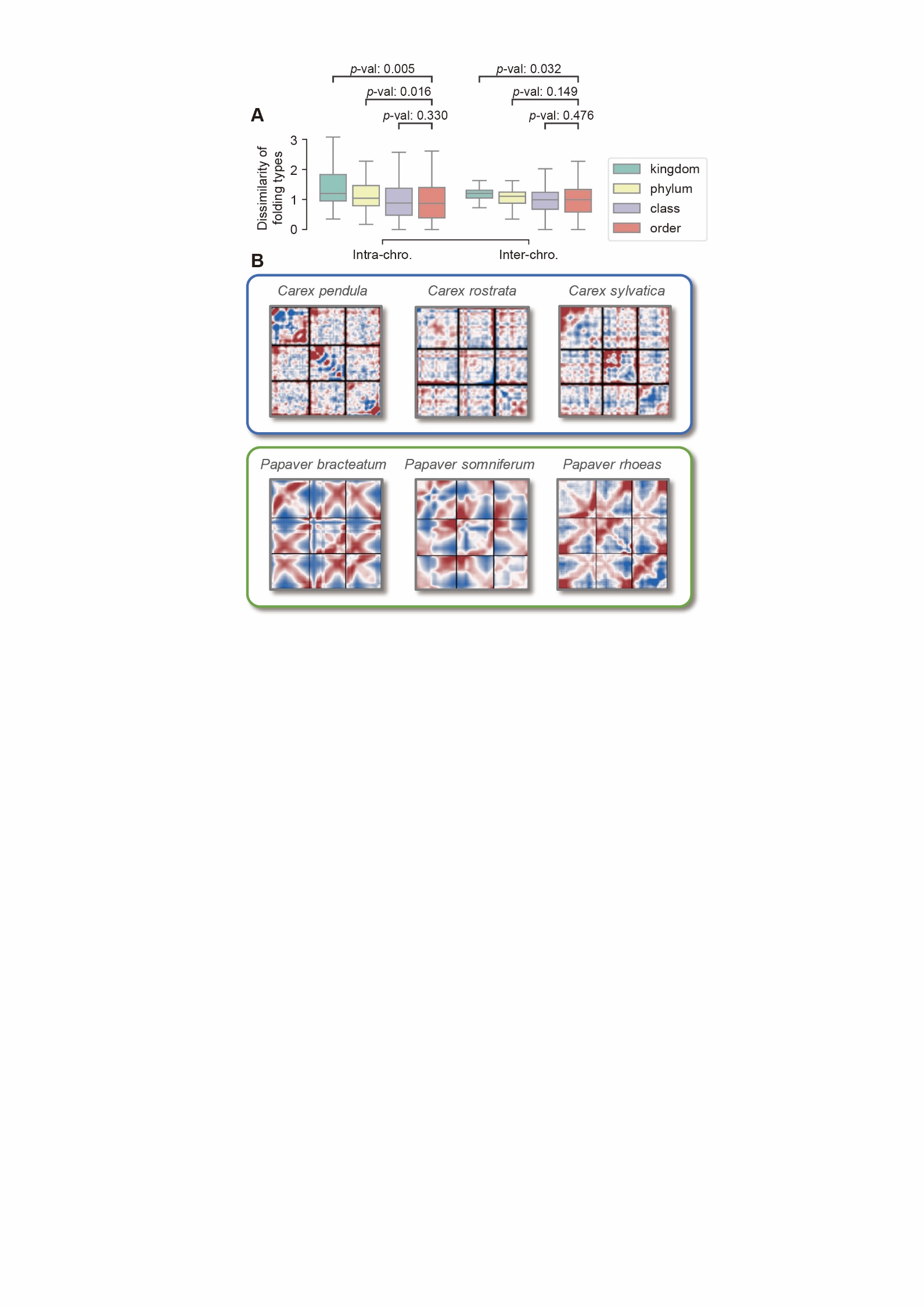

#### Figure S7 Phylogenetic Conservation and Divergence in Global Folding.

A. Architectural variation is restricted by phylogenetic relatedness. Standard deviations (SDs) of global folding scores decrease with finer taxonomic resolution. *P*-values are calculated by the Mann-Whitney U test with a greater alternative hypothesis, which includes 3 kingdoms, 11 phyla, 15 classes, and 33 orders. The reduced sample sizes resulting from finer taxonomic divisions likely explain the non-significant differences between closely related taxonomic divisions (for example between class and order).
B. Exemplified lineage-specific global folding. Two lineages from the same kingdom (plants) exhibit distinct architectures. The upper species from *Carex* genus lacks strong global folding patterns, while the lower species from *Papaver* genus exhibit center-end-axis patterns.

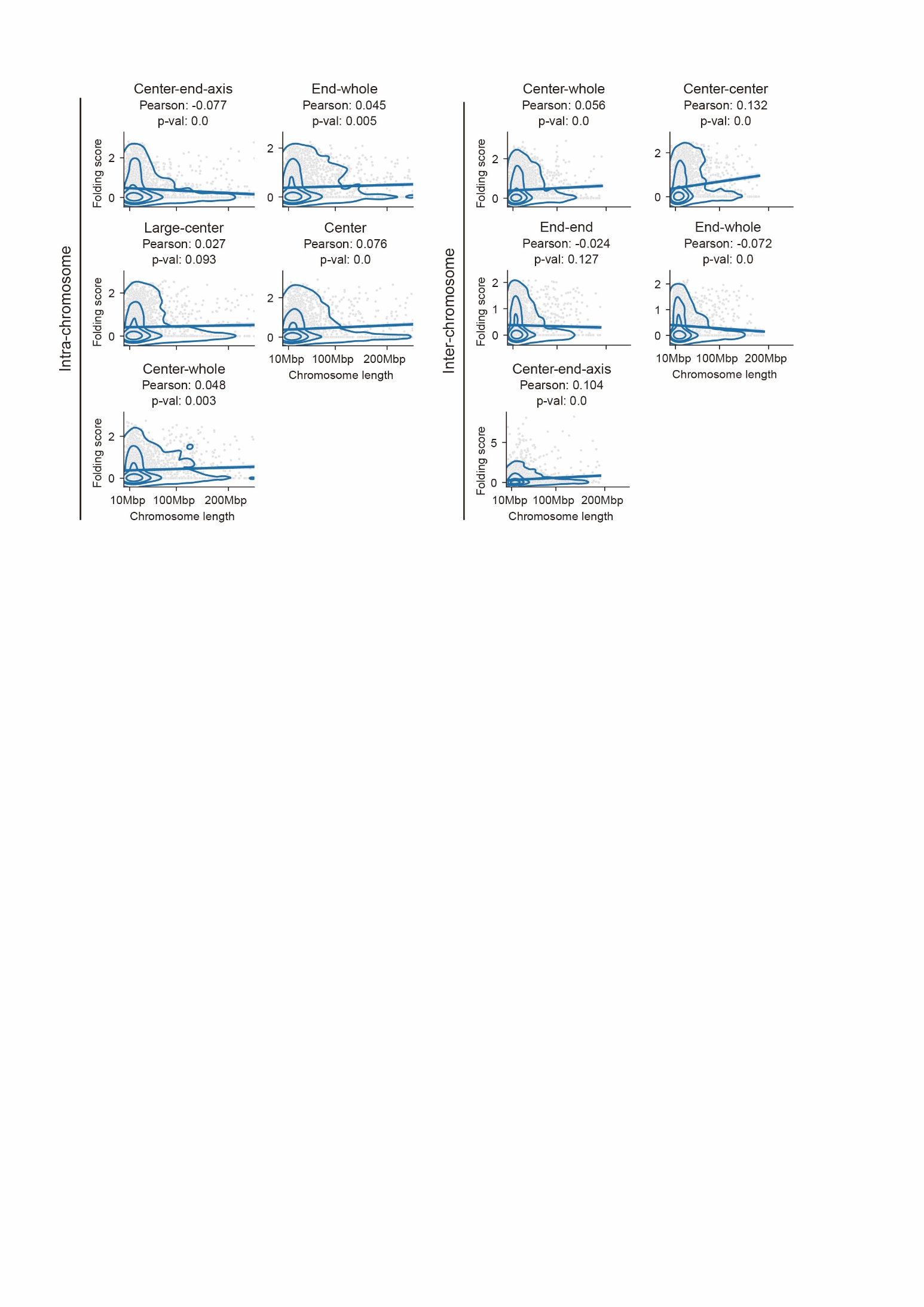

#### Figure S8 Global Folding Is Uncoupled from Chromosomal Morphology.

Pearson correlation analysis reveals no significant association (|r| < 0.13 for all folding types) between global folding scores and chromosomal length. 20,000 randomly sampled chromosomes are shown for each correlation. Results are stratified by five intra- (top row) and five inter-chromosomal (bottom row) global folding types.

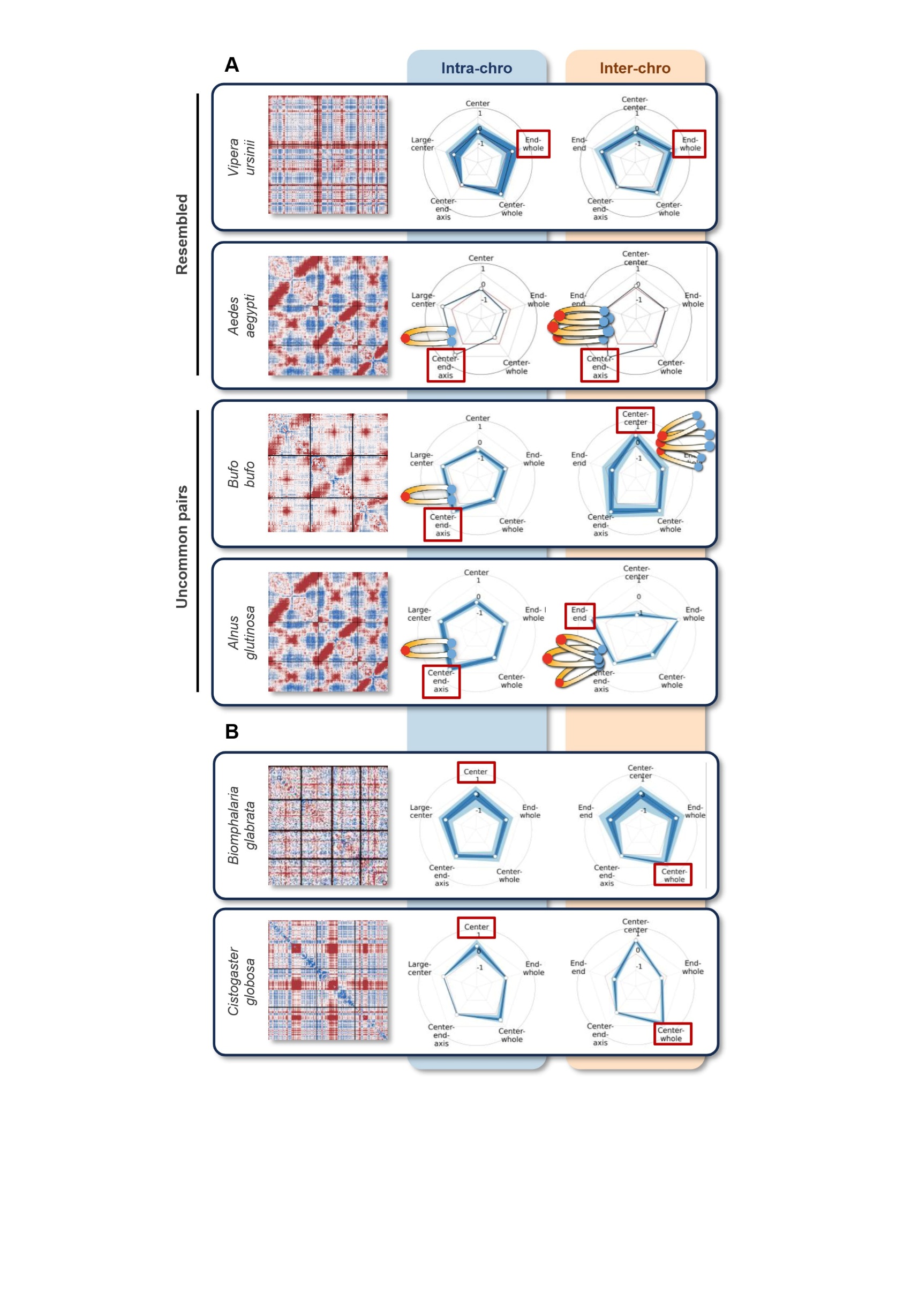

#### Figure S9 Variety in global folding type interplay.

A. Variety in the interplay between intra- and inter-global folding types. The upper two panels exhibit species with resembled global folding pairs. The lower two panels exemplify the uncommon pairs. Normalized maps are shown on left for each species, displaying the first three chromosomes. Rader maps exhibiting the intra- and inter-global folding type strength are shown on right. Illustrations for center-end-axis related types are displayed. The strongest types are marked by the red rectangles.

Although the three illustrated species exhibit the same intra-global folding type, center-end axis, the inter-global folding types differs. *Aedes aegypti* displays the typical parallel arrangements of chromosomes, whiles *Bufo bufo* exhibits stronger center anchors clustering but weak end anchors clustering, and *Alnus glutinosa* exhibits the opposite. This indicates the variety in the interplay between intra- and inter-global folding type.

B. Example species of different global folding types but sharing the same anchor, displaying the first three or four chromosomes of each. The number of species belonging to each pair is shown in Figure S8. The strongest types are marked by the red rectangles.

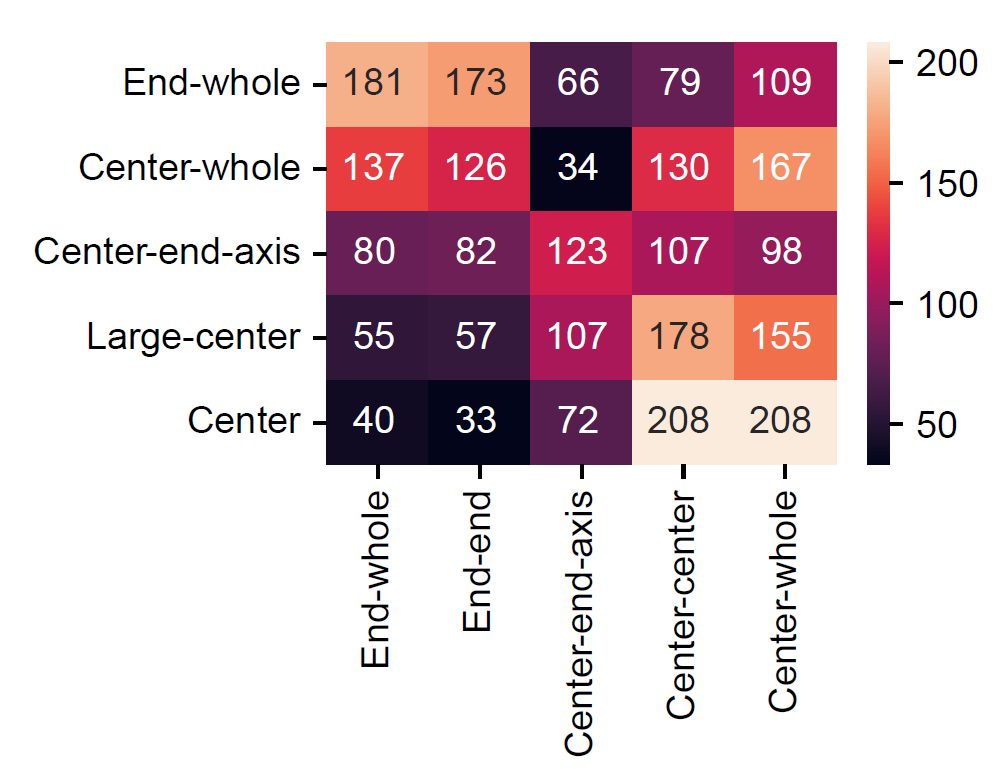

#### Figure S10 Variety in global folding type interplay (pair number).

Analysis of 1,020 species reveals widespread diversity in global folding pairs. Rows: five intra-chromosomal folding types; columns: five inter-chromosomal folding types. Numbers represent species counts for each pairwise combination.

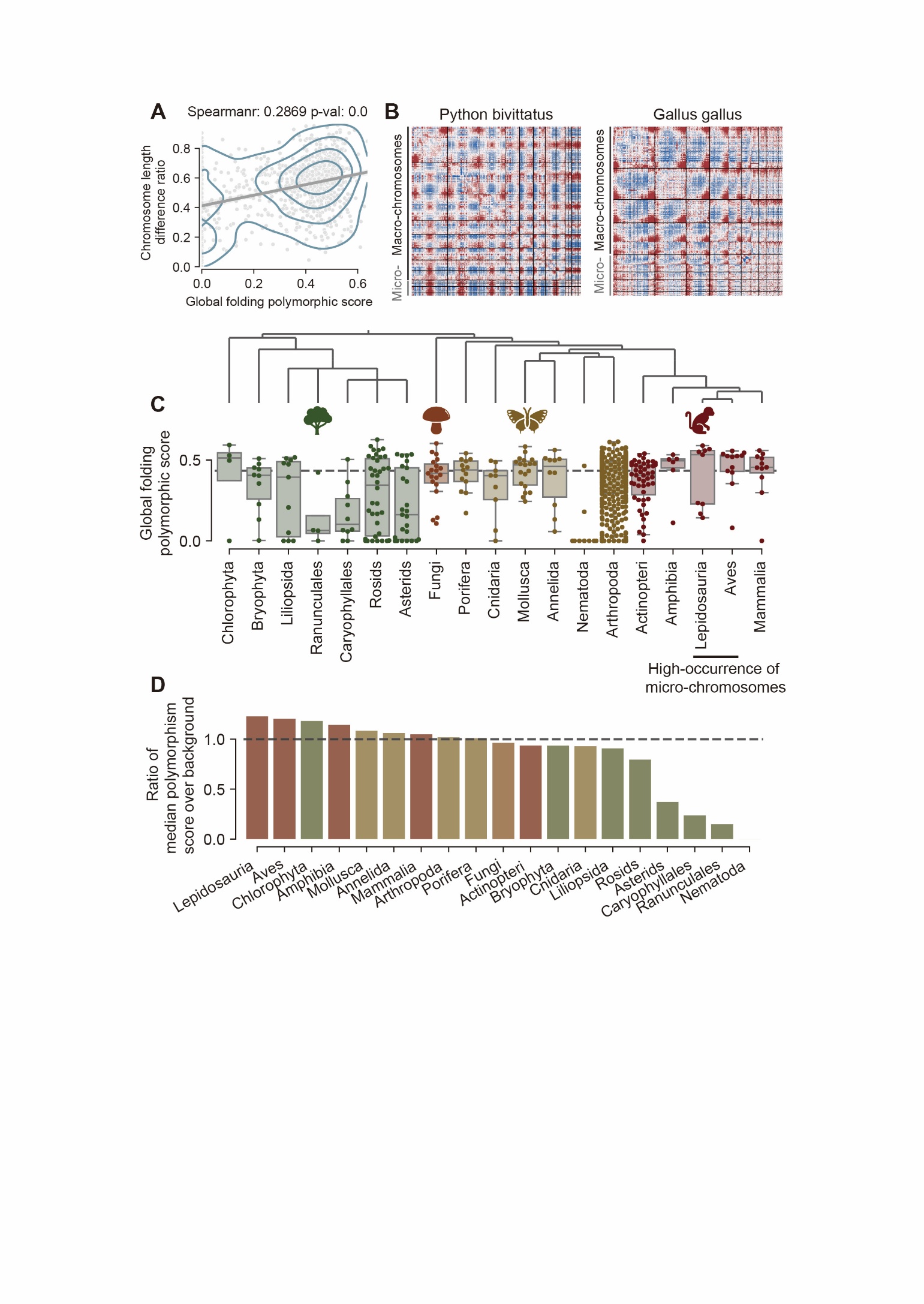

#### Figure S11 Prevalence of Global Folding Polymorphism.

A. Chromosomal length variation associates with folding polymorphism, indicated by the correlation between global folding polymorphic scores and chromosome length heterogeneity across 1,025 species.
B. Representative species with global folding polymorphism in addition to Figure 2D.
C. Global folding polymorphic score of main taxonomical groups, indicating the broad distribution of polymorphism.
D. Sorted ratios of median polymorphic score over background (marked by the black line). Detailed data is shown in Supplementary File 5.

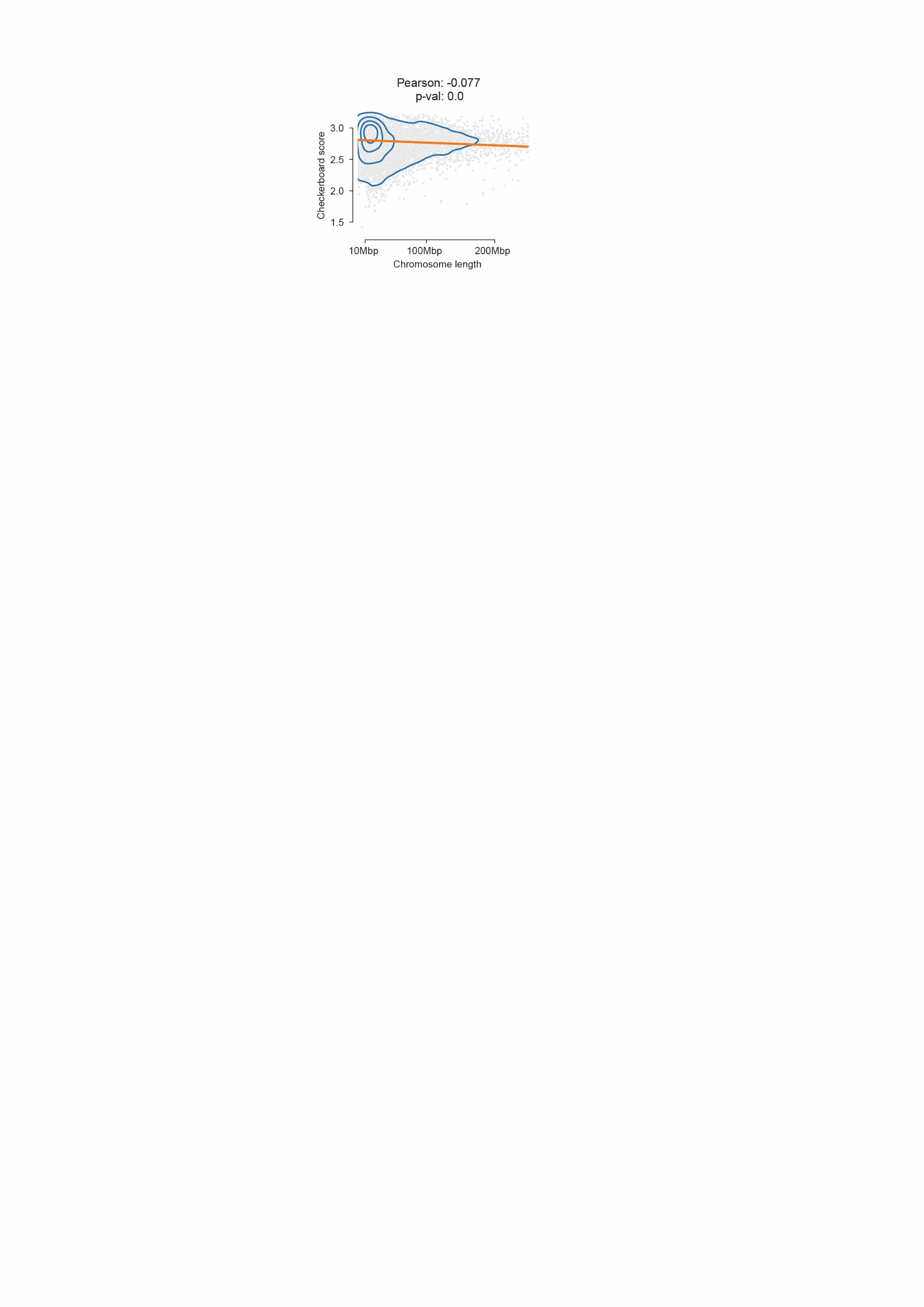

#### Figure S12 Checkerboard Architecture Is Uncoupled from Chromosomal Morphology.

Correlations between chromosomal length and checkerboard score (r = -0.077). 20,000 chromosomes are randomly sampled for efficient plotting.

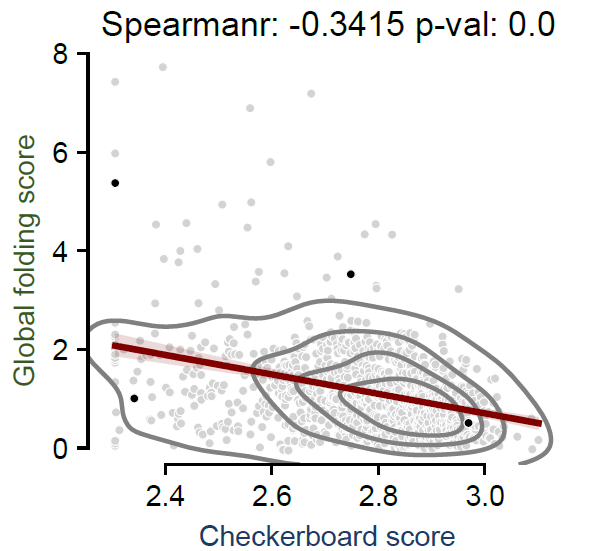

#### Figure S13 Negative correlation between checkerboard and inter-chromosomal folding.

Checkerboard scores exhibit a negative correlation with inter-chromosomal global folding scores across 1,025 species (Spearman ρ = −0.3415, p < 1×10−15). Correlation with intra-global folding score is shown in Figure 5A. The four example species shown in Figure 5A are marked by the black dots in this plot. This suggests the competing relationship between checkerboard and global folding.

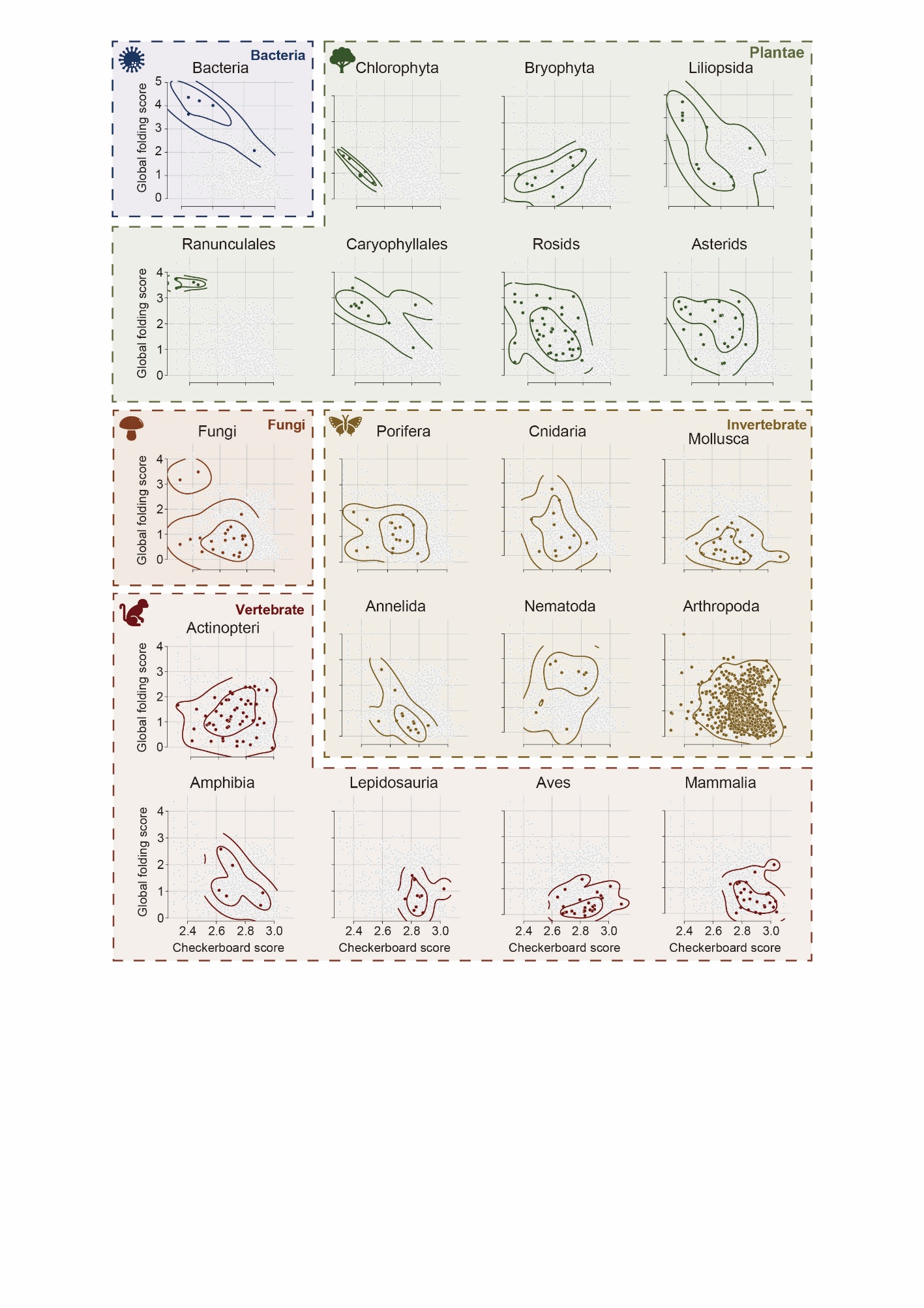

#### Figure S14 The evolutionary trajectory of 3D genome as shown by checkerboard score and intra-chromosomal global folding scores.

Scatter plot of checkerboard scores versus intra-global folding scores for 21 taxonomic groups (group sizes shown in Fig. 1A), overlaid on the background distribution of 1,025 species (gray points).

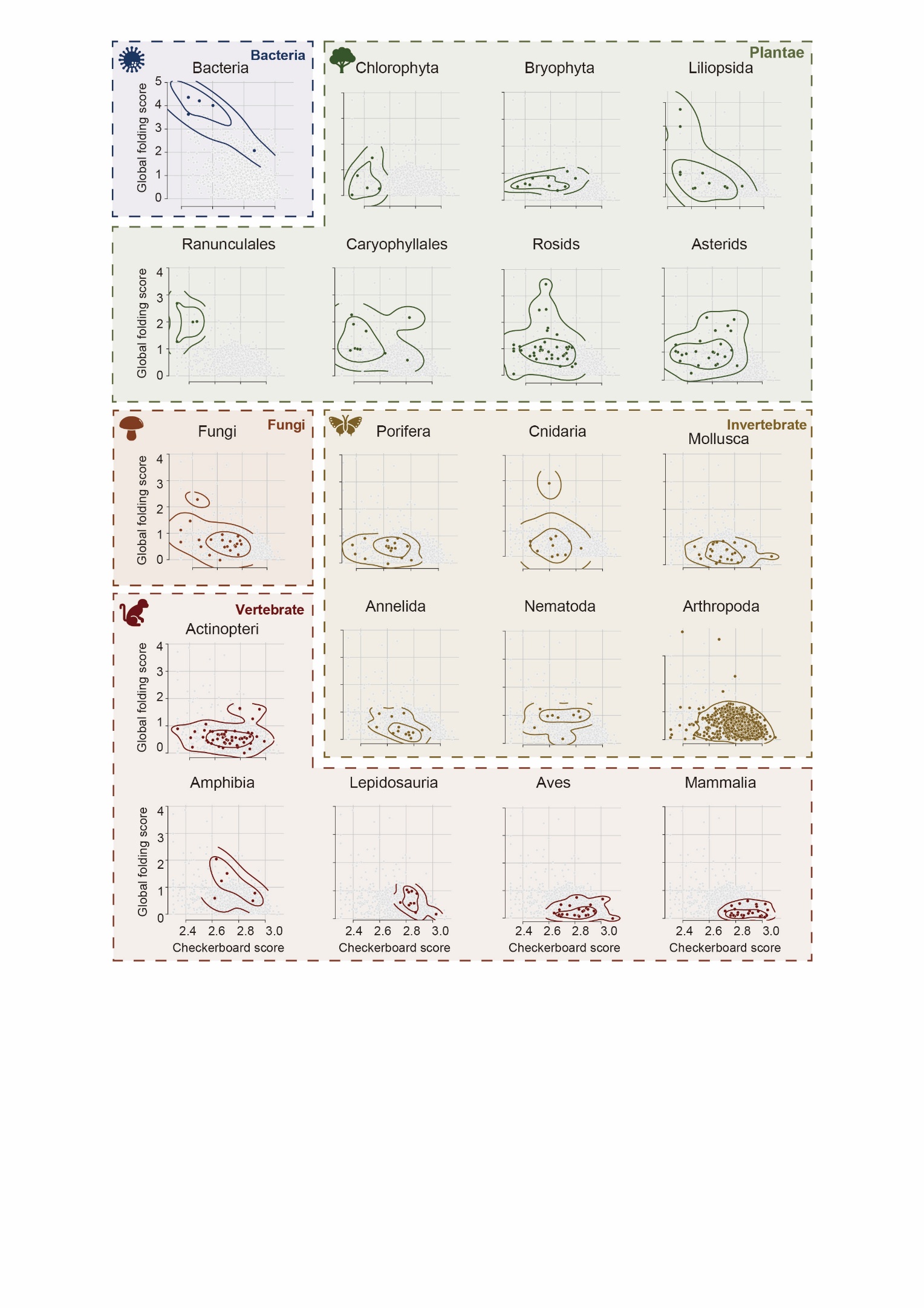

#### Figure S15 The evolutionary trajectory of 3D genome as shown by checkerboard score and inter-chromosomal global folding scores.

Scatter plot of checkerboard scores versus inter-global folding scores for 21 taxonomic groups (group sizes shown in Fig. 1A), overlaid on the background distribution of 1,025 species (gray points).

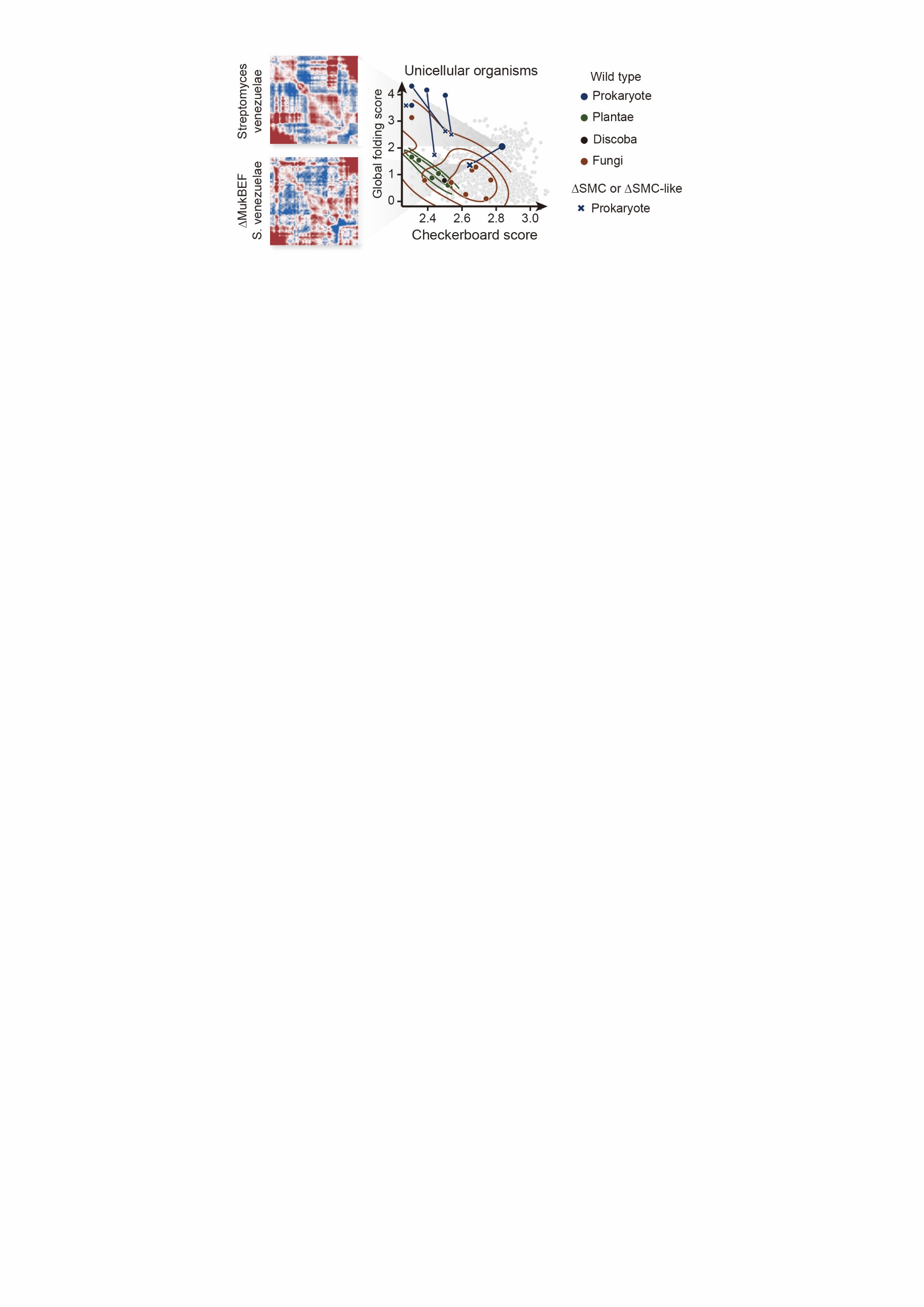

#### Figure S16 MukBEF Drives Checkerboard Patterns in *E. coli*.

Normalized Hi-C maps of *Escherichia coli* (wild-type) and its MukBEF knockout mutant. Wild-type exhibits the interaction stripes reflecting spatial compartmentalization, while the mutant shows weaker patterns.

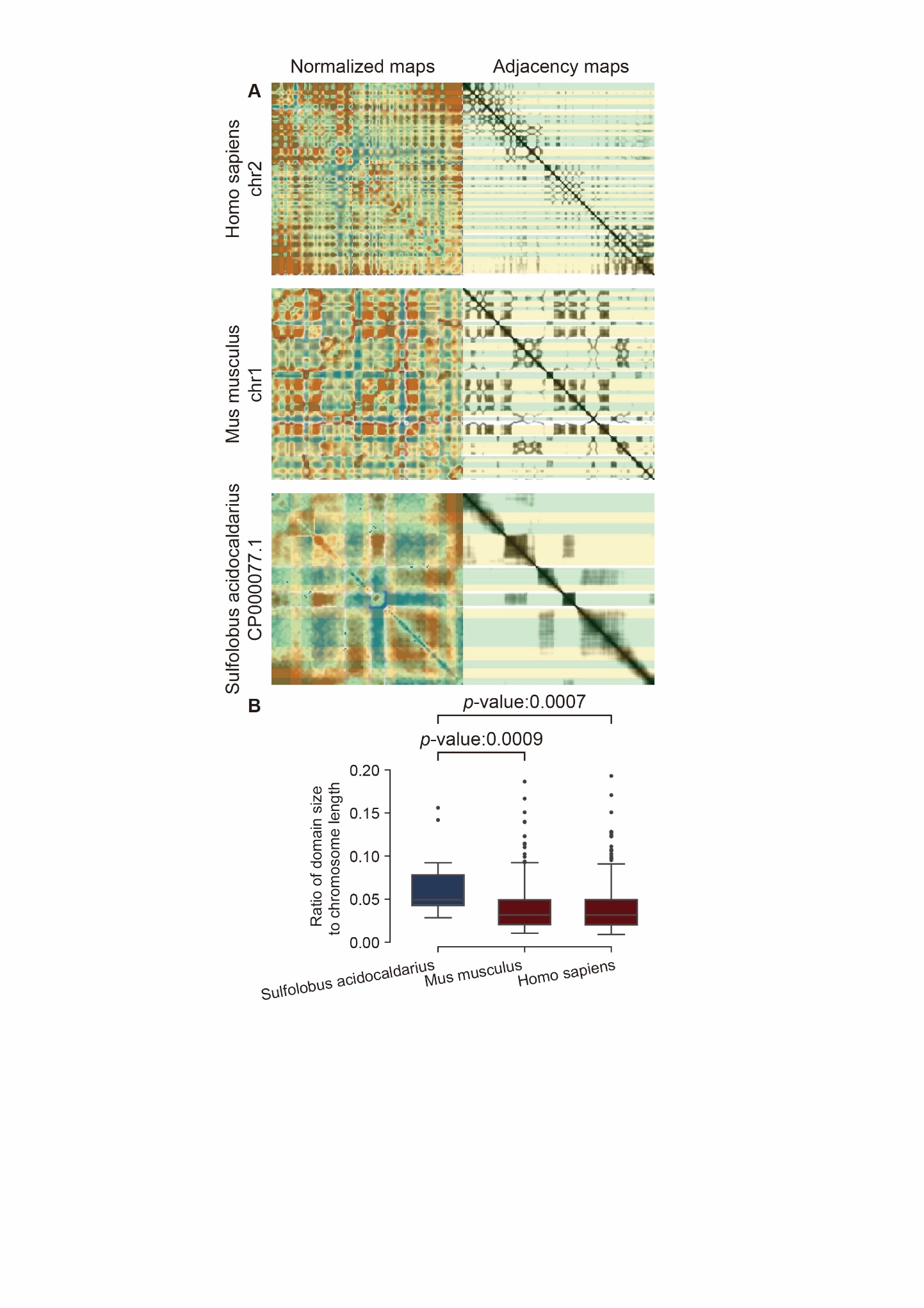

#### Figure S17 Checkerboard Domain Sizes among Species.

A. Comparative visualization of checkerboard domain sizes. Normalized Hi-C maps (left) and adjacency matrices (right; by cosine distance) from *Homo sapiens* (Chr. 2), *Mus musculus* (Chr. 1), and *Streptomyces venezuelae*. Compartment domains are marked by the yellow or green boxes.
B. Quantitative scaling of compartment size. Distribution of compartment-to-chromosome length ratios for human, mouse and *S. venezuelae*, as measured by their ratio to chromosome length. 13 domains are found in *S. venezuelae* (median ratio 4.96%), 385 in mice (3.17%; Mann Whitney U test *p*-value: 0.0009) and 415 in human (3.14%; 0.0007).

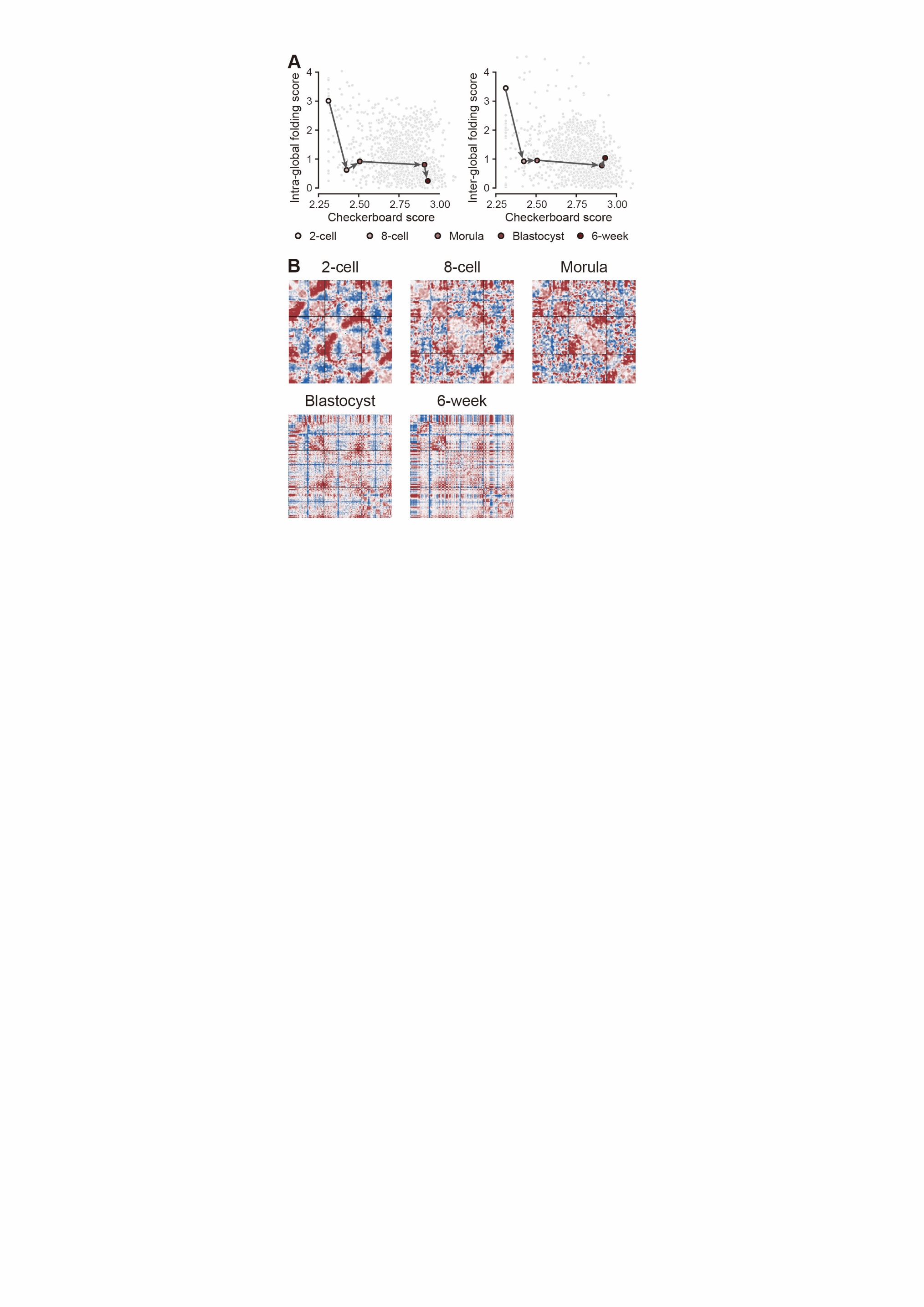

#### Figure S18 Architectural Transition During Human Embryogenesis

A programmed transition of high-order architectures is observed during human embryogenesis. It indicates that global folding maintains genome plasticity for totipotency, while checkerboard architecture enables cell fate determination through spatially coordinated gene regulation.

A. Developmental trajectory of 3D genome reorganization. Scatter plots trace the progression from global folding to checkerboard patterns during early human embryogenesis. The developmental order is indicated by the arrows, with each stage is labeled below.
B. Normalized maps of human early embryo development, displaying the first three chromosomes of each.

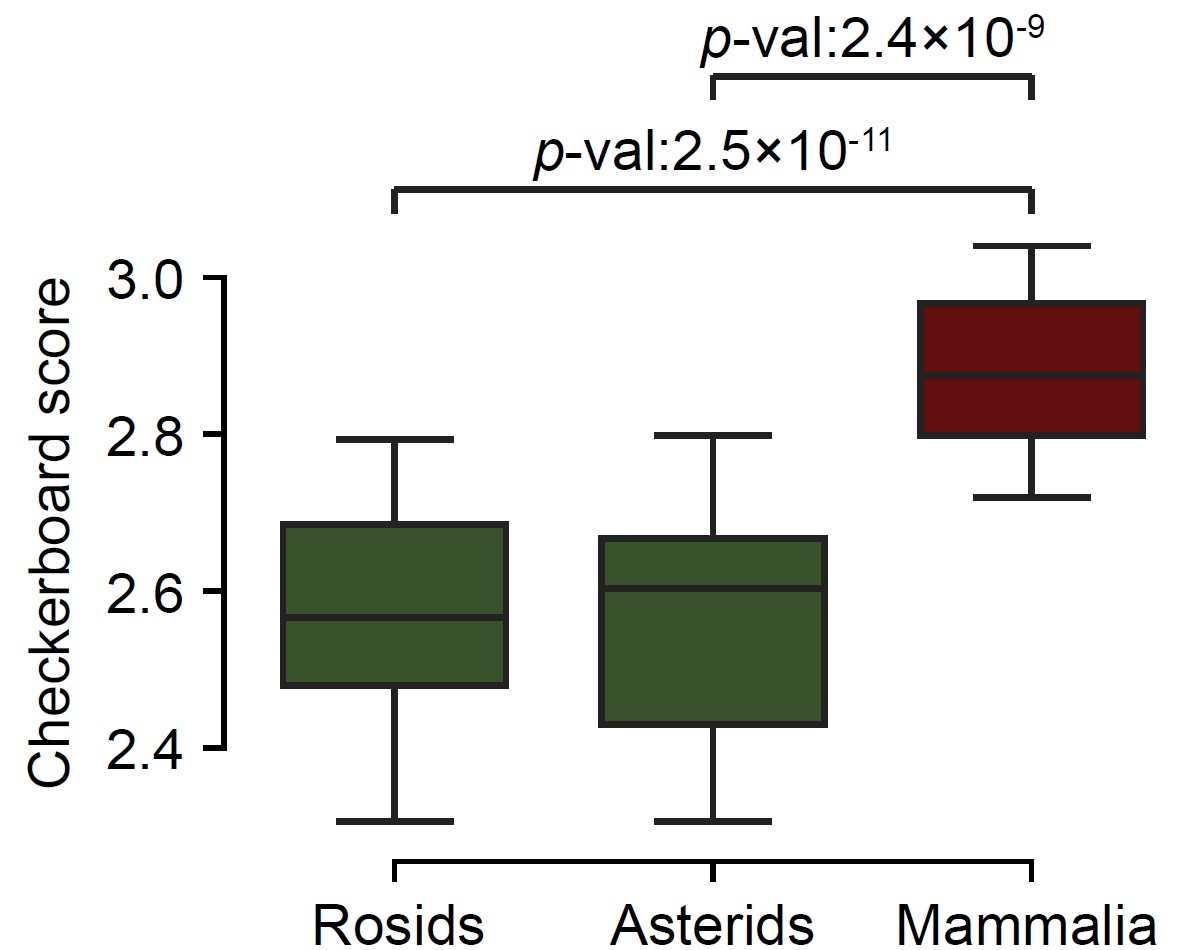

#### Figure S19 Checkerboard Strength across Plant and Animal Lineages

Boxplot of checkerboard scores between Rosids (n=39), Asterids (n=24) and Mammalia (n=23). *P*-value is calculated by Mann-whitney U test. Checkerboard scores differ significantly between Mammalia (median CBS = 2.87) and eudicot clades (Rosids: 2.57, Asterids: 2.60).

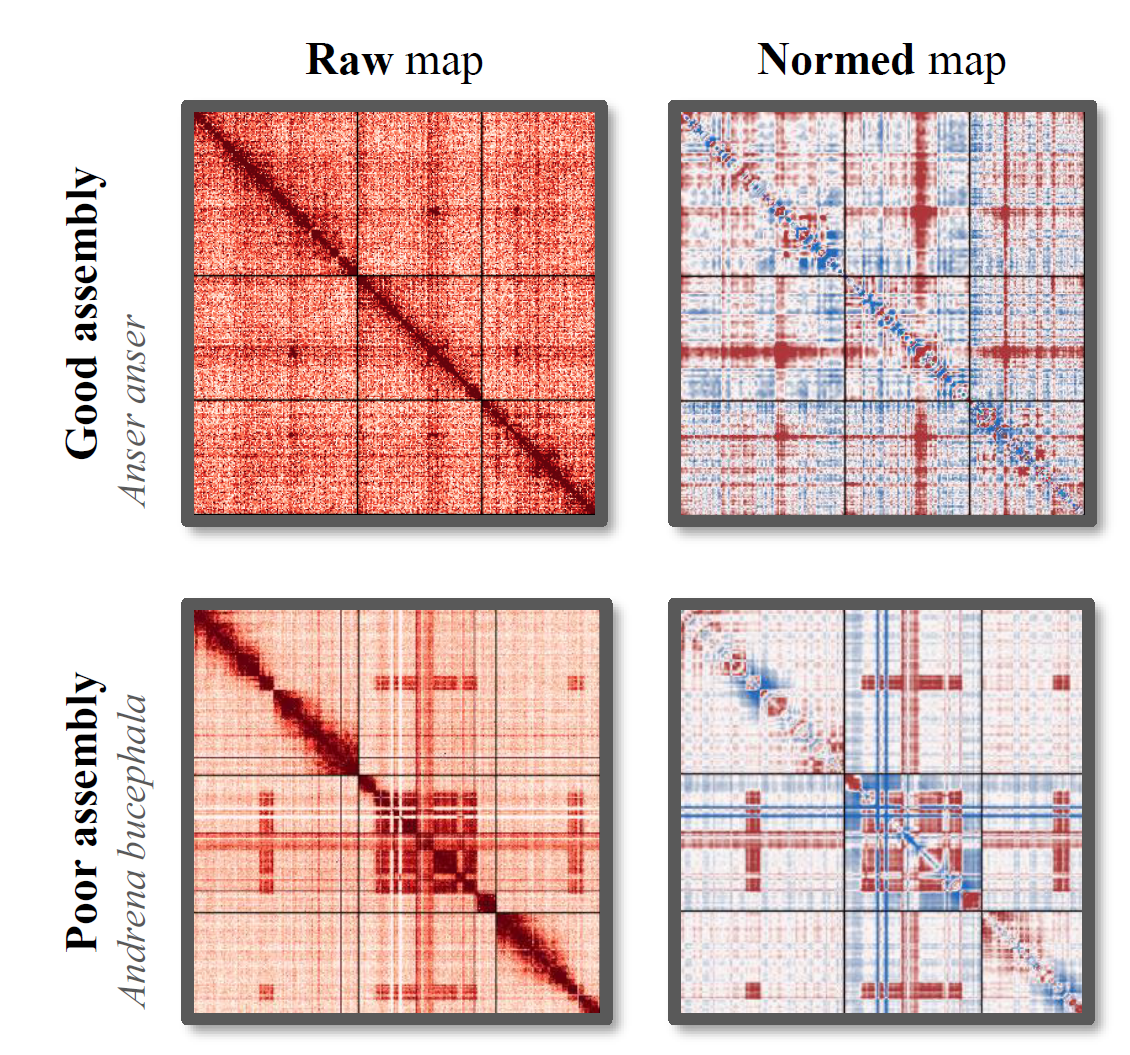

#### Figure S20 Assembly Quality Assessment Based on Hi-C

Top: Example species of high-quality assemblies. Species with center-whole global folding pattern exhibit gradual interaction decay from interaction-dense genomic bins without sharp transition.

Bottom: Example species of poor assemblies. Low-quality assemblies display sharp artifactual interaction spikes at erroneous scaffold junctions.

High-quality Hi-C assemblies are prerequisite for resolving true chromatin architectures, as poor assemblies introduce topological biases.

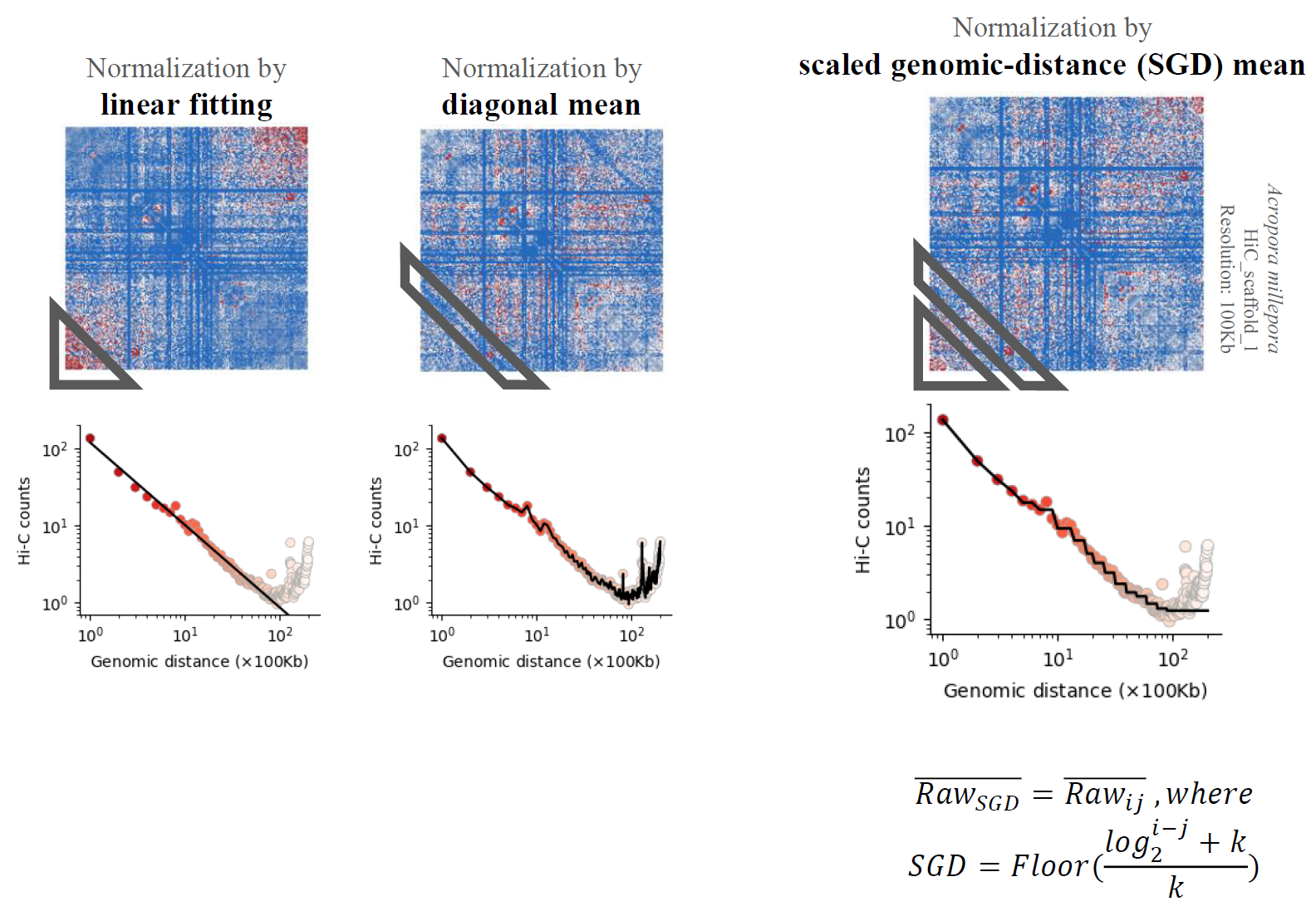

#### Figure S21 Comparative performance of Hi-C normalization strategies

Top: Normalized maps of *Acropora millepora* (Chromosome 1) processed by three normalization strategies.

Bottom: Interaction frequencies (Red dots) and expected frequencies (black lines). NormDis with SGD normalization strategy exhibit most accuracy in removing distance biases, avoiding overcorrection (linear fitting) or outlier susceptibility (diagonal mean).

1. Linear fitting: Artificially elevates long-distance interactions, marked by the gray triangle at the lower left corner.
2. Diagonal mean: Distorted by a long-distance outlier, marked by the gray quadrilateral.
3. NormDis: Corrects both biases, preserving true biological signals.

Detailed information is described in Methods.

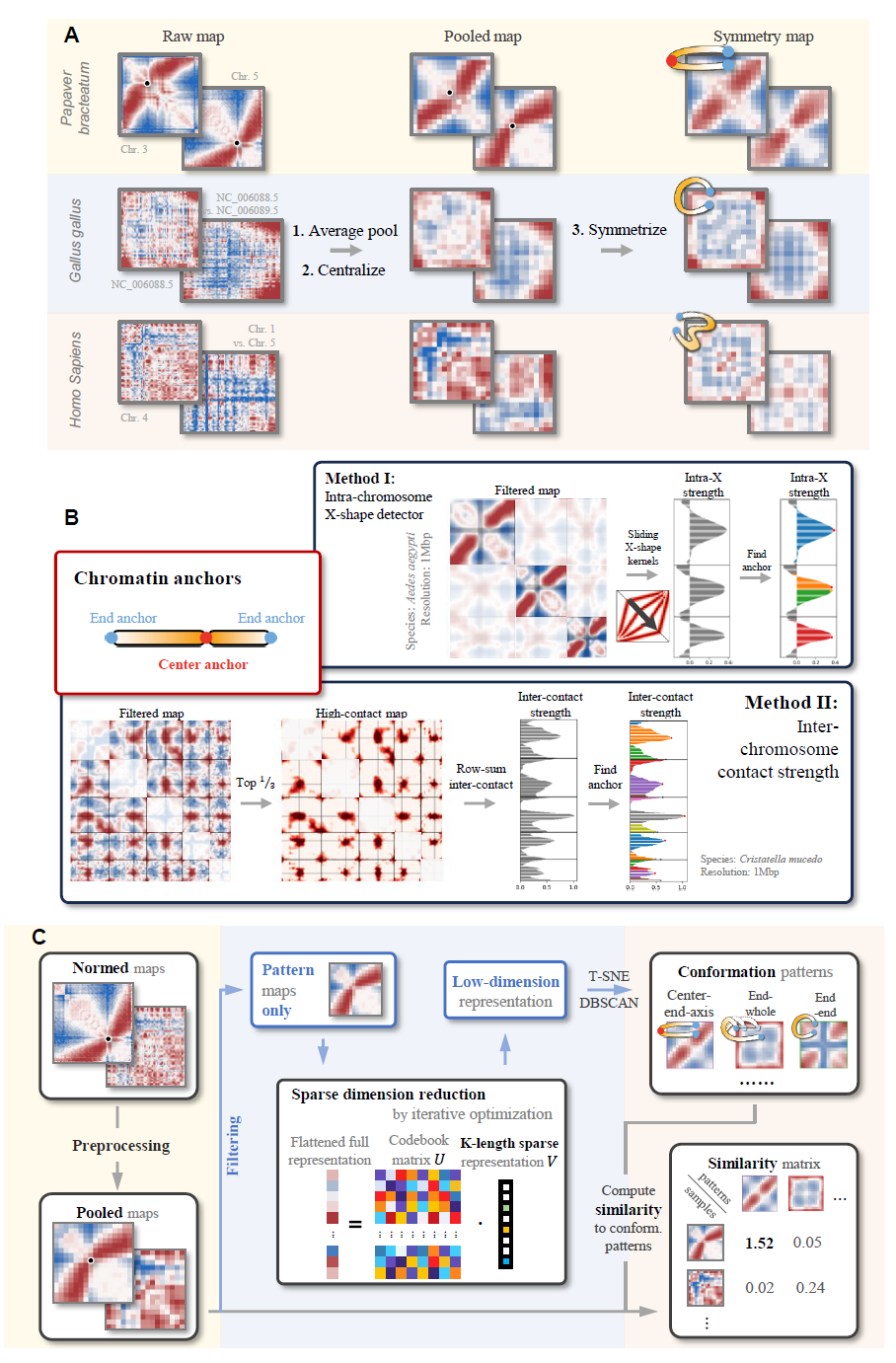

#### Figure S22 Systematic Pipeline for Global Folding Pattern Mining.

A. Preprocessing for large-scale pattern enhancement, including average pooling, centralization and symmetrization, detailed in Methods. Representative examples include *Papaver somniferum* (center-anchor), *Gallus gallus* (end-anchor) and *Homo sapiens* (no obvious global folding).
B. CenterFinder algorithm for center anchor detection. Dual-mode computational framework identifies folding centers: "Intra X" mode is optimized for center-end-axis patterns, while "Inter row sum" mode is optimized for anchors with elevated inter-chromosomal contacts.
C. Overview of global folding pattern mining. Four-stage analytical workflow includes training set curation, pre-processing, pattern mining and quantification of global folding.

This pipeline resolves global folding architectures, enabling cross-species comparison of large-scale genome organization.

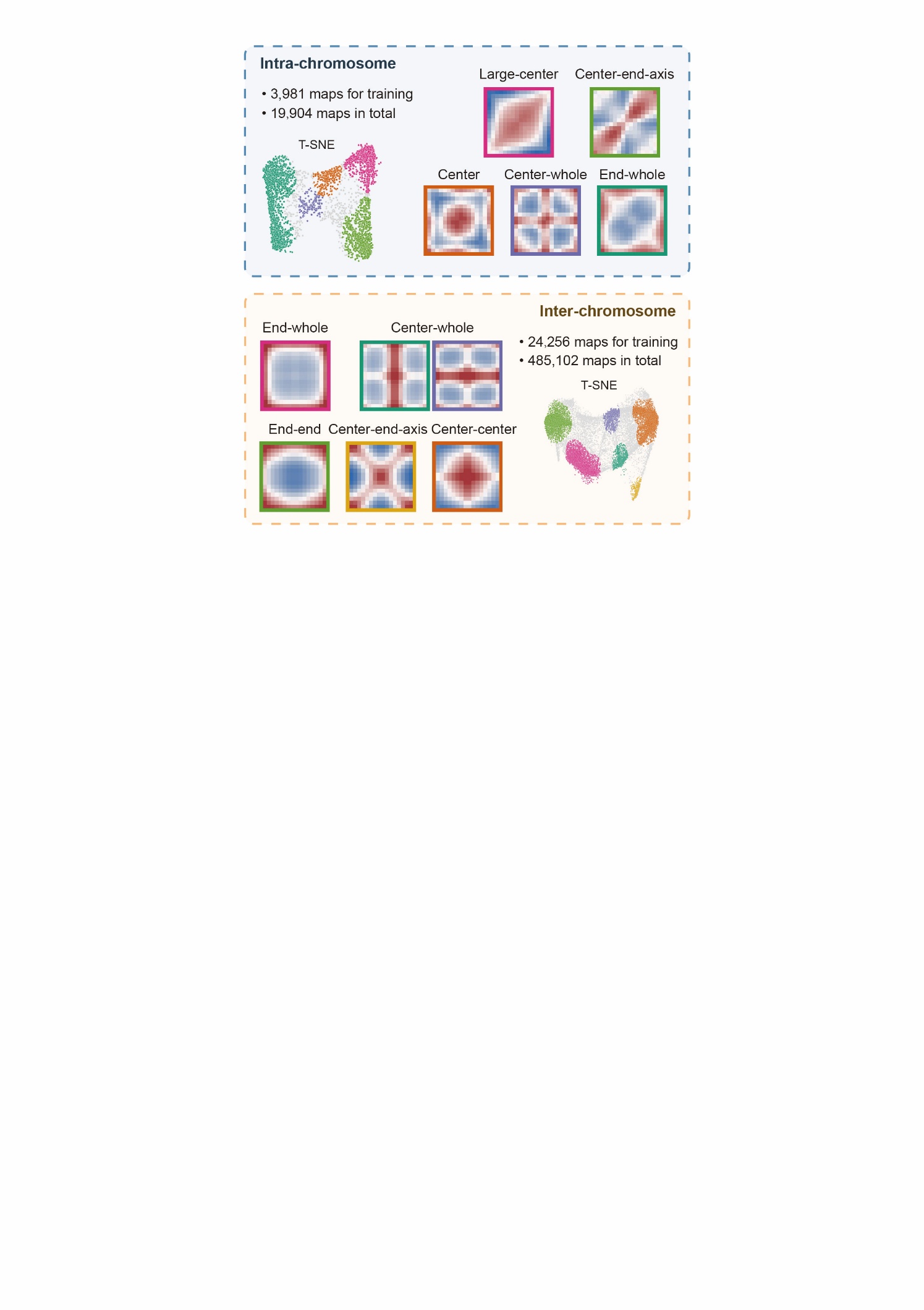

#### Figure S23 Robust Reproducibility of Global Folding Pattern Mining.

Global folding patterns were re-identified through independent runs with randomized dictionary matrix initialization and automated training set selection. The conserved recovery of global folding patterns across randomized runs validates our pipeline’s algorithmic stability and insensitivity to initial conditions.

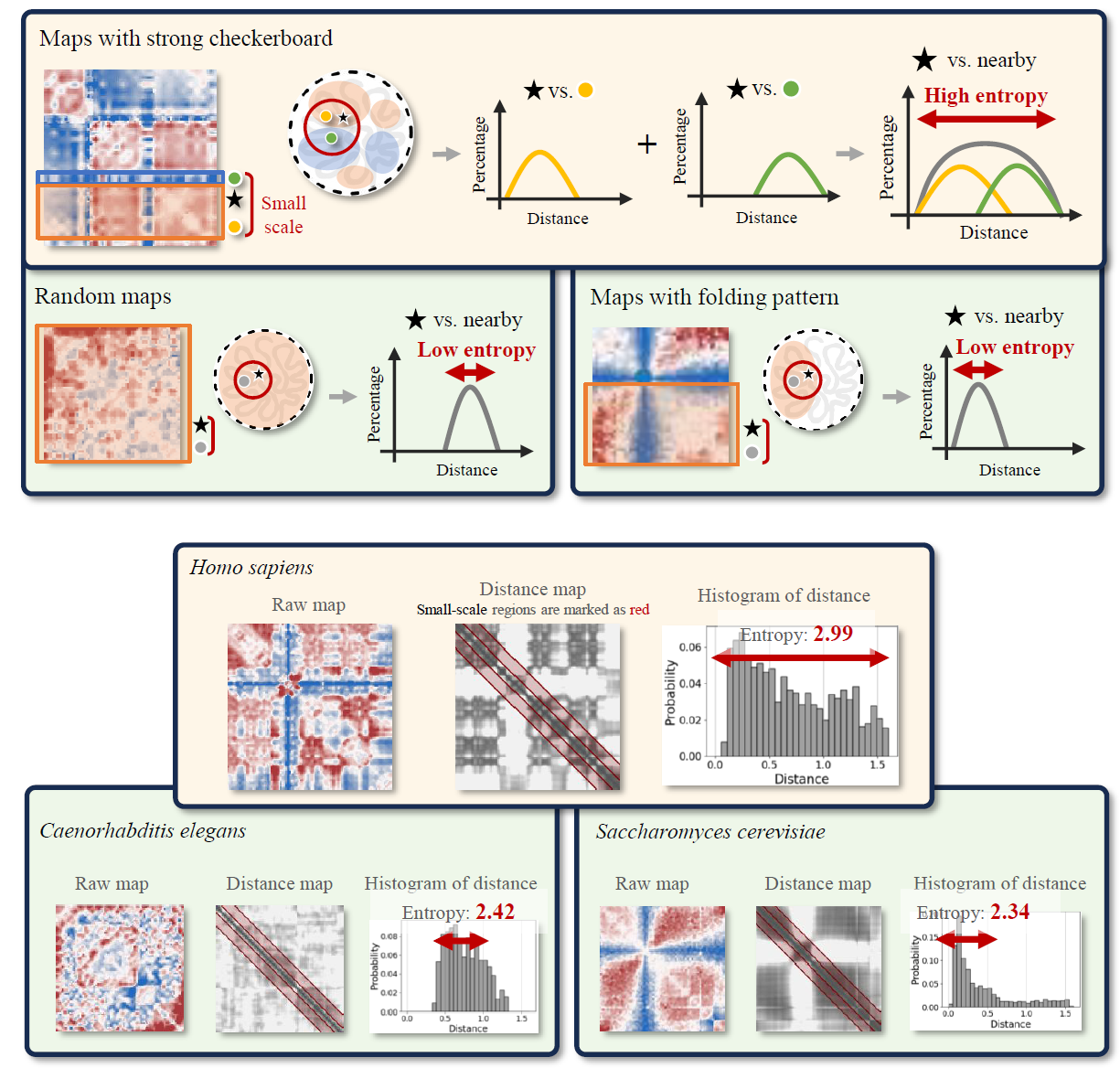

#### Figure S24 Quantitative Framework for Checkerboard Patterns

Illustrations for methods of quantifying checkerboard patterns.

The upper panel represents three chromatin organization paradigms: checkerboard-dominant, random and global folding-dominant. Illustrations of the underlying physical structures are shown in black edged circles, with colored circles denoting the compartments. The lower panel provides three example species, representing the three types of maps: checkerboard-dominant (*Homo sapiens*, high entropy 2.99), global folding-dominant (*Saccharomyces cerevisiae*, low entropy 2.34) and random (*Caenorhabditis elegans*, low entropy 2.42). This entropy-based approach objectively quantifies compartmentalization strength.

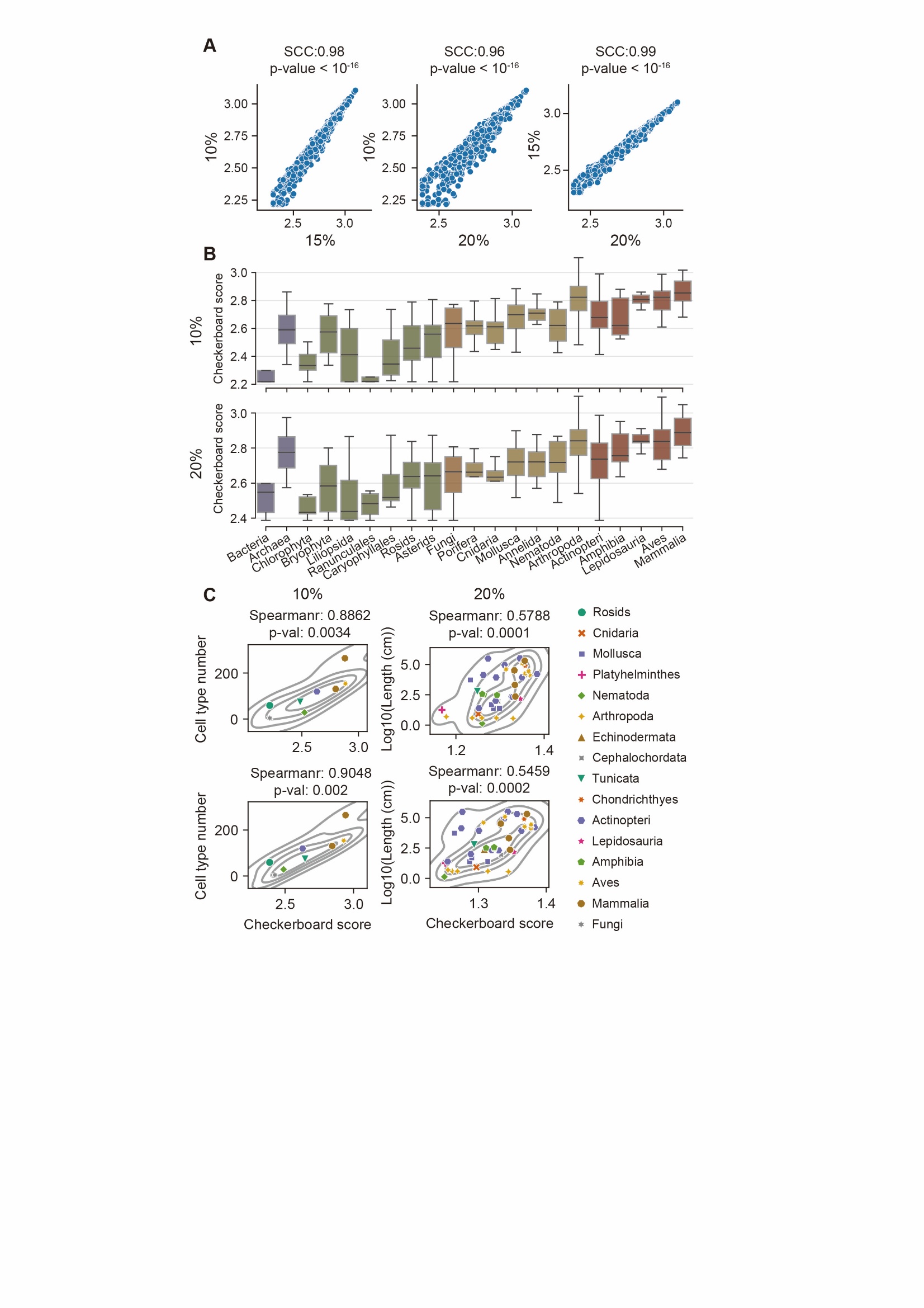

#### Figure S25 Threshold-Independent Robustness of Checkerboard Quantification

A. High consistency across long-distance thresholds of checkerboard quantification. Checkerboard scores exhibit strong correlation between 10%, 15%, and 20% thresholds across 1,025 species: 10% vs. 15% (Spearman correlation coefficient, SCC = 0.98, *p* <10e-16), 10% vs. 20% (SCC: 0.96, *p* <10e-16), and 15% vs. 20% (SCC: 0.99, *p* <10e-16).
B. Consistency of checkboard scores in taxonomic groups. Checkerboard scores for 21 taxonomic groups remain concordant across thresholds. Results of 15% interaction similarity thresholds across 1,025 species are shown in Figure 3.
C. Consistency of correlations between checkboard scores and species complexity.

Correlations of 15% interaction similarity thresholds across 1,025 species are shown in Figure 3. The threshold invariance of checkerboard quantification validates its reliability for cross-species comparisons.

### Supplementary Tables

#### Supplementary Table 1

Hi-C dataset information and processing details.

#### Supplementary Table 2

Intra-global folding scores (GFS) of 1,025 species.

#### Supplementary Table 3

Inter-global folding scores (GFS) of 1,025 species.

#### Supplementary Table 4

Checkerboard scores (CBS) of 1,025 species.

#### Supplementary Table 5

Global folding scores (GFS) and checkerboard scores (CBS) for mature tissues from same species.

#### Supplementary Table 6

Global folding scores (GFS) and significance of Wilcoxon signed-rank test for the main five taxonomic group.

#### Supplementary Table 7

Differences of global folding scores across different taxonomic levels.

#### Supplementary Table 8

Global folding polymorphism scores for 21 taxonomic group.

#### Supplementary Table 9

Global folding scores (GFS) and significance of Wilcoxon signed-rank test for 21 taxonomic group.

#### Supplementary Table 10

Species complexity as proxied by cell type number (CTN) or body length (kg).

#### Supplementary Table 11

Checkerboard scores (CBS) and significance of Wilcoxon signed-rank test for 21 taxonomic group.

#### Supplementary Table 12

CenterFinder modes used for each species.

#### Supplementary Table 13

Manually re-chose center anchors for each species.
